# Probabilistic cell/domain-type assignment of spatial transcriptomics data with SpatialAnno

**DOI:** 10.1101/2023.02.08.527590

**Authors:** Xingjie Shi, Yi Yang, Xiaohui Ma, Yong Zhou, Zhenxing Guo, Chaolong Wang, Jin Liu

**Affiliations:** KLATASDS-MOE, Academy of Statistics and Interdisciplinary Sciences, School of Statistics, East China Normal University; Centre for Quantitative Medicine, Health Services & Systems Research, Duke-NUS Medical School; College for Mathematics and Statistics, South-Central Minzu University; School of Data Science, The Chinese University of Hong Kong, Shenzhen; Department of Epidemiology and Biostatistics, School of Public Health, Tongji Medical College, Huazhong University of Science and Technology

## Abstract

In the analysis of both single-cell RNA sequencing (scRNA-seq) and spatially resolved transcriptomics (SRT) data, classifying cells/spots into cell/domain types is an essential analytic step for many secondary analyses. Most of the existing annotation methods have been developed for scRNA-seq datasets without any consideration of spatial information. Here, we present SpatialAnno, an efficient and accurate annotation method for spatial transcriptomics datasets, with the capability to effectively leverage a large number of non-marker genes as well as “qualitative” information about marker genes without using a reference dataset. Uniquely, SpatialAnno estimates low-dimensional embeddings for a large number of non-marker genes via a factor model while promoting spatial smoothness among neighboring spots via a Potts model. Using both simulated and four real spatial transcriptomics datasets from the 10x Visium, ST, Slide-seqV1/2, and seqFISH platforms, we showcase the method’s improved spatial annotation accuracy, including its robustness to the inclusion of marker genes for irrelevant cell/domain types and to various degrees of marker gene misspecification. SpatialAnno is computationally scalable and applicable to SRT datasets from different platforms. Furthermore, the estimated embeddings for cellular biological effects facilitate many downstream analyses.

## Introduction

With the rapid advancement of spatially resolved transcriptomics (SRT) technologies, it has become feasible to comprehensively characterize the gene expression profiles of tissues while retaining information on their physical locations. Among the already developed SRT methods, *in situ* hybridization (ISH) technologies, such as MERFISH^1^ and seqFISH^2^, provide single-molecule resolution for targeted genes but require prior knowledge of the genes of interest; while *in situ* capturing technologies, such as 10x Visium, Slide-seqV1/2^3^, and Stereo-seq^4^, are unbiased and provide transcriptome-wide expression measurements. Among the *in situ* capturing technologies, there has been a dramatic improvement in spatial resolution, with spot sizes ranging from 55 *μ*m in 10x Visium, 10 *μ*m in Slide-seqV2, to <1 *μ*m in Stereo-seq. These SRT technologies provide an opportunity to study how the spatial organization of gene expression in tissues relates to tissue functions^5^. To characterize the transcriptomic landscape within a spatial context, assigning cell/domain types in relation to tissue location is an essential analytic step that provides comprehensive spatially resolved maps of tissue heterogeneity^6^.

Conventionally, spatial annotation relies on the manual assignment of cell/domain clusters using known marker genes that are readily available from existing studies or databases^7,8^. A general workflow begins with the unsupervised clustering of spots based on their transcriptomic profiles; this is followed by an examination of the differentially expressed genes (DEGs) specific to each cluster; and finally, the DEGs are manually matched with known marker genes to assign cell/domain types to spatial spots. This type of workflow requires sufficient knowledge of the biology and markers of the cell/domain types, but it can be time-consuming, labor-intensive, and less reproducible^6,9^. Moreover, these workflows are sensitive to the choice of clustering methods, presenting challenges in the downstream interpretations^10^. An improved strategy for spatial annotation is to automatically annotate the identified clusters using either reference data or leveraging existing information on the cell/domain types. Performing annotations with reference data has been shown to be successful in the context of single-cell RNA sequencing (scRNA-seq) analysis. For example, scmap performs cell annotation by projecting existing reference data with known cell types onto cells in the study data^11^. However, the success of this type of analysis relies on the availability of reference data that are “similar” to the study data. On the other hand, the availability of data on cell-type-specific maker genes from existing studies or databases, potentially obtained using either low-throughput or high-throughput systems, further necessitates the efficient utilization of marker-gene information in a “qualitative” manner. To this end, a number of methods have been developed for scRNA-seq data without any consideration of spatial information, including SCINA^12^, Garnett^13^, CellAssign^14^, and scSorter^15^.

To efficiently utilize the existing knowledge base on marker genes for cell/domain types, an ideal annotation method for SRT datasets should be capable of leveraging this “qualitative” information on marker genes with data on non-marker genes while incorporating spatial information to promote spatial smoothness in the cell/domain-type annotation. Because the proportion of non-marker genes is much larger than that of marker genes, non-marker genes also harbor substantial amounts of biological information that can be used to separate cell/domain types. Annotation methods capable of leveraging marker with non-marker genes can improve our ability to detect spatial cell/domain clusters ^14,15^. However, the high-dimensional nature of non-marker genes makes the annotation task more challenging and, moreover, requires proper and efficient modeling of this information. Furthermore, for SRT datasets, especially those from tissue sections with laminar structures, e.g., brain regions, a desirable spatial annotation method would additionally be able to leverage spatial information.

To address the challenges presented by spatial annotation, we propose the use of a probabilistic model, SpatialAnno, which performs cell/domain-type assignments for SRT data and has the capability of leveraging non-marker genes to assign cell/domain types via a factor model while accounting for spatial information via a Potts model ^16,17^. To effectively leverage a large number of non-marker genes and overcome the curse of dimensionality, SpatialAnno uniquely models expression levels in a factor model governed by separable cell/domain-type low-dimensional embeddings. As a result, SpatialAnno not only performs spatial cell/domain-type assignments with better accuracy, but also estimates cell/domain-type-aware embeddings that can facilitate downstream analyses. We illustrate the benefits of SpatialAnno through extensive simulations and analyses of a diverse range of example datasets collated using different spatial transcriptomics technologies. To show the improved spatial annotation accuracy, we applied SpatialAnno to analyze a 10x Visium datasets for 12 human dorsolateral prefrontal cortex (DLPFC) samples. To illustrate the effectiveness of SpatialAnno in leveraging non-marker genes, we analyzed a mouse olfactory bulb (OB) dataset generated using the ST technology. Using Slide-seqV1/2 datasets for the mouse hippocampus, we demonstrated that SpatialAnno can correctly identify cell-type distribution at near-cell resolution. The utility of SpatialAnno to estimate low-dimensional embeddings is demonstrated by a seqFISH dataset for the mouse embryo.

## Results

### Overview of SpatialAnno

Similarly to other methods that assign known cell/domain types to cells using information about marker genes, SpatialAnno takes as input normalized gene expression matrix, spatial location information, and a list of marker genes for known cell/domain types (Fig. 1a). SpatialAnno automatically performs cell/domain-type assignments while providing low-dimensional embeddings for all spatial spots. Based on the latent cell/domain type for each spot, SpatialAnno builds a “semi-supervised” Gaussian mixture model to modulate the over-expression of marker genes and a hierarchical factor model to relate non-marker gene expression to the cell/domain separable latent embeddings while accounting for the spatial smoothness of the cell/domain types with a Potts model (Fig. 1b). Uniquely, SpatialAnno, via the factor model, allows for the assignment of cell/domain types that leverage a large number of non-marker genes, and, via the Potts model, is more likely to assign the same cell/domain type to neighboring spots, promoting spatial smoothness in the cell/domain types. Notably, with expression data for both marker and non-marker genes, SpatialAnno simultaneously assigns each spot known cell/domain types while obtaining low-dimensional embeddings for each spot, which can facilitate other downstream analyses. Similarly to other methods, SpatialAnno automatically labels spatial spots that do not belong to any known cell/domain type as “unknown”, preventing incorrect assignment when novel cell/domain types are present.

**Figure 1:**
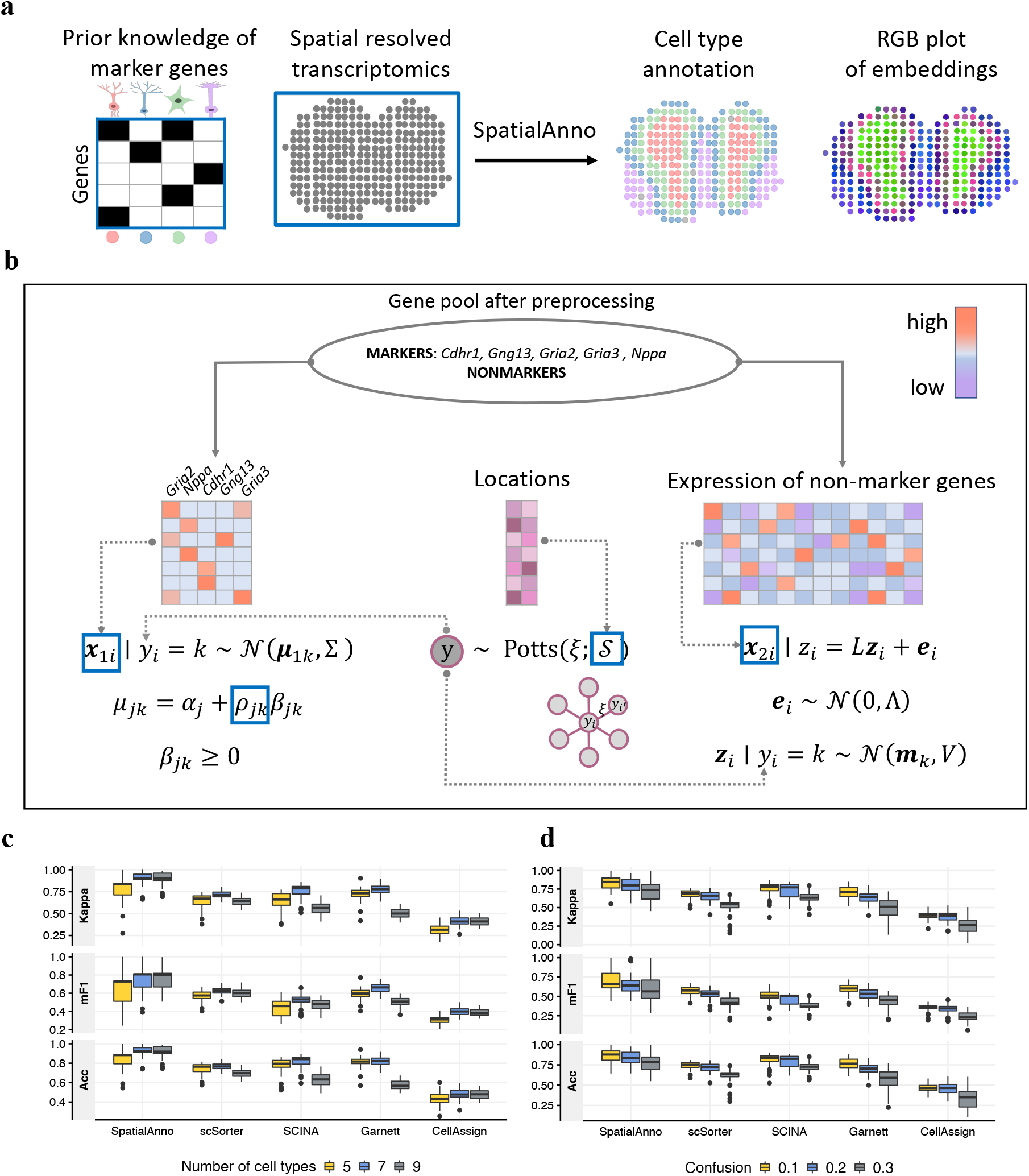
Schematic overview of SpatialAnno and its performance in simulation studies. **a** SpatialAnno employs spatial transcriptomics data along with a known marker-gene list in its analysis. With these two datasets as input, SpatialAnno performs spatial annotation via a probabilistic model that combines both marker and non-marker gene expression data, and produces both domain/cell-type assignments and low-dimensional embeddings for all spatial locations as output. **b** Overview of the SpatialAnno probabilistic model. Latent cell/domain types (shown in the grey circle) and observed data (shown in the blue boxes) are shown along with the distributional assumptions. **c** Kappa, mF1, and ACC of SpatialAnno, scSorter, SCINA, Garnett, and CellAssign for simulation data from seven cortical layers; different numbers of cell/domain types are provided as a list of marker genes. **d** Kappa, mF1 and ACC of SpatialAnno, scSorter, SCINA, CellAssign and Garnett for simulation data from seven cortical layers; different proportions of marker genes are erroneously specified.

### Validation using simulated data

We conducted simulations to evaluate the performance of SpatialAnno and compared the results with those of non-spatial annotation methods commonly applied to scRNA-seq data: SCINA, Garnett, CellAssign, and scSorter (see Methods). The simulation details are provided in the Methods section. Briefly, we simulated gene expression counts using a splatter model ^18^ for seven cortical layers using labels from the DLPFC data. Then, we selected five marker genes for each layer based on the log-fold change in expression (see Methods). In total, we obtained 35 marker genes and 2000 non-marker genes for 3639 spots from seven layers. For each simulated SRT dataset, we applied SpatialAnno and the four other methods to perform spatial domain annotation. We used Cohen’s Kappa, mean F1 (mF1) score, and classification accuracy (ACC) (see Methods) to quantify the concordance between the detected spatial domains and the seven labeled cortical layers^11,14^. We performed 50 replicate simulations for each setting.

When the correct number of layers was specified, SpatialAnno (Kappa=0.903, mF1=0.807, and ACC = 0.922) outperformed all other methods in terms of annotation accuracy (Fig. 1c; number of cell/domain types = 7). After varying the number of cell/domain types with marker genes, the SpatialAnno annotation still outperformed all other methods (Fig. 1c; number of cell/domain types = 5 or 9). Unsurprisingly, SpatialAnno performed worse when there were five cell/domain types with marker genes (Kappa=0.839, mF1=0.729, and ACC = 0.883) than seven or nine (Kappa=0.900, mF1=0.803, and ACC = 0.918). The latter two cases (seven and nine cell/domain types) led to comparable annotation performances for SpatialAnno and CellAssign. In contrast, annotation performance decreased for the other methods when we included marker genes for irrelevant cell/domain types. We examined the robustness of SpatialAnno when there were various degrees of marker gene misspecification (Fig. 1d). As the proportion of misspecified marker genes increased, the annotation performance decreased for all methods, but SpatialAnno still outperformed all other methods in terms of annotation accuracy (Kappa, mF1, and ACC).

Next, we examined the effectiveness of SpatialAnno, which leverages various amounts of non-marker information compared with the scSorter and Garnett methods, also capable of leveraging non-marker genes (Supplementary Fig. 1a). As the number of non-marker genes increased from 60 to 2000, SpatialAnno showed 10.3%, 21.9% and 8.1% improvements in annotation accuracy for Kappa, mF1 and ACC, respectively, while the annotation accuracies of scSorter and Garnett were almost unchanged, with the changes being −0.6% and −0.6% for Kappa, 1.7% and −0.1% for mF1, and −0.1% and −0.7% for ACC, respectively. These results suggest that SpatialAnno can effectively leverage various numbers of non-marker genes.

In addition to the spatial spots being accurately annotated, the low-dimensional embedding of non-marker genes from SpatialAnno was cell/domain-type informative. Clustering performance using low-dimensional embeddings with either marker genes or non-marker genes, or a combination of the two, with a comparable adjusted rand index (ARI) between marker and non-marker genes, is shown in Supplementary Fig. 1b & c. Not surprisingly, combining both embeddings for marker and non-marker genes led to improved ARIs in all scenarios, demonstrating the benefits of borrowing information from non-marker genes when annotating cell/domain types. In addition, the Pearson’s correlation coefficients for the relationship between the observed expression and the estimated labels, given the embeddings from SpatialAnno, were much smaller than those for the principal component analysis (PCA), but comparable to those for the DR-SC^19^ (Supplementary Fig. 1d & e). These results suggest SpatialAnno embeddings can capture cell/domain-type-relevant information for each spot, thus facilitating the downstream analysis.

Finally, we evaluated the computational efficiency of all methods for different numbers of cell/domain types, as shown in Supplementary Fig. 1f. SpatialAnno was computationally efficient and comparable in efficiency to SCINA and scSorter, and all three were faster than Garnett and CellAssign.

### SpatialAnno improves annotations of known layers in human dorso-lateral prefrontal cortex

We applied SpatialAnno and the four methods to the analysis of human dorsolateral prefrontal cortex (DLPFC) 10x Visium data^20^. In this dataset, there were 12 tissue sections from three adult donors with a median depth of 291 million reads for each sample, a median of 3844 spatial spots per section, and a mean of 33,538 genes per spot (Supplementary Table 1). Each tissue section was manually annotated to a DLPFC layer and white matter (WM) based on the cytoarchitecture^20^. Taking sample ID151507 as a reference, we constructed a marker-gene list that contained five marker genes for each of the seven layers (see Methods).

Taking manual annotations as ground truth, we first evaluated the performance of spatial annotation using Kappa, mF1, and ACC for each of the 12 tissue sections (Fig. 2a). SpatialAnno annotated spatial domains more accurately (median Kappa=0.524, median mF1=0.494, and median ACC=0.628) than scSorter (median Kappa=0.381, median mF1=0.366, and median ACC=0.489), SCINA (median Kappa=0.209, median mF1=0.337, and median ACC=0.307), Garnett (median Kappa=0.24, median mF1=0.32, and median ACC=0.339), and CellAssign (median Kappa=0.253, median mF1=0.29 and median ACC=0.326). The heatmap of the spatial assignments from SpatialAnno and the other methods and the manual annotations for sample ID151673 are shown in Fig. 2b. SpatialAnno achieved the best annotation accuracy (Kappa=0.634, mF1=0.619, and ACC=0.685), while the annotations from scSorter, SCINA, and CellAssign were only accurate for the WM, and Garnett completely failed to assign the WM region. Notably, the domains identified in SpatialAnno were spatially smooth, continuous, and well matched with the elevated expression levels of marker genes for each layer (Fig. 2c and Supplementary Fig. 2-13), such as *Pcp4* and *Mobp* that are marker genes for layer 5 and WM, respectively^20,21^.

**Figure 2:**
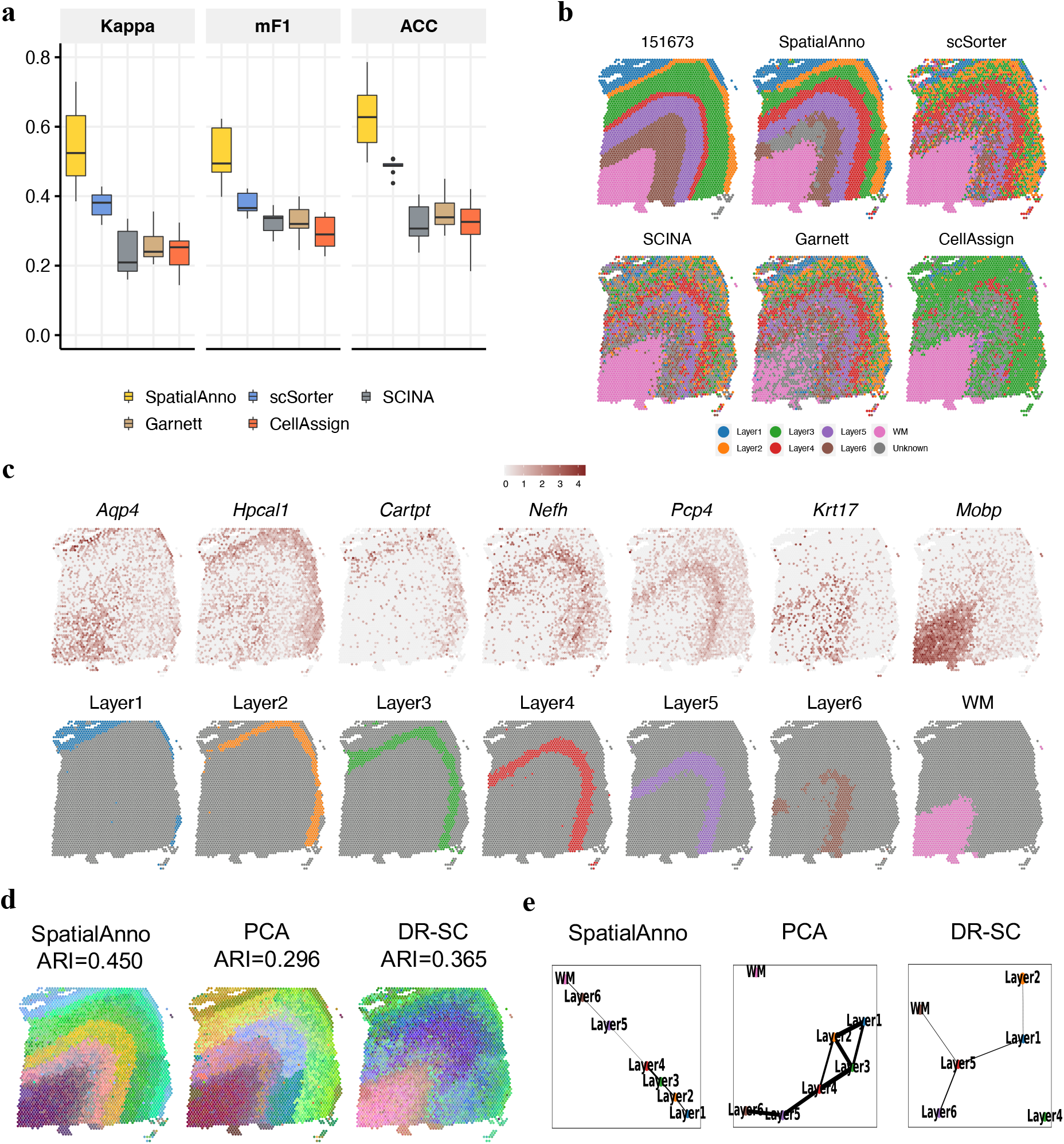
Spatial domain annotation in the DLPFC 10x Visium dataset. **a** Boxplots of Kappa, mF1, and ACC showing the accuracy of different methods for domain annotation across 12 tissue sections. **b** Spatial domain annotation in tissue sample ID151673 for ground truth, SpatialAnno, scSorter, SCINA, Garnett and CellAssign. **c** Top, expression levels of corresponding layer-specific marker genes. Bottom, annotations by SpatialAnno are shown on each spot. **d** RGB plots for low-dimensional embedding inferred by SpatialAnno, PCA, and DR-SC. As end-to-end annotation approaches, scSorter, SCINA, Garnett, and CellAssign cannot be utilized to extract low-dimensional embeddings. **e** PAGA graphs generated by SpatialAnno, PCA, and DR-SC embeddings for DLPFC Section ID151673.

To evaluate the robustness of SpatialAnno, we obtained marker genes from the other DLPFC tissue section that contained seven layers and performed spatial annotation for the remainder of the 11 tissue sections (see Methods). Using the top 5/10/15 DEGs as marker genes for each layer, SpatialAnno achieved the best annotation accuracy according to Kappa, mF1, and ACC. The annotation accuracies of all other methods for the other tissue sections were slight worse than for those when sample ID151507 was used as a reference (Supplementary Fig. 14a), which is consistent with the simulations involving the misspecification of marker genes (Fig. 1d). This suggests that annotation accuracy can be impaired when inaccurate marker genes are used. However, this difference became negligible when the number of marker genes for each layer was 15. Furthermore, we examined the robustness of SpatialAnno using marker genes for irrelevant cell types, those not present in the studied SRT dataset. For samples ID151669-151672 from Donor 2, which only contained five cortical layers, we applied SpatialAnno and other methods using marker genes for the seven layers. As shown in Supplementary Fig. 14b, SpatialAnno achieved the best annotation performance for these samples.

Uniquely amongst the methods, SpatialAnno’s estimated embeddings were highly informative for the DLPFC layers in the 12 sections. The clustering accuracies, determined using the ARI for embeddings from marker, non-marker, and a combination of the two, respectively, were shown in Supplementary Fig. 14c, with the largest ARI value for embeddings from a combination of the two. Clearly, embeddings from non-marker genes harbored substantial amount of information about spatial domains, even more than the marker genes. When using a combination of marker and non-marker genes, the embeddings led to improved clustering performance, suggesting that annotation based on both marker and non-marker genes improved the annotation accuracy. Red/green/blue (RGB) plots using three tSNE components for the embeddings in sample ID151673 estimated by SpatialAnno revealed a more clear laminar structure for DLPFC than those by PCA or DR-SC (Fig. 2d). Such stronger structure predictivity from SpatialAnno is numerically supported by its higher ARI (0.450) compared to PCA (ARI=0.296) and DR-SC (ARI=0.365). Moreover, an estimated PAGA graph^22^ using SpatialAnno embeddings demonstrated the almost linear development trajectory from WM to layer 1, while the PAGA graphs using both PCA and DR-SC embeddings were less clearly delineated (Fig. 2e and Supplementary Fig. 2-13).

### SpatialAnno correctly identifies cells in mouse olfactory bulb

To quantitatively demonstrate the performance of SpatialAnno compared with SCINA, scSorter, CellAssign, and Garnett in domain-type annotation, we analyzed one mouse OB data generated using ST technology. This dataset represented 12 tissue sections with a median of 16,024 gene expression measurements among a median of 266 spots (Supplementary Table 2).

Taking the four anatomic layers manually annotated based on H&E staining as ground truth (Fig. 3a), we first evaluated the performance of the spatial annotation using Kappa, mF1, and ACC for section 12 (Fig. 3b). SpatialAnno annotated spatial domains more accurately (Kappa=0.739, mF1 = 0.812, and ACC=0.800) than scSorter (Kappa=0.608, mF1=0.718, and ACC=0.696), SCINA (Kappa=0.598, mF1=0.670, and ACC=0.689), CellAssign (Kappa=0.395, mF1=0.607, and ACC=0.707), and Garnett (Kappa=0.552, mF1=0.686, and ACC=0.646). We examined the robustness of SpatialAnno by including marker genes for two irrelevant cell types (endothelial and mural cells) that were not present in this section, and SpatialAnno achieved the best annotation performance (Supplementary Fig. 15a). To illustrate the effectiveness of leveraging non-marker information, we evaluated the performance of the spatial annotation by SpatialAnno, scSorter, and Garnett with 30, 300, or 3000 non-marker genes, as only these three methods are able to leverage non-marker gene information. SpatialAnno achieved higher annotation accuracy when more non-marker genes were used, while the difference in performance between 300 and 3000 non-marker genes was minimal for SpatialAnno (Supplementary Fig. 15b). In contrast, scSorter and Garnett performed similarly with 30 or 300 non-marker genes, but their performance deteriorated when 3000 non-marker genes were applied.

**Figure 3:**
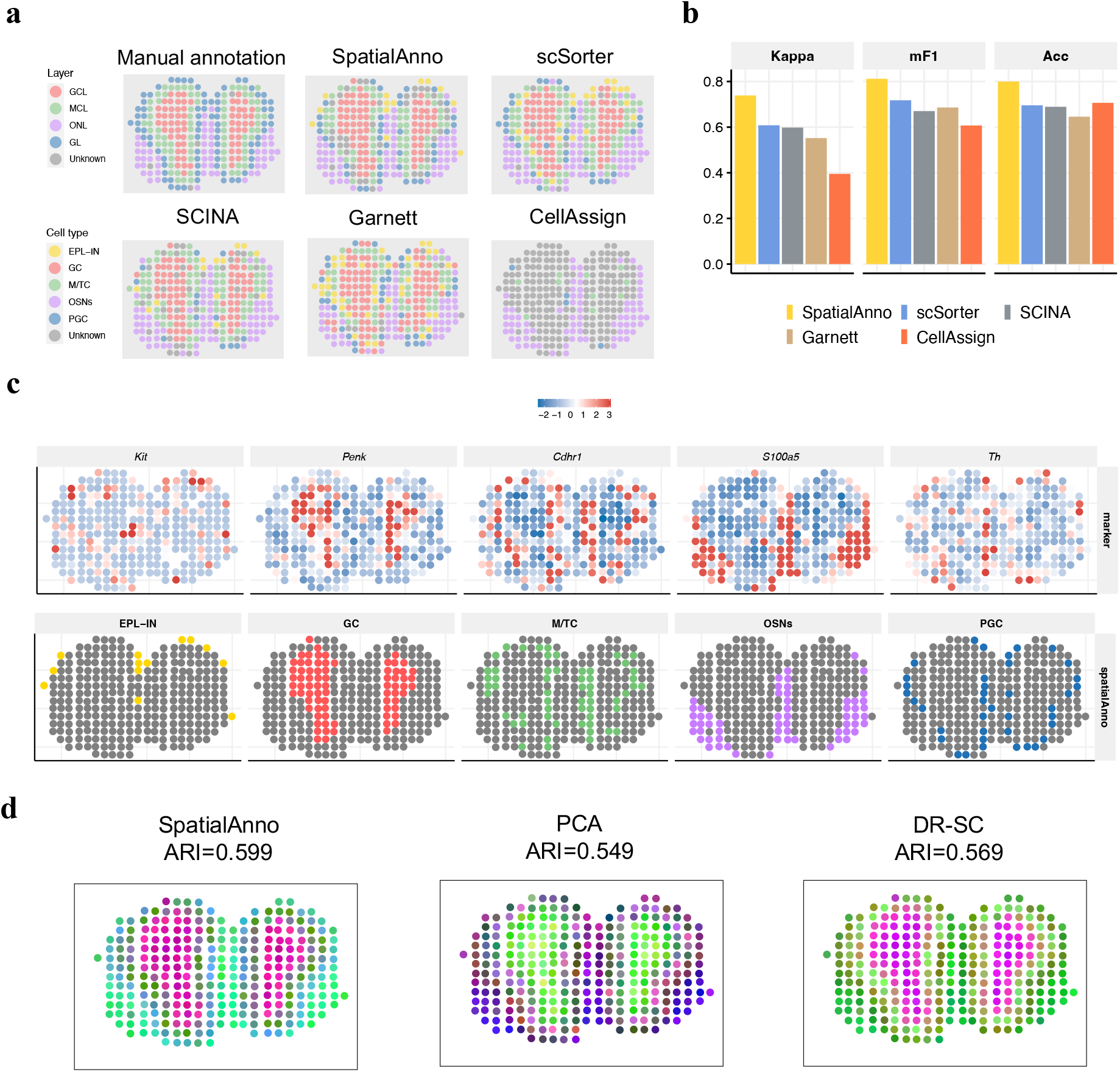
Spatial domain annotation in the mouse olfactory bulb dataset. **a** Spatial domain annotations for ground truth, SpatialAnno, scSorter, SCINA, Garnett, and CellAssign. **b** Bar plots of Kappa, mF1 and ACC showing the domain-type annotation accuracy of different methods. **c** Top, expression levels of corresponding cell-type-specific marker genes. Bottom, annotations by SpatialAnno are shown on each spot. **d**, RGB plots of low-dimensional embeddings inferred by SpatialAnno, PCA, and DR-SC. As end-to-end annotation approaches, scSorter, SCINA, Garnett, and CellAssign cannot be utilized to extract low-dimensional embeddings.

SpatialAnno recovered the laminar structure of the mouse OB across 12 sections (Supplementary Fig. 16). The mouse OB has a multi-layered cellular architecture in the order, from the inner to outer layer, of granule cell layer (GCL), mitral cell layer (MCL), glomerular layer (GL), the nerve layer (ONL). Detailed assignments by SpatialAnno and the other four methods for section 12 are shown in Fig. 3a. The cell types annotated by SpatialAnno accurately represented this laminar structure, while CellAssign incorrectly assigned “unknown” cells to regions belonging to GCL, MCL, and GL. Moreover, the annotation patterns of Garnett were rather chaotic, while scSorter and SCINA failed to distinguish periglomerular cells (PGC) in the GL.

We further examined the expressions of marker genes specific to each layer, including *Kit* for external plexiform layer interneuron (EPL-IN)^23^, *Penk* for granule cells (GC)^24^, *Cdhr1* for mitral and tufted cells (M/TC)^25^, *S100a5* for olfactory sensory neurons (OSN)^26^, and *Th* for PGC^27^ (Fig. 3c). Although the three methods provided similar assignments for GC, M/TC, OSN, and PGC, their assignments for EPL-IN were quite different. EPL-IN are located adjacent to GL in the external plexiform layer comprised of PGC^23^. SpatialAnno assigned spots near PGC to EPL-IN; however, scSorter and Garnett did not (Supplementary Fig. 17). As the ground truth for the EPL-IN locations was unknown, we manually combined the inferred EPL-IN with the adjacent layers in different ways: (1) by combining the inferred EPL-IN and PGC and (2) by combining the inferred EPL-IN, M/TC, and PGC. SpatialAnno still achieved the best annotation accuracy (Supplementary Fig. 15c & d).

Another key benefit of SpatialAnno is its ability to extract low-dimensional embeddings relevant to different cell types from the high-dimensional non-marker genes, which is useful for many downstream analyses. We summarized the low-dimensional embeddings inferred by SpatialAnno (Supplementary Fig. 15e), PCA, and DR-SC into three-dimensional tSNE components and visualized the resulting components in the RGB plot. The RGB plot (Fig. 3d) shows the multi-layered architecture of the mouse OB, with neighboring spots sharing more similar colors to those farther away. To compare the predictive powers of these low-dimensional embeddings for the four anatomic layers annotated based on H&E staining, we applied the Louvain community detection algorithm to spot clustering using the *Seurat* R package. The clusters identified by SpatialAnno depicted the multi-layered structures more accurately (ARI = 0.599) than those of PCA (ARI = 0.549) or DR-SC (ARI = 0.569).

### SpatialAnno reveals cell-type distribution in mouse hippocampus with SRT data at near-cell resolution

To show the cell-type distribution in the mouse hippocampus, we applied SpatialAnno and the other methods to the analysis of a mouse hippocampus dataset generated using Slide-seqV2, which quantifies transcriptome-wide expression levels at near-cellular resolution with 10-*μ*m barcoded beads^3^. This dataset contains expressions for 23,264 genes over 53,208 spatial locations (Supplementary Table 3). As shown in the Allen Reference Atlas (Fig. 4a), the primary regions in the mouse hippocampus were composed of the cornu ammonis (CA1-3) and dentate gyrus (DG).

**Figure 4:**
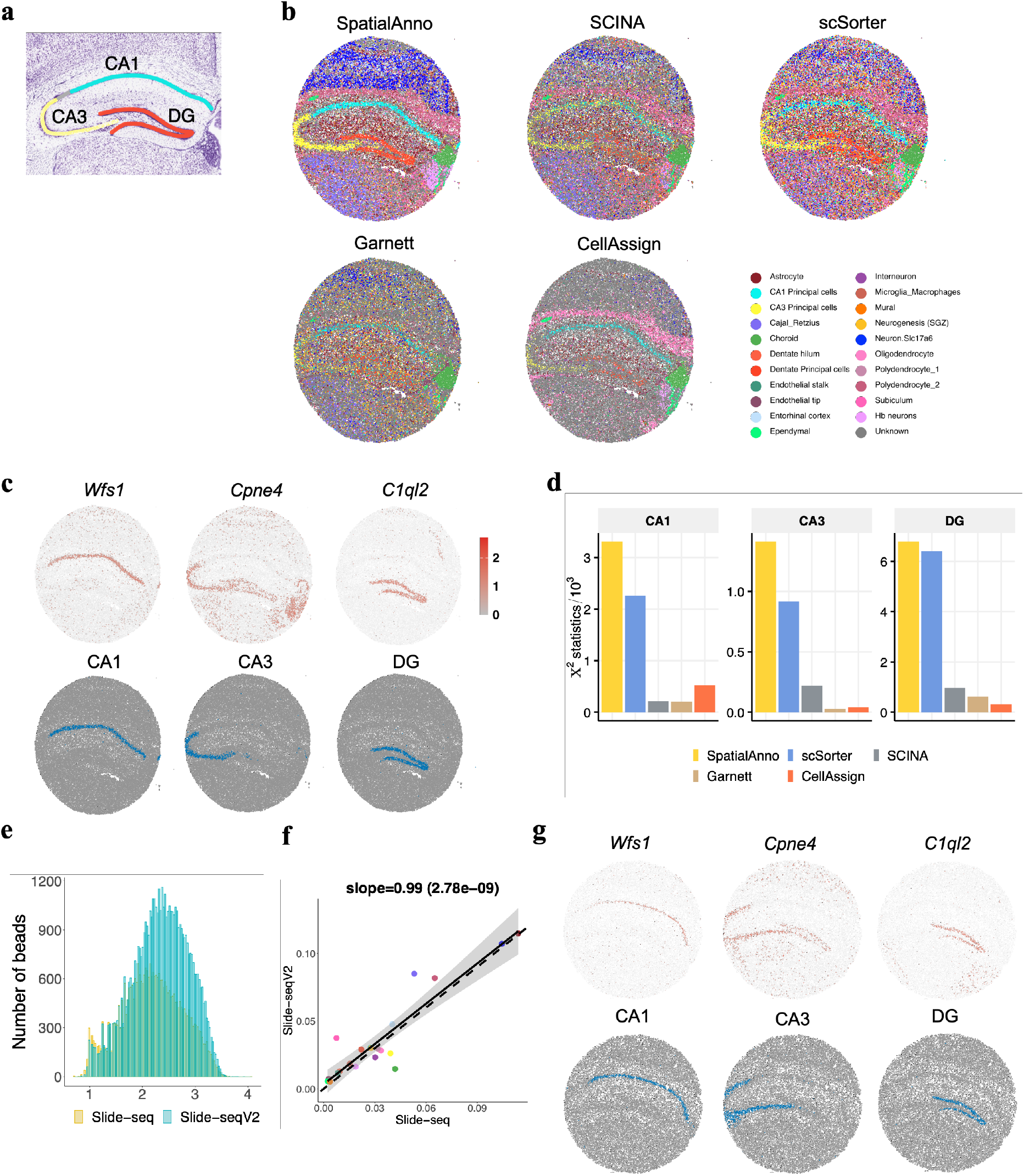
Spatial cell-type annotation of the mouse hippocampus dataset. **a** Annotation of hippocampus structures from the Allen Reference Atlas of an adult mouse brain. **b** Spatial annotation of the Slide-seqV2 hippocampus section by SpatialAnno, scSorter, SCINA, Garnett, and CellAssign. **c** Top, expression levels of corresponding cell-type-specific marker genes. Bottom, annotations by SpatialAnno of the Slide-seqV2 hippocampus section are shown on each spot. The examined cell types were CA1 cells, CA3 cells and dentate cells. **d** Results of Pearson’s chi-squared test of correlation between expression patterns of marker genes and the three hippocampal subfields identified by different methods. **e** Total UMIs per bead for Slide-seq (yellow, *n* = 34, 199 spots) versus Slide-seqV2 (blue, *n* = 53, 208 spots) in the mouse hippocampus sections. **f** Scatter plot of cell-type proportions identified by SpatialAnno in Slide-seq and Slide-seqV2 datasets. **g** Top, expression levels of corresponding cell type specific marker genes. Bottom, annotation by SpatialAnno of the Slide-seq hippocampus section is shown on each spot.

SpatialAnno clearly identified a “cord-like” structure as well as an “arrow-like” structure in the hippocampal subfields in CA1, CA3, and DG (Fig. 4b), which is consistent with the annotation of hippocampus structures in the Allen Reference Atlas (Fig. 4a). In contrast to SpatialAnno, the other methods SCINA, Garnett, and CellAssign showed blurred/incorrect localizations for the primary hippocampal subfields in CA3 and DG and were unable to reveal the main structures of the mouse hippocampus (Fig. 4b and Supplementary Fig. 18-20). The hippocampal subfields identified by scSorter were surrounded by a blurry border, with many different cell types allocated to the same region. Additionally, all the methods except SpatialAnno failed to accurately allocate the habenula (Hb)neurons, which should reside left to and below the choroid plexus. Careful examination of marker genes further demonstrated the superior accuracy of SpatialAnno (Fig. 4c), i.e., *Wfs1*, *Cpne4*, and *C1ql2* for CA1, CA3, and DG, respectively.

We quantified the annotation performance of the different methods by examining the correlations between the expression patterns of the marker genes and the three hippocampal subfields identified by the different methods. Pearson’s chi-squared test demonstrated a substantial improvement in the magnitude of associations provided by SpatialAnno (Fig. 4d). The RGB plot for SpatialAnno displayed clear regional segregation of the hippocampus (Supplementary Fig. 21a). Specifically, compared with the RGB plots for PCA and DR-SC, the plot for SpatialAnno clearly depicted the Hb region.

Finally, we validated the cell-type distributions identified for an independent slide from the mouse hippocampus profiled using Slide-seq. As with the initial version of Slide-seqV2, the transcript detection sensitivity of Slide-seq is relatively low (Fig. 4e). By applying SpatialAnno to this validation dataset, we showed the consistency of the cell-type distributions between the two slides, as illustrated in Fig. 4f. SpatialAnno successfully identified the hippocampal subfields in this Slide-seq data (Supplementary Fig. 21b-d and Supplementary Fig. 22-24). The annotated regions for CA1, CA3, and DG with their marker gene expressions are shown in Fig. 4g.

### Embeddings estimated by SpatialAnno lead to biologically relevant trajectories in mouse embryo

We further applied SpatialAnno and the other methods to the analysis of a dataset obtained from three mouse embryo sections collated at the 8-12 somite stage using seqFISH ^2^, which has the capability of probing the expression of a targeted gene set at the single-cell resolution ^2^. Each of the three mouse embryo sections contained expression level measurements for 351 genes, chosen to recover the cell-type identities at these developmental stages, from around 20,000 cells, as well as their physical locations (Supplementary Table 4). After selecting 168 marker genes for 21 cell types (see Methods), 183 non-marker genes remained for annotation analysis.

The original study provided manual annotations for the cells based on their nearest neighbors in the Gastrulation atlas^28^. For each method, we summarized the annotation accuracy using both Kappa, mF1 and ACC for each embryo section (Fig. 5a and Supplementary Fig. 25). SpatialAnno achieved the highest Kappa, mF1 and ACC in two out of the three sections and was only surpassed by CellAssign for the second embryo section. For Embryo 1, the annotations of different methods are shown in Fig. 5b. Clearly, cell-type distributions identified by SpatialAnno were well matched with the expression of their corresponding marker genes (Fig. 5c).

**Figure 5:**
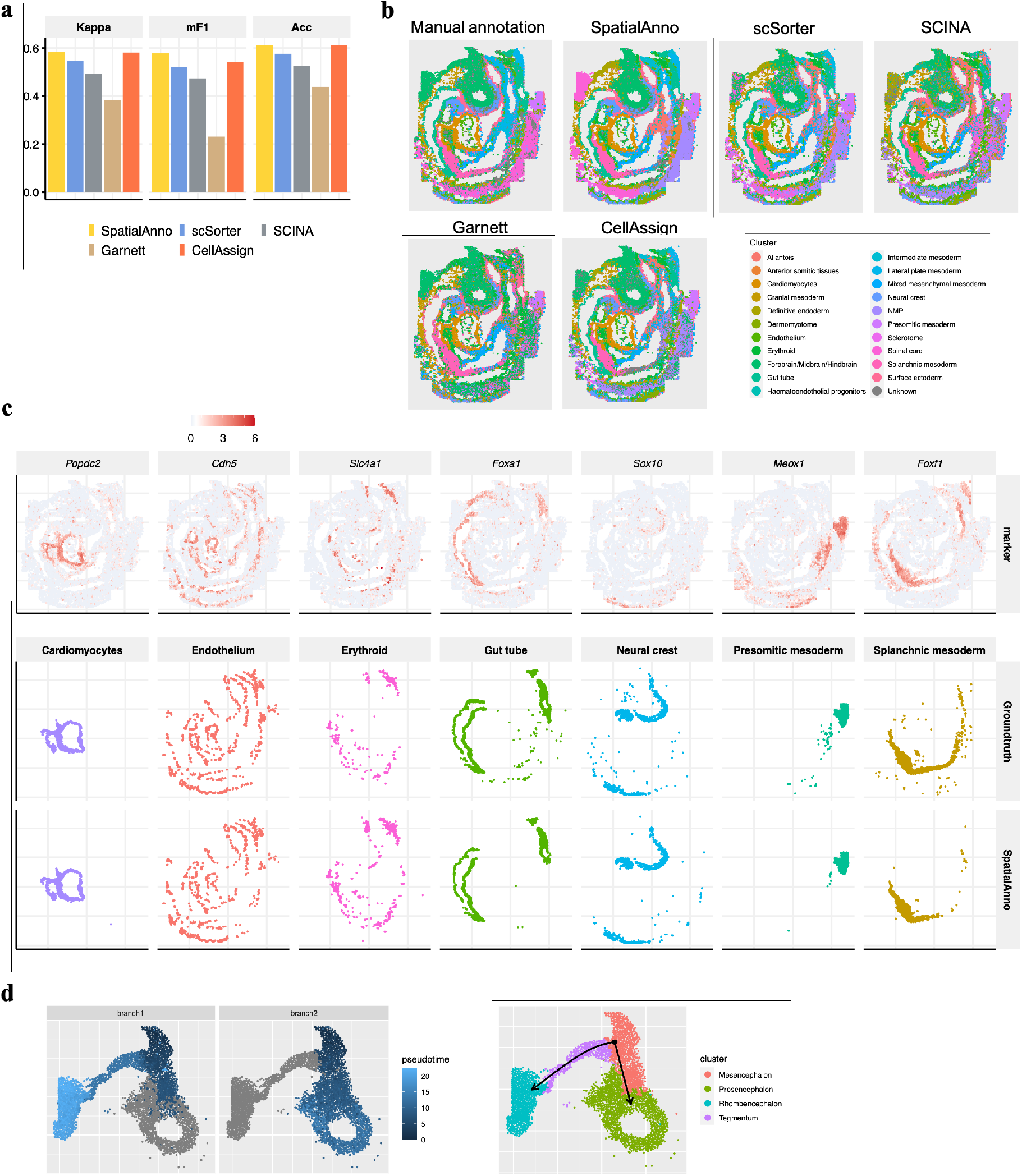
Spatial cell-type annotation of the mouse embryo dataset. **a** Bar plots of Kappa, mF1 and ACC showing the cell-type annotation accuracy of different methods. **b** Spatial annotations for ground truth, SpatialAnno, scSorter, SCINA, Garnett, and CellAssign. **c** Top, expression levels of corresponding cell-type-specific marker genes. Bottom, annotations of ground truth and SpatialAnno are shown on each spot. **d**, Left: latent time trajectory generated by slingshot on low dimensional embeddings of SpatialAnno. Right: clustering of the forebrain/midbrain/hindbrain cells into four spatially distinct clusters representing different regions of the developing brain.

For the embeddings uniquely estimated by SpatialAnno, we performed trajectory inference on the brain cells to investigate the spatiotemporal development of the mouse brain and detected two linear trajectories (Fig. 5d). We observed the lowest pseudotime values in the mesencephalon, which diffused smoothly towards the tegmentum followed by the rhombencephalon in one branch, and towards the prosencephalon in another branch (Fig. 5d). More importantly, the diffusion patterns were spatially continuous and smooth. The detected trajectories delineated the spatial trajectories of mouse brain development, which are in agreement with the findings of recent studies^2,29^. In contrast, the trajectories identified using embeddings from either PCA or DR-SC lacked spatial continuity (Supplementary Fig. 26a & b). We further examined genes associated with the inferred pseudotime, and a heatmap of the expression levels of the top 20 significant genes suggested there were interesting expression patterns over pseudotime (Supplementary Fig. 26c). A mesencephalon and prosencephalon maker gene, *Otx2* ^30,31^, showed higher expression levels in the early stage of development, while at a later stage, its expression levels were substantially suppressed (Supplementary Fig. 26d). In contrast, the expression levels of a gene enriched in the rhombencephalon, *Sfrp1* ^32^, changed from low to high (Supplementary Fig. 26d). These results concur with the formation of the midbrain-hindbrain boundary ^33,34^, and this is supported by the observation that these two genes could be used to identify the precise boundary between the mesencephalon and rhombencephalon (Supplementary Fig. 26e).

## Discussion

SpatialAnno takes, as input, the normalized gene expression matrix, the physical location of each spot, and a list of marker genes for known cell/domain types. The output of SpatialAnno comprises the estimated posterior probability of each spot belonging to each cell/domain type and the low-dimensional embeddings of each spot for non-marker genes. To efficiently capitalize on both marker and non-marker genes, SpatialAnno uniquely models the expression levels of non-marker genes via a factor model governed by cell/domain-type separable low-dimensional embeddings and simultaneously promotes spatial smoothness via a Potts model. As a result, SpatialAnno provides improved spatial cell/domain-type assignments, and its estimated low-dimensional embeddings are cell-type-relevant and can facilitate downstream analyses such as trajectory inference. SpatialAnno is computationally efficient, easily scalable to spatially resolved transcriptomics with tens of thousands of spatial locations and thousands of genes (Supplementary Table 5). With simulation studies, we demonstrated that SpatialAnno presents improved spatial annotation accuracy with either correct, under- or over-specification of the number of cell/domain types, robustness to the marker gene misspecification, and efficient leveraging of non-marker genes compared with other annotation methods.

We examined the SRT data generated using different platforms, such as 10x Visium, ST, Slide-seqV1/2, and seqFISH, with various spatial resolutions. Using both DLPFC 10x Visium datasets and mouse OB ST datasets with manual annotations, we demonstrated the improved annotation accuracy of SpatialAnno with the capability of recovering laminar structures, while the identified PAGA graph using embeddings in SpatialAnno recovers an almost linear trajectory from WM to layer 1. In DLPFC datasets, the domains identified were well matched with the elevated expression for marker genes, such as *Pcp4* and *Mobp* that are marker genes for layer 5 and WM, respectively ^20,21^, whereas *Pcp4* encodes Purkinje cell protein 4 and *Mobp* encodes the myelin-associated oligodendrocyte basic protein. Using mouse hippocampus Slide-seqV1/2 datasets, we demonstrated that SpatialAnno can successfully detect the primary hippocampal subfields for CA1, CA3, and DG, with almost a perfect correlation between cell-type proportions in both datasets and the elevated expression levels for *Wfs1*, *Cpne4*, and *C1ql2* are well matched with CA1, CA3, and DG regions identified by SpatialAnno, respectively. *Wfs1* showed differential expression in hippocampal field CA1 and has been reported to be highly expressed in the CA1 region^35^. *Cpne4*, a known marker gene for hippocampal subfield CA3, was highly expressed in a region identified as CA3^36^. In addition, *C1ql2*, a marker gene for dentate principal cells, was expressed in a region identified as DG^37^. When applied to mouse embryo seqFISH datasets, SpatialAnno not only provided improved annotation accuracy, but uniquely estimated cell-type-aware embeddings leading to the identification of two trajectories in brain regions, originating in mesencephalon towards the rhombencephalon and prosencephalon, respectively. Moreover, cell-type distributions identified by SpatialAnno were well matched with the expression of their corresponding marker genes. For example, *Popdc2*, a cardiomyocyte marker, was expressed in the developing heart tube ^38^. *Foxa1*, a gut endoderm marker, showed the highest expression levels in the developing gut tube along the anterior–posterior axis of the embryo^39^. In addition, *Foxf1*, a mesoderm marker that encodes a forkhead transcription factor expressed in the splanchnic mesenchyme surrounding the gut, was highly expressed at the identified splanchnic mesoderm^40^.

SpatialAnno paves the way for future spatial annotation analyses in multiple scenarios. For example, a similar strategy can be applied to the problem of cell-type assignment in other spatial omics data, such as spatial resolved single-cell chromatin accessibility data ^41^ and spatial proteomics^42^. To establish a complete spatial atlas of organism architecture, a critical bottleneck is to perform an automatic cell-type assignment with both considerations of molecular features with/without prior knowledge as well as their spatial organization, SpatialAnno can substantially reduce both the irreproducibility and human effort in the processes of manual cell/domain-type assignment^42^.

The benefits of SpatialAnno come with some caveats that may require further exploration. First, SpatialAnno is applicable for spatial annotation in a single tissue slide. With multiple tissue slides available, methods that are capable of integrating multiple SRT datasets for cell/domain-type annotation are sincerely needed ^43^. Second, SpatialAnno was designed to perform annotation analysis of data with a single modality. However, incorporating multi-modal data with data of other modalities can further improve annotation accuracy. Third, many of the early SRT technologies do not have a single-cell resolution, and SpatialAnno is only able to assign domains with prior knowledge of each spot for those datasets. Cell-type annotation for this type of dataset further requires simultaneous deconvolution with spatial cellular annotation.

## Methods

### SpatialAnno method overview

#### Probabilistic models for marker and non-marker gene expression

We herein present an overview of SpatialAnno, with its inference details provided in the Supplementary Notes. SpatialAnno requires both spatial transcriptomics data and a list of gene names for known cell/domain-type markers. The marker-gene list can be obtained from either the available publications, databases, or DEGs in scRNA-seq data (see Methods). In the SpatialAnno model, we denote *X* as the spot-by-gene expression matrix on *n* spatial locations. These locations have known spatial coordinates and unknown labels *y_i_*, *i* = 1*,…, n*. We can separate genes into a group of *m* marker genes and a group of *p* non-marker genes, denoted as ***x***_1*i*_ = (*x_i_*_1_*,…, x_im_*)^⊤^ and ***x***_2*i*_ = (*x_i,m_*_+1_*,…, x_i,m_*_+*p*_)^⊤^, respectively. Suppose prior knowledge of marker genes for *K* cell/domain types is encoded as an indicator matrix *ρ* of dimension *m* × *K*, with *ρ_jk_* = 1 if gene *j* is a maker for cell/domain type *k* and 0 otherwise. Following ^44–46^, we assume that the expression measurements have already been normalized through variance stabilizing transformation and further centered for each gene to have zero mean (see Methods).

SpatialAnno models the centered normalized expression vector, ***x***_1*i*_, for marker genes in cell *i*, and latent label, *y_i_*, as

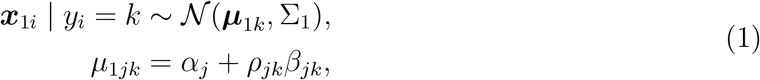

with the constraint that *β_jk_* ≥ 0. Here, *α_j_* is the base expression level for gene *j* in the marker group. The intuition is that if gene *j* is a marker for cell/domain type *k*, then we expect the expression of *j* to be higher in these cell/domain types^14^ with an increased magnitude *β_jk_*. Note that there is no restriction stating maker genes cannot be expressed in other cell/domain types. We assume the covariance 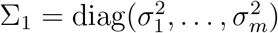. This simplification significantly reduces the computational cost.

For the high-dimensional non-marker genes, SpatialAnno models their centered normalized expression vector, ***x***_2*i*_, and latent label, *y_i_*, as

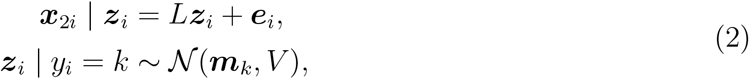

where factor ***z**_i_* ∈ *R^q^* represents a *q*-dimensional embedding of ***x***_2*i*_; *L* is a *p* × *q* factor loading matrix; ***m**_k_* ∈ *R^q^* is the mean vector for the *k*th cell/domain type, and *V* is the covariance matrix that is shared across cell/domain types; and ***e**_i_* is the residual error and follows an independent normal distribution with mean zero and variance Λ, which is a diagonal matrix, or 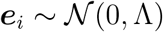.

#### Potts model for cell/domain labels

In the analysis of SRT datasets, the neighboring locations on the same tissue section often have similar cell/domain types. Thus, spots in neighboring locations contain immense amounts of information for annotating locations of interest. To promote neighborhood similarity in cell/domain types, we follow previous computation ^19,47^ and assume that cell/domain type *y_i_* ∈ {1*,…, K*} follows a Potts model characterized by an interaction parameter *ξ* and a neighborhood graph 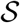,

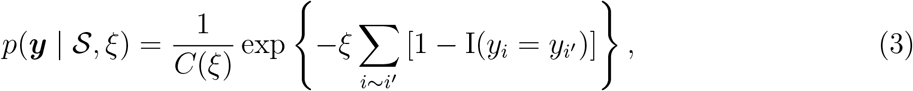

where *i* ~ *i*′ denotes all neighboring pairs in the neighborhood graph 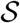; I(*y_i_* = *y_i_′*) is an indicator function that equals 1 if both the *i*th and *i*′th locations belong to the same cell/domain type and equals 0 otherwise; *ξ* is an unknown interaction parameter that determines the extent of cell/domain type similarity among neighboring locations; and *C*(*ξ*) is the normalizing constant, also known as the partition function that ensures the above probability mass function has a summation of one across all possible configurations of ***y***.

SpatialAnno modulates the over-expression of marker genes in Equation (1), the high-dimensional non-marker gene expressions in Equation (2), and the spatial smoothness across spots with a Potts model in Equation (3). The hierarchical probabilistic framework of SpatialAnno enables us to develop an efficient optimization algorithm through restricted expectation-maximization (EM)^48^ to estimate the probability of each location of a given cell/domain type. Briefly, our algorithm treats all parameters 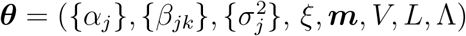 as unknown and estimates these parameters based on the data at hand to ensure optimal annotation performance. Algorithm details are provided in the Supplementary Notes.

SpatialAnno has several advantages that facilitate highly accurate assignments and various downstream analyses of spatial transcriptomics. First, by modelling the spatial correlation as labels, SpatialAnno borrows the cell-type information across spatial locations for spatially informed cell/domain type annotation. Second, SpatialAnno models the high-dimensional expression values of non-marker genes with the factor model, which can efficiently utilize the expression of non-marker genes to help verify and adjust label assignments. Third, modelling the high-dimensional expression values of non-marker genes allows SpatialAnno to infer cell-type-relevant embeddings, facilitating effective spatial transcriptomics visualization and spatial trajectory inference.

#### Spatial annotation and cell/domain relevant embeddings

To leverage the spatial location information, we construct a neighborhood graph 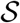 among locations by identifying the nearest neighbors for each spot. Specifically, the neighborhood *N_i_* for a spot *i* is defined by applying a proximity threshold. Let 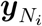 denote the configurations of the neighbors of spot *i*. The probability that spot *i* is associated with cell/domain type *k* given ***x**_i_*_1_, ***x**_i_*_2_ and its neighbor configuration 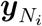 is specified by the following equation (Supplementary Notes) :

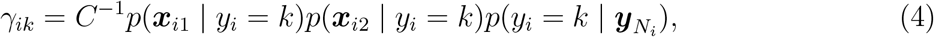

where *C* is a normalization constant. In the right-hand side, *p*(***x**_i_*_1_ | *y_i_* = *k*) and *p*(***x**_i_*_2_ | *y_i_* = *k*) model the effect of the expression levels of marker and non-marker genes, respectively, whereas 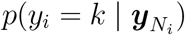 accounts for the effect of the neighbor configuration. The last term is determined by Eq. (3).

A key feature of SpatialAnno is its ability to extract cell/domain relevant embeddings for each spot. By modelling the expression levels of non-marker genes with factor models, SpatialAnno can extract cell/domain-type aware embeddings that can facilitate downstream analyses. Based on Eq. (2) and Bayes’ theorem, the conditional distribution of latent factors ***z**_i_* given (***x**_i_*_1_, ***x**_i_*_2_, *y_i_* = *k*) follows a multivariate normal distribution 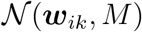 with mean ***w**_ik_* and variance *M* (Supplementary Notes). The low-dimensional embeddings for spot *i* are estimated by the posterior expectation of its latent factors ***z**_i_*:

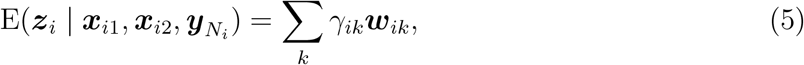

which are weighted averages that take into account the relative importance of each type. In this way, the embeddings are encouraged to be label-relevant.

### Compared methods and evaluation metrics

On all the simulated datasets and real datasets, we compared SpatialAnno with four annotation methods: (1) SCINA^12^ implemented in the R package *SCINA* (version 1.2.0); (2) Garnett^13^ implemented in the R package *garnett* (version 0.1.21); (3) CellAssign ^14^ implemented in the R package *cellassign* (version 0.99.21); and (4) scSorter ^15^ implemented in the R package *scSorter* (version 0.0.2). We used the recommended default parameter settings in their tutorials. Among these methods, SCINA and CellAssign only use the expression of marker genes, and scSorter and Garnett can borrow information from the expression of non-marker genes. We evaluated the annotation performance using three metrics, i.e., Kappa, mF1 score, and ACC, as suggested in previous annotation studies of single-cell data ^11,49^. ACC was defined as the proportion of spots that were classified into the correct types. Kappa is generally thought to be a more robust measure than ACC, as it takes into account the possibility of the agreement occurring by chance. The cell-level F1 score considers each cell to be an individual classification task with a true cell-type assignment (and potentially multiple incorrect cell-type assignments) for the purposes of calculating precision and recall (Supplementary Notes).

We also compared the low-dimensional embeddings estimated in SpatialAnno with those from PCA and DR-SC^19^. In detail, we first extracted the top 15-dimensional components and then summarized those top components as three tSNE components and visualizing the resulting tSNE components with RGB colors in the RGB plot. To show that the estimated embeddings carry the most information about cell/domain types, we evaluated the conditional correlation coefficients between the true cell/domain labels and the observed gene expression, given the estimated embeddings in SpatialAnno. Furthermore, the embeddings in SpatialAnno improve clustering performance. With embeddings from SpatialAnno, PCA, and DR-SC, we performed clustering analysis using the Louvain community detection algorithm implemented in the R package *Seurat* (version 4.1.1), and evaluated clustering performance using the ARI ^50^.

### Simulations

We performed comprehensive simulations to evaluate the performance of SpatialAnno and compared it with that of alternative annotation methods. The spatial locations of 3639 spots were taken from DLPFC section 151673. Cell/domain types were assigned with the manually annotations from the original studies ^20^. We simulated gene expression data for each spot using the *splatter* package (version 1.20.0). The parameter for the proportion of DEGs (*de.prob*) in each layer was set to 0.5. The DE strength was determined by both the mean parameter *de.facloc* and scale parameter *de.facScale*, the former ranges from 0.1 to 0.8, and the latter was set to within [0.1,1], corresponding to the log fold change in expression from one-fold to two-fold across different types. All the other parameters were set based on their estimates in the seven layers from DLPFC section 151673.

Five marker genes for each cell/domain type were selected from the top DEGs based on log-fold change in expression. We tested the accuracy and robustness of SpatialAnno with the following settings that reflect real-world scenarios.

I. To test the robustness of SpatialAnno to the erroneous specification of the number of cell/domain types, we considered three scenarios. In the first scenario, marker genes for all seven cell/domain types were provided and no unknown cell/domain types existed in the expression data. In the second scenario, marker genes for two cell/domain types were removed to create a scenario in which fewer cell/domain types were specified in the marker gene matrix than actually exist in the data. Thus, cells from these two cell/domain types should be assigned to “unknown”. In the third scenario, the marker genes for nine cell/domain types were added, but two cell/domain types did not appear in the expression data. This mimics a scenario in which there are more cell/domain types are specified in the marker gene matrix than actually exist in the data.
II. To evaluate the robustness of SpatialAnno to the marker gene misspecification, we next created a scenario in which marker genes may be incomplete or incorrect. We randomly flipped a fraction of entries in the binary marker gene matrix *ρ* to introduce errors. Specifically, the procedure consisted of two steps. In the first step, a proportion of entries in *ρ* that contained one were flipped. In the second step, the same number of entries flipped in the first step were flipped for the entries that contained zero in the original *ρ*. The considered proportions were set to be either 10%, 20%, or 30%. Other settings were similar to those in the first scenario in Simulation I.
III. To assess the capability of SpatialAnno to utilize high-dimensional non-marker genes, we varied the number of non-marker genes as 60, 100, 500, 1000, and 2000. In this setting, we only compared scSorter and Garnett, as only these methods can utilize non-marker genes. Other settings were similar to those in the first scenario in Simulation I.

For each simulation setting, we performed 50 replicate simulations. In each replicate, we applied SpatialAnno and the other methods to annotate each spot.

## Data analysis

### Normalization of SRT data

For all datasets, we normalized the raw expression count matrix using the variance stabilizing transformation function, SCTranscorm, provided in *Seurat* (version 4.1.1). We performed gene filtering using *SPARK* (version 1.1.1)^51^ for data with transcriptome-wide measurements. The most spatially variable genes (see Data resource) were selected as input for the annotation methods SpatialAnno, scSorter, SCINA, and Garnett. CellAssign took the raw count matrix of these genes as input.

### Selection of marker genes

We obtained a marker gene list primarily following the protocols of CellAssign. We (1) performed differential expression analysis of a reference scRNA-seq/SRT data using the FindAllMarkers function in the R package *Seurat* (version 4.1.1) and selected the top 30 DEGs ordered by the log2(fold-change) with upregulation, (2) removed those with an insignificant adjusted *p*-value and those detection percentages across different cell/domain types were similar (differences between pct.1 and pct.2 values from the FindAllMarkers function are lower than 0.3), and (3) filtered out genes that were of low expression in the spatial transcriptomic data. We finally selected the top-ranked genes with the smallest *p*-values.

### Clustering analysis

To examine the information captured by SpaitalAnno embeddings of non-marker genes, we performed clustering analysis using three different sets of embeddings as input in both the simulated and DLPFC data. The three embedding sets include the top 15 PCs in marker genes by PCA, 15-dimensional embeddings in non-marker genes by SpatialAnno, and combined. We then performed clustering analysis using the Louvain community detection algorithm.

### Spatial trajectory inference

To construct a spatial map of the DLPFC Visium data, we employed the PAGA algorithm ^22^ implemented in the Python package *SCANPY* (version 1.9.1) to preserve the global topology in the embeddings of non-marker genes. The cluster labels for PCA embeddings and DR-SC embeddings were estimated using the spatial clustering methods implemented in the R packages *BayesSpace* (version 1.5.1)^52^ and *DR-SC* (version 2.9.0) ^19^, respectively.

To estimate the developmental trajectories among the various locations in the brain regions, we applied Slingshot^53^ to the low-dimensional embeddings. As SpatialAnno only extracts embeddings of non-marker genes, we combined them with the embeddings of marker genes by PCA. The cluster labels used in Slingshot were obtained from the spatial clustering method DR-SC^19^. To detect DEGs along the estimated pseudotime, we used the function testPseudotime in the R package *TSCAN* (version 1.37.0) ^54^.

## Data resources

### Human dorsolateral prefrontal cortex data obtained by 10x Visium

We downloaded a human DLPFC data set^20^ that was generated by the 10x Visium platform from http://spatial.libd.org/spatialLIBD/. In this dataset, there were 12 tissue sections, which contained a total of 33,538 genes measured on average over 3973 spots. We used the sample ID151673, which contains expression measurements of 33,538 genes on 3639 spots, as the main analysis example. We presented the results for the other 11 samples in the Supplementary Figures. For all the sections, we extracted the top 2000 spatially variable genes with SPARK-X^51^ before performing annotations.

To identify layer-specific marker genes for annotation, we used tissue section 151507 as the reference data. This dataset contained 33,538 genes for 4,226 spots. For each layer, the top 5 DEGs were selected as its marker genes. The final marker gene list is available in Supplementary Table 6.

### Mouse olfactory bulb data by spatial transcriptomics (ST)

We obtained the mouse olfactory bulb ST data from the spatial transcriptomics research website (https://www.spatialresearch.org/). This data consists of gene expression levels in the form of read counts that were collected for a number of spatial locations. We followed the methods of previous studies^55,56^ to focus on the mouse OB section 12, which contains 16,034 genes and 282 spatial locations. We presented the results for the other 11 sections in the Supplementary Figures. We extracted the top 3000 most highly variable genes with function SCTransform implemented in *Seurat* (version 4.0.5) ^57^ before performing annotations.

To construct the marker gene list for annotation, we perform differential expression analysis on scRNA-seq data^23^ from the Gene Expression Omnibus (GEO; accession number GSE121891). This scRNA-seq data was collected from the mouse olfactory bulb and contains 18,560 genes and 12,801 cells for five cell types: granule cells (GC, *n*=8614), olfactory sensory neurons (OSNs, *n* = 1200), periglomerular cells (PGC, *n* = 1693), mitral and tufted cells (M-TC, *n* = 1133), and external plexiform layer interneurons (EPL-IN, *n* = 161). For each cell type, the top four DEGs were selected as its marker genes. The final marker gene list is available in Supplementary Table 7.

### Mouse hippocampus Slide-seq data and Slide-seqV2 data

We obtained the mouse hippocampus Slide-seq dataset and Slide-seqV2 dataset^3^ from the Broad Institute’s Single Cell Portal (https://singlecell.broadinstitute.org/single_cell/study/SCP948/robust-decomposition-of-cell-type-mixtures-in-spatial-transcriptomics) The Slide-seq dataset consists of gene expression measurements in the form of read counts for 22,457 genes and 34,199 spatial locations. The Slide-seqV2 dataset consists of gene expression measurements in the form of read counts for 23,264 genes and 53,208 spatial locations. In the analysis, we filtered out genes that had fewer than 20 counts on all locations and filtered out locations that had fewer than 20 genes with nonzero counts. These filtering criteria led to final sets of 14,481 genes and 31,664 cells for Slide-seq dataset, and 16,121 genes and 51,212 cells for Slide-seqV2 dataset. In addtion, for both datasets, we extracted the top 2000 most spatially variable genes with SPARK-X^51^ before performing annotations.

To construct marker genes for annotation, we obtained the DropViZ scRNA-seq dataset^58^ from the Broad Institute’s Single Cell Portal. This data was collected from the mouse hippocampus, which contained 22,245 genes and 52,846 cells for 19 cell types. For each cell type, the top five DEGs were selected as marker genes. Besides the 19 cell types, we added another two cell types, *Slc17a6* neurons and Hb neurons, and their marker genes were extracted from the original study^58^. The final marker gene list used is available in Supplementary Table 8.

### Mouse embryo data by seqFISH

We obtained the mouse embryo seqFISH data^2^ from https://marionilab.cruk.cam.ac.uk/SpatialMouseAtlas/. This dataset consists of 387 selected target genes from three mouse embryo tissue sections for 19,451, 14,891, and 23,194 cells, respectively. We calculated normalized expression log counts for each cell using logNormCounts function in the R package *scuttle* ^59^ with cell-specific size factors.

To construct the marker gene list, we used Embryo 3 as a reference; this dataset contains 24 cell types. For each cell type, the top eight DEGs were selected as marker genes. We removed marker genes for two cell types, ExE endoderm cells and blood progenitors, as there were too few (less than 30) of these cells. The cell type “Low quality” was also removed. The final marker gene list used contained 21 cell types and is available in Supplementary Table 9.

## Data availability

This study made use of publicly available datasets. These include the mouse OB dataset (https://www.spatialresearch.org/), DLPFC dataset on the 10x Visium platform are accessible at (https://github.com/LieberInstitute/spatialLIBD), seqFISH dataset (https://doi.org/10.18129/B9.bioc.MouseGastrulationData), and mouse hippocampus Slide-seq and Slide-seqV2 datasets (https://singlecell.broadinstitute.org/single_cell/study/SCP948/robust-decomposition-of-cell-type-mixtures-in-spatial-transcriptomics).

## Code availability

The SpatialAnno software and source code have been deposited at https://github.com/Shufeyangyi2015310117/SpatialAnno. Example codes for using SpatialAnno are publicly available at https://shufeyangyi2015310117.github.io/SpatialAnno/index.html. All scripts used to reproduce all the analyses can be found at https://github.com/Shufeyangyi2015310117/SpatialAnno_Analysis and https://doi.org/10.5281/zenodo.7413083.

## Author contributions

X.S. and J.L. initiated and designed the study. X.S. and Y.Y. developed the method, implemented the software, performed simulations and analyzed real data. X.S. and J.L. wrote the manuscript, and all authors edited and revised the manuscript.

