## Supplementary Figures and Tables for "Probabilistic cell/domain-type assignment of spatial transcriptomics data with SpatialAnno"

#### Supplementary Figure 1. Additional simulation results

**a** Kappa, mF1 and ACC of SpatialAnno, scSorter, and Garnett for simulation data, with different numbers of non-marker genes provided as input. Two-sided Wilcoxon Rank Sum test was used to pair wisely test the difference between metrices with 60 and 2000 non-marker genes, and the *p*-value is shown. **b** Clustering results measured by adjusted rand index (ARI, the higher the better) using low-dimensional embeddings either from marker genes by PCA or non-marker genes by SpatialAnno, or combined for simulation data, providing different number of cell/domain types with marker genes as input. **c** As in **b**, with various degrees of mis-specification in marker genes. **d** Pearson's correlation coefficients between observed expression and the inferred labels, conditioned on embeddings from SpatialAnno, PCA, and DR-SC, providing different number of cell/domain types with marker genes as input. **e** As in **d**, with various degrees of mis-specification in marker genes. **f** Runtime (in seconds) benchmarking of different methods with different number of cell/domain types.

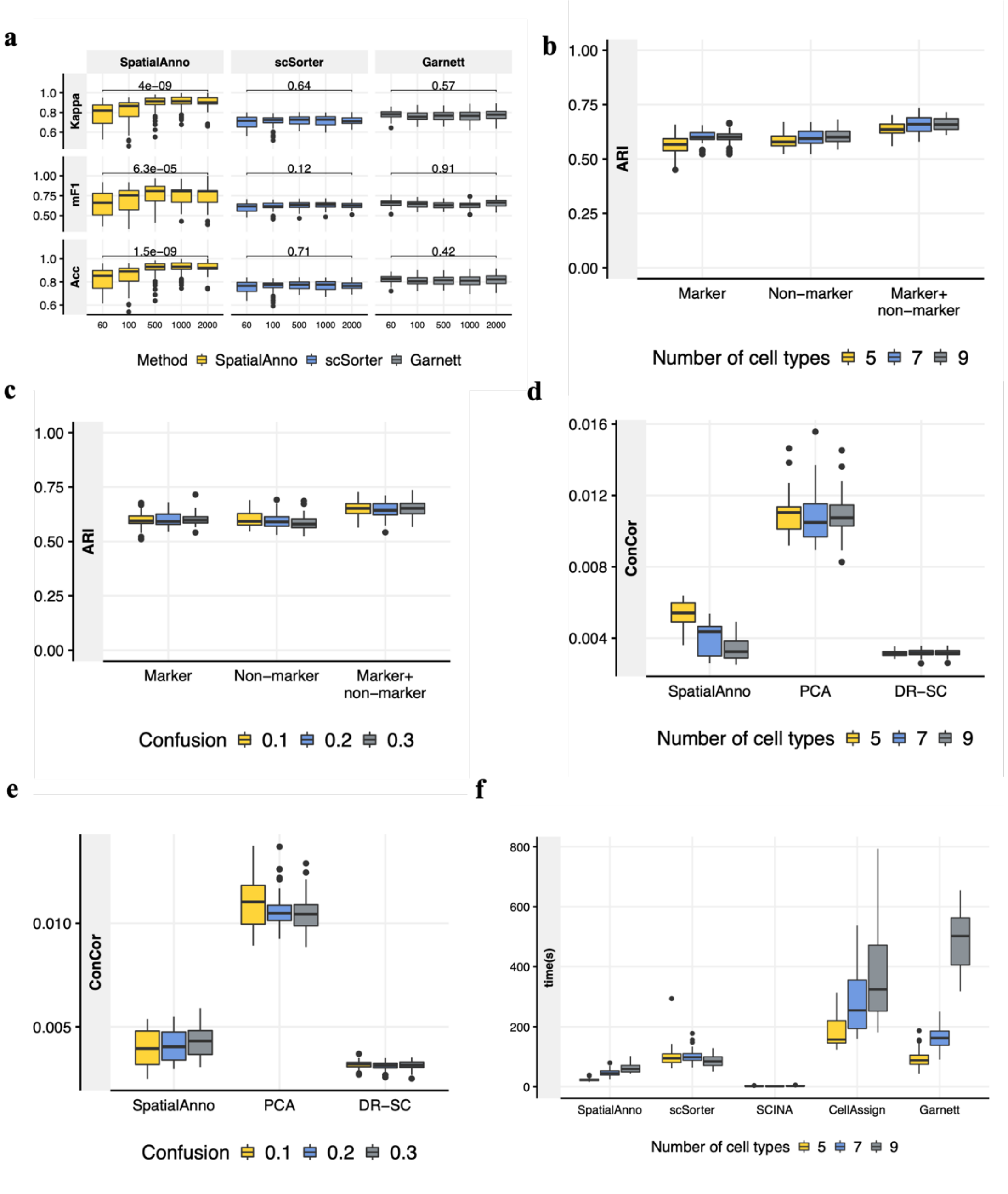

**Supplementary Figure 2. Spatial domain annotation in the DLPFC section 151507**
**a** Spatial domain annotations of tissue section 151507 are shown for ground truth, SpatialAnno, scSorter, SCINA, Garnett, and CellAssign. **b** Top, annotation by SpatialAnno for each spot. Bottom, expression levels of corresponding layer-specific marker genes. **c** Top, RGB plots for low-dimensional embedding inferred by SpatialAnno, PCA, and DR-SC. As end-to-end annotation approaches, scSorter, SCINA, Garnett, and CellAssign cannot be utilized to extract low-dimensional embedding. Bottom, PAGA graphs generated by SpatialAnno, PCA, and DR-SC embeddings.

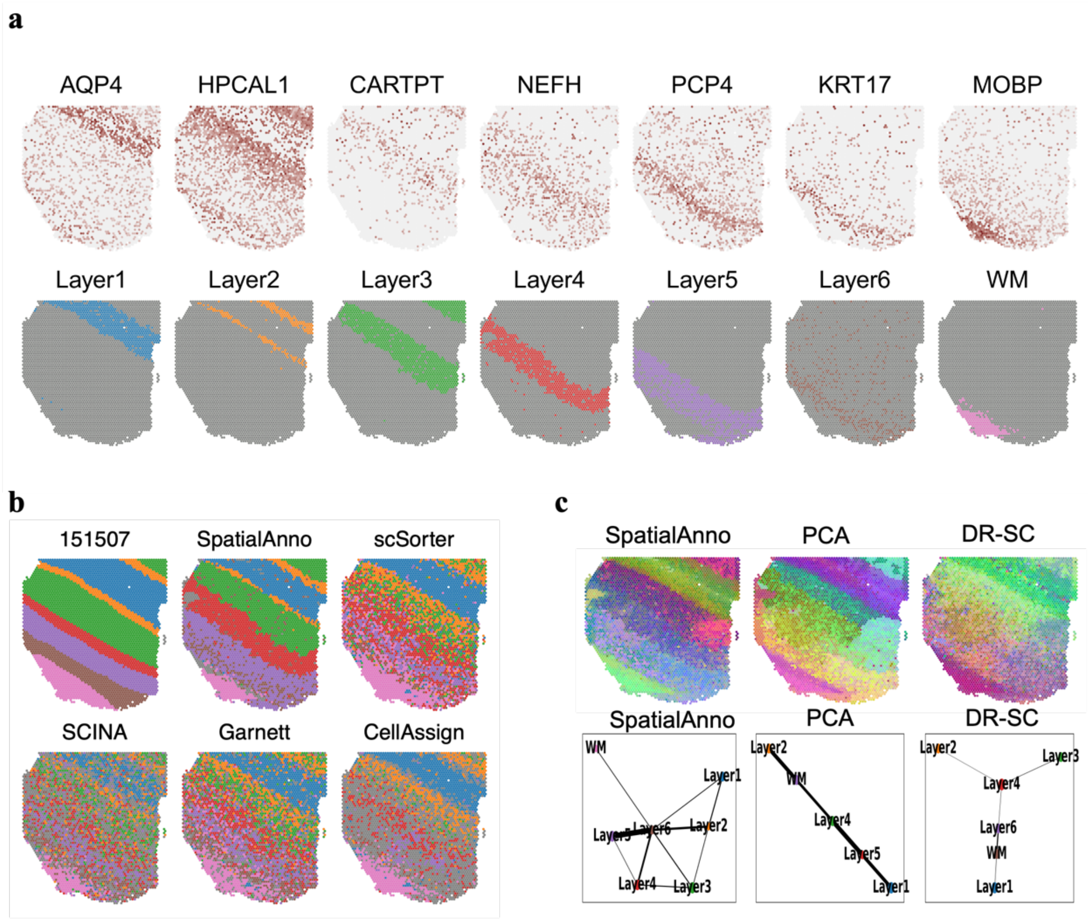

**Supplementary Figure 3. Spatial domain annotation of DLPFC section 151508**
**a** Spatial domain annotations of tissue section 151508 are shown for ground truth, SpatialAnno, scSorter, SCINA, Garnett, and CellAssign. **b** Top, annotation by SpatialAnno for each spot. Bottom, expression levels of corresponding layer-specific marker genes. **c** Top, RGB plots for the low dimensional embedding inferred by SpatialAnno, PCA, and DR-SC. As end-to-end annotation approaches, scSorter, SCINA, Garnett, and CellAssign cannot be utilized to extract low-dimensional embeddings. Bottom, PAGA graphs generated by SpatialAnno, PCA, and DR-SC embeddings.

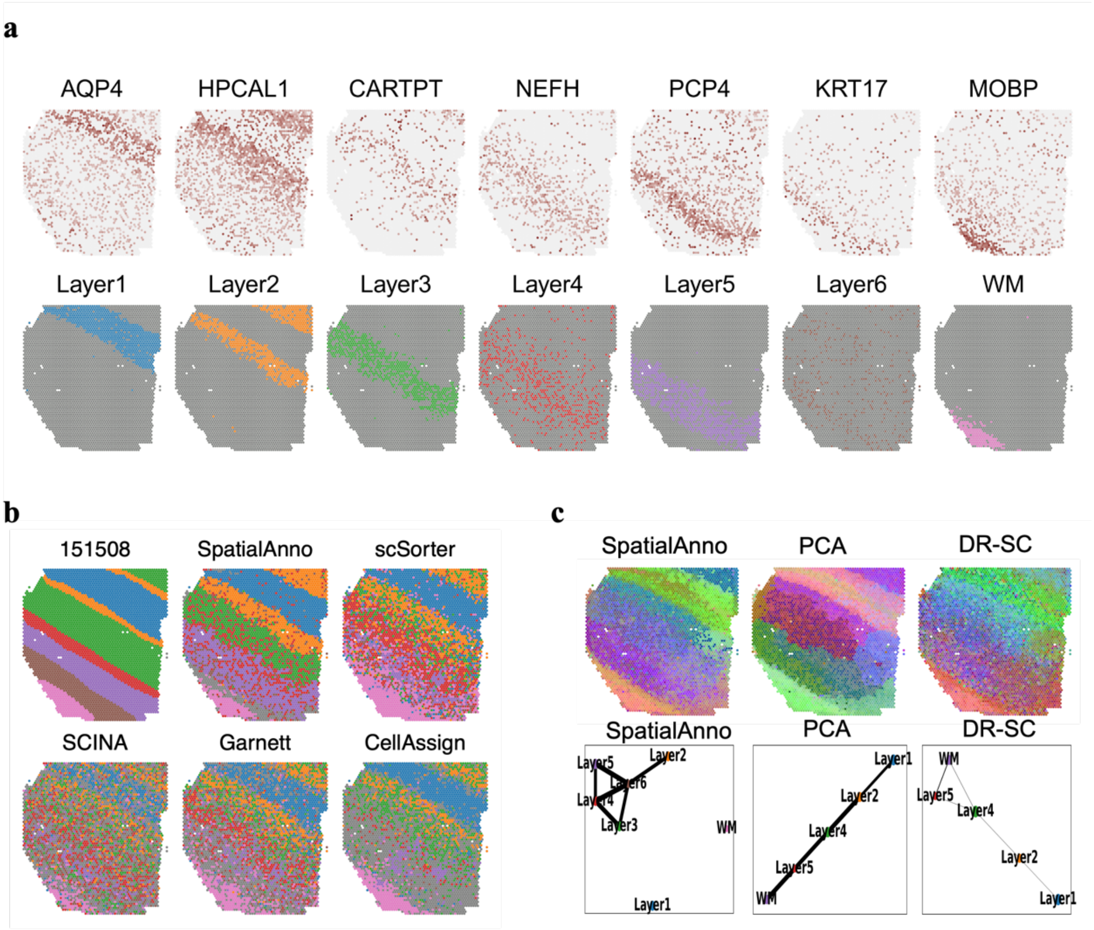

**Supplementary Figure 4. Spatial domain annotation of DLPFC section 151509**

**a** Spatial domain annotations of tissue section 151509 are shown for ground truth, SpatialAnno, scSorter, SCINA, Garnett, and CellAssign. **b** Top, annotation by SpatialAnno for each spot. Bottom, expression levels of corresponding layer-specific marker genes. **c** Top, RGB plots for low-dimensional embedding inferred by SpatialAnno, PCA, and DR-SC. As end-to-end annotation approaches, scSorter, SCINA, Garnett, and CellAssign cannot be utilized to extract low-dimensional embeddings. Bottom, PAGA graphs generated by SpatialAnno, PCA, and DR-SC embeddings.

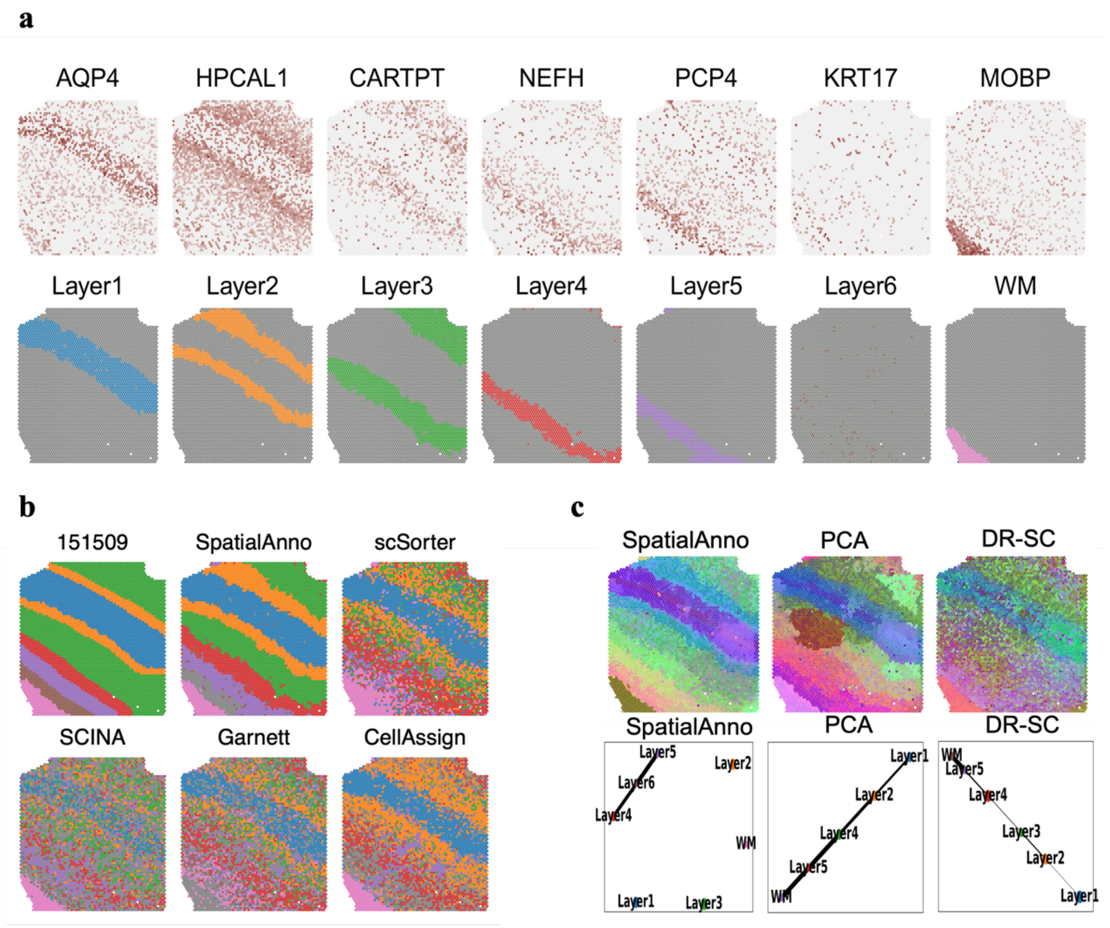

**Supplementary Figure 5. Spatial domain annotation of DLPFC section 151510**

**a** Spatial domain annotations of tissue section 151510 are shown for ground truth, SpatialAnno, scSorter, SCINA, Garnett, and CellAssign. **b** Top, annotation of SpatialAnno for each spot. Bottom, expression levels of corresponding layer-specific marker genes. **c** Top, RGB plots for low-dimensional embedding inferred by SpatialAnno, PCA, and DR-SC. As end-to-end annotation approaches, scSorter, SCINA, Garnett, and CellAssign cannot be utilized to extract low-dimensional embeddings. Bottom, PAGA graphs generated by SpatialAnno, PCA, and DR-SC embeddings.

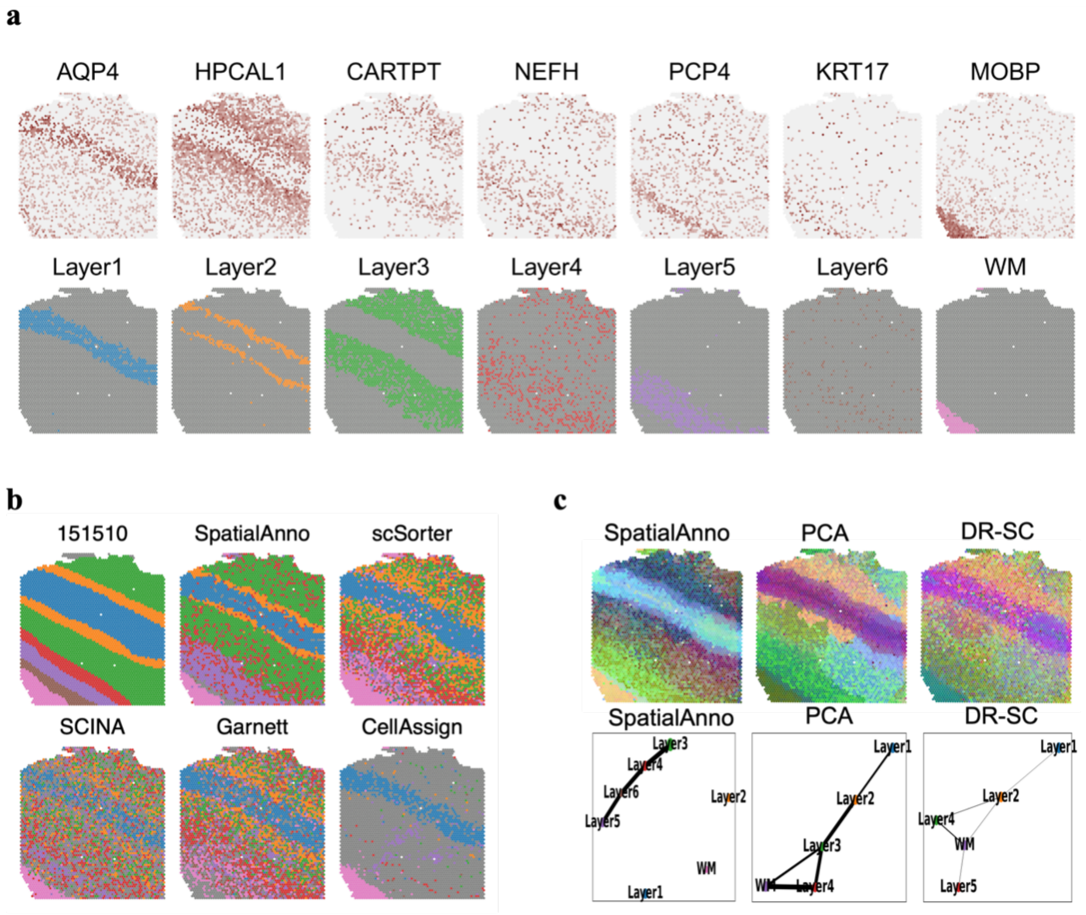

**Supplementary Figure 6. Spatial domain annotation of DLPFC section 151669**

**a** Spatial domain annotations of tissue section 151669 are shown for ground truth, SpatialAnno, scSorter, SCINA, Garnett, and CellAssign. **b** Top, annotation of SpatialAnno for each spot. Bottom, expression levels of corresponding layer-specific marker genes. **c** Top, RGB plots for low-dimensional embedding inferred by SpatialAnno, PCA, and DR-SC. As end-to-end annotation approaches, scSorter, SCINA, Garnett, and CellAssign cannot be utilized to extract low-dimensional embeddings. Bottom, PAGA graphs generated by SpatialAnno, PCA, and DR-SC embeddings.

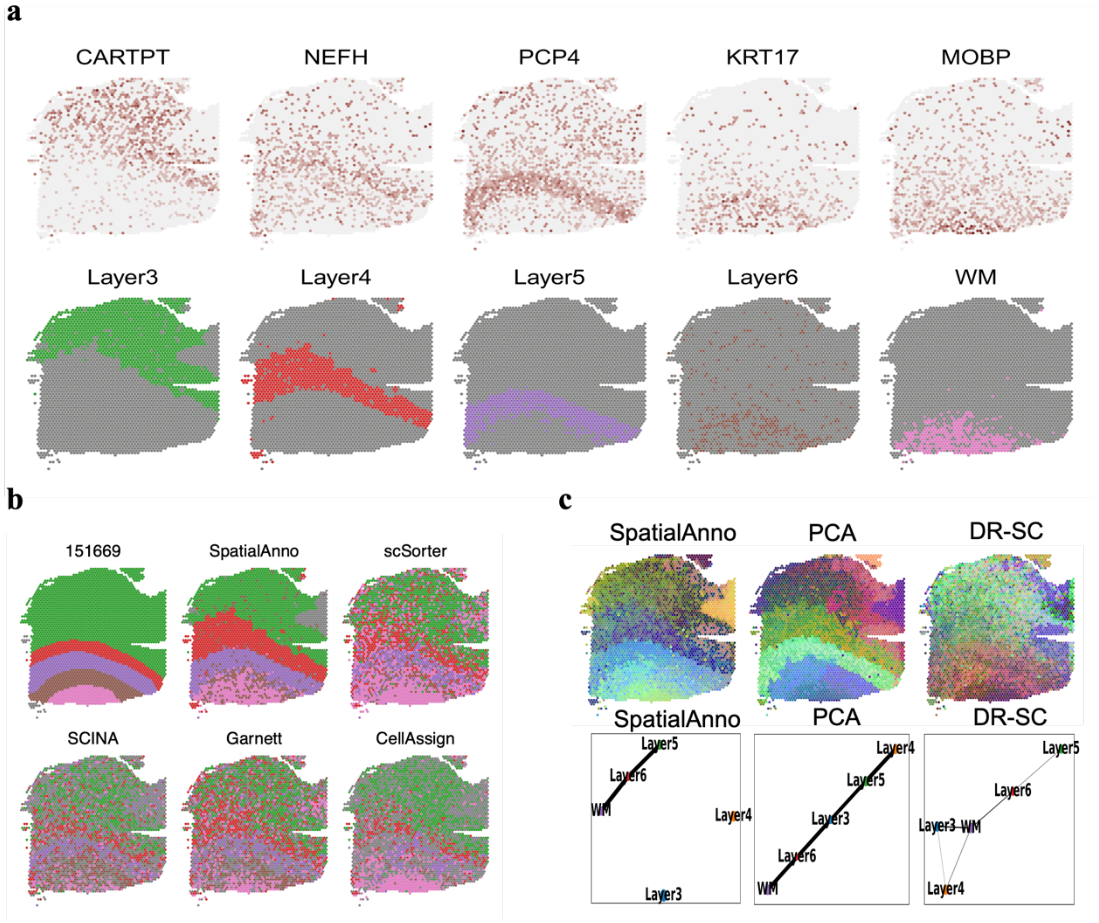

**Supplementary Figure 7. Spatial domain annotation of DLPFC section 151670**

**a** Spatial domain annotations of tissue section 151670 are shown for ground truth, SpatialAnno, scSorter, SCINA, Garnett, and CellAssign. **b** Top, annotation of SpatialAnno for each spot. Bottom, expression levels of corresponding layer-specific marker genes. **c** Top, RGB plots for low-dimensional embedding inferred by SpatialAnno, PCA, and DR-SC. As end-to-end annotation approaches, scSorter, SCINA, Garnett, and CellAssign cannot be utilized to extract low-dimensional embeddings. Bottom, PAGA graphs generated by SpatialAnno, PCA, and DR-SC embeddings.

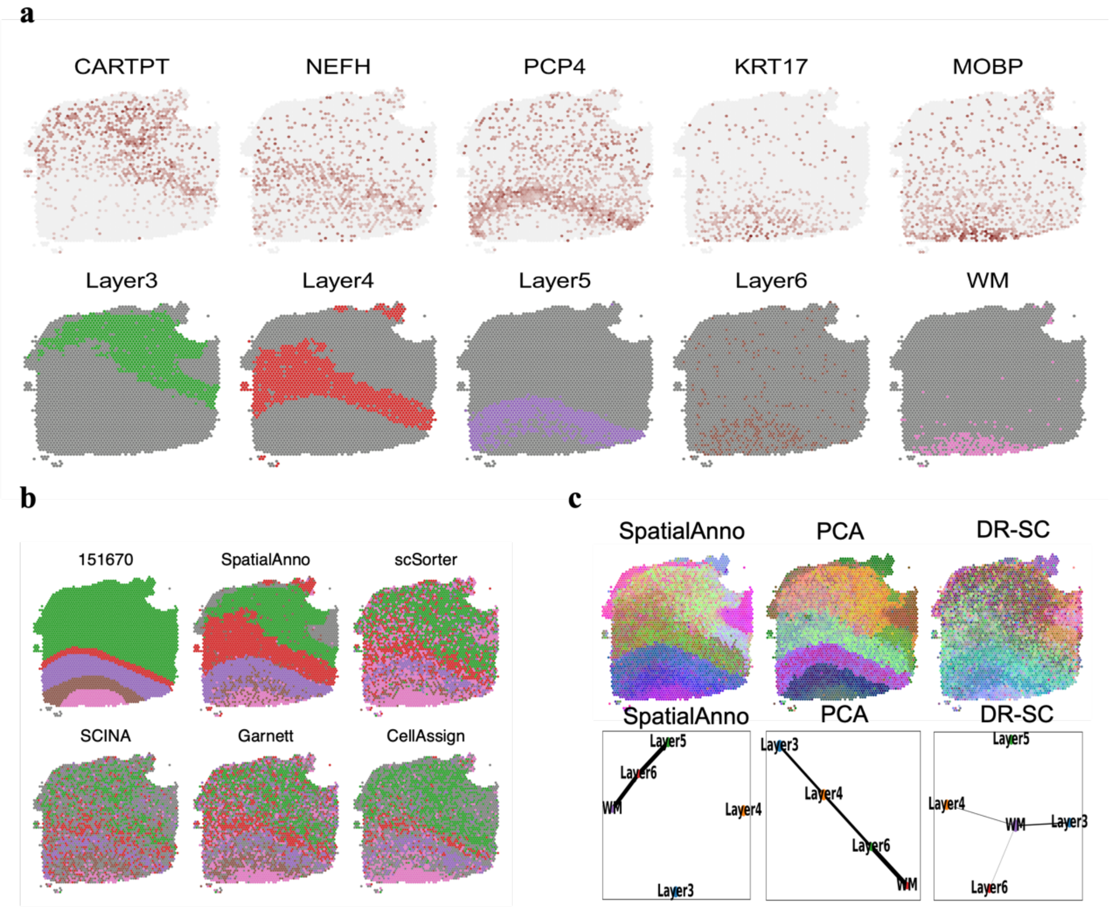

**Supplementary Figure 8. Spatial domain annotation of DLPFC section 151671**

**a** Spatial domain annotations of tissue section 151671 are shown for ground truth, SpatialAnno, scSorter, SCINA, Garnett, and CellAssign. **b** Top, annotation of SpatialAnno for each spot. Bottom, expression levels of corresponding layer-specific marker genes. **c** Top, RGB plots for low-dimensional embedding inferred by SpatialAnno, PCA, and DR-SC. As end-to-end annotation approaches, scSorter, SCINA, Garnett, and CellAssign cannot be utilized to extract low-dimensional embeddings. Bottom, PAGA graphs generated by SpatialAnno, PCA, and DR-SC embeddings.

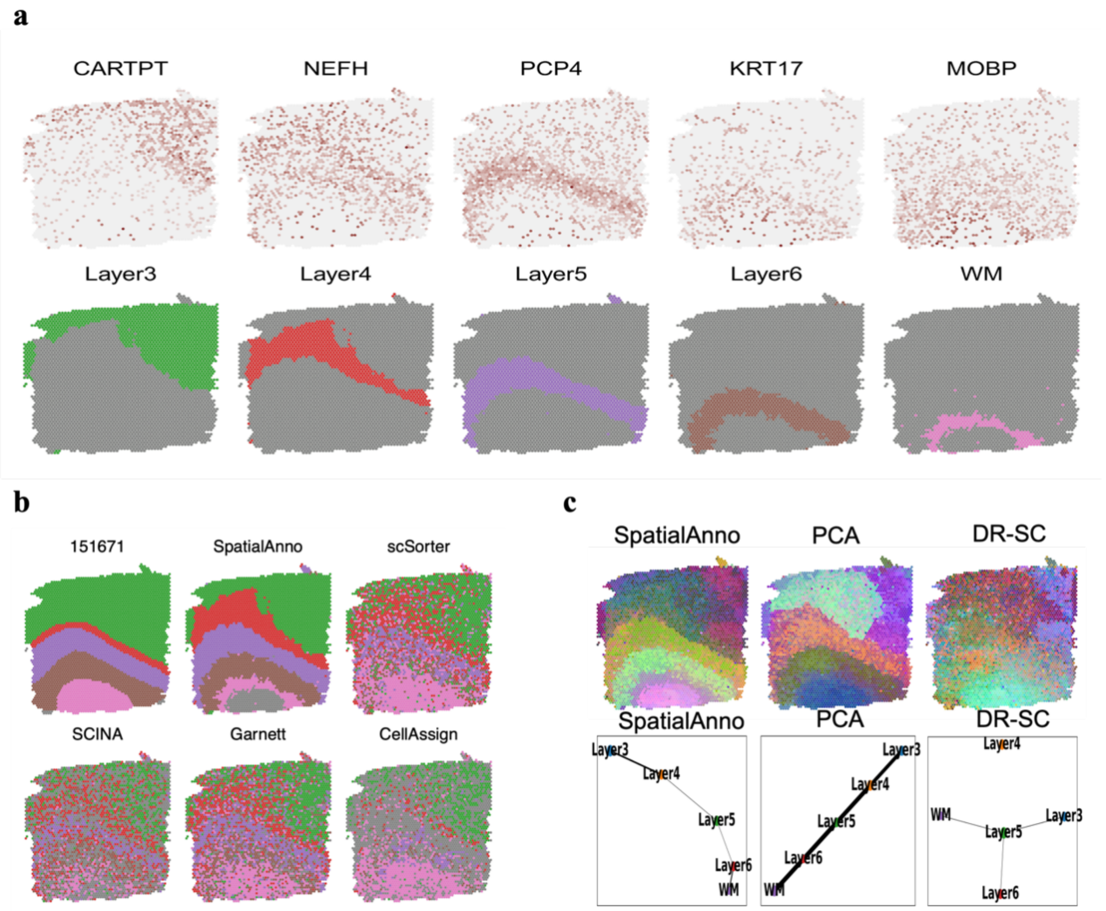

**Supplementary Figure 9. Spatial domain annotation of DLPFC section 151672**

**a** Spatial domain annotation of tissue section 151672 are shown for ground truth, SpatialAnno, scSorter, SCINA, Garnett, and CellAssign. **b** Top, annotation of SpatialAnno for each spot. Bottom, expression levels of corresponding layer-specific marker genes. **c** Top, RGB plots for low-dimensional embedding inferred by SpatialAnno, PCA, and DR-SC. As end-to-end annotation approaches, scSorter, SCINA, Garnett, and CellAssign cannot be utilized to extract low-dimensional embeddings. Bottom, PAGA graphs generated by SpatialAnno, PCA, and DR-SC embeddings.

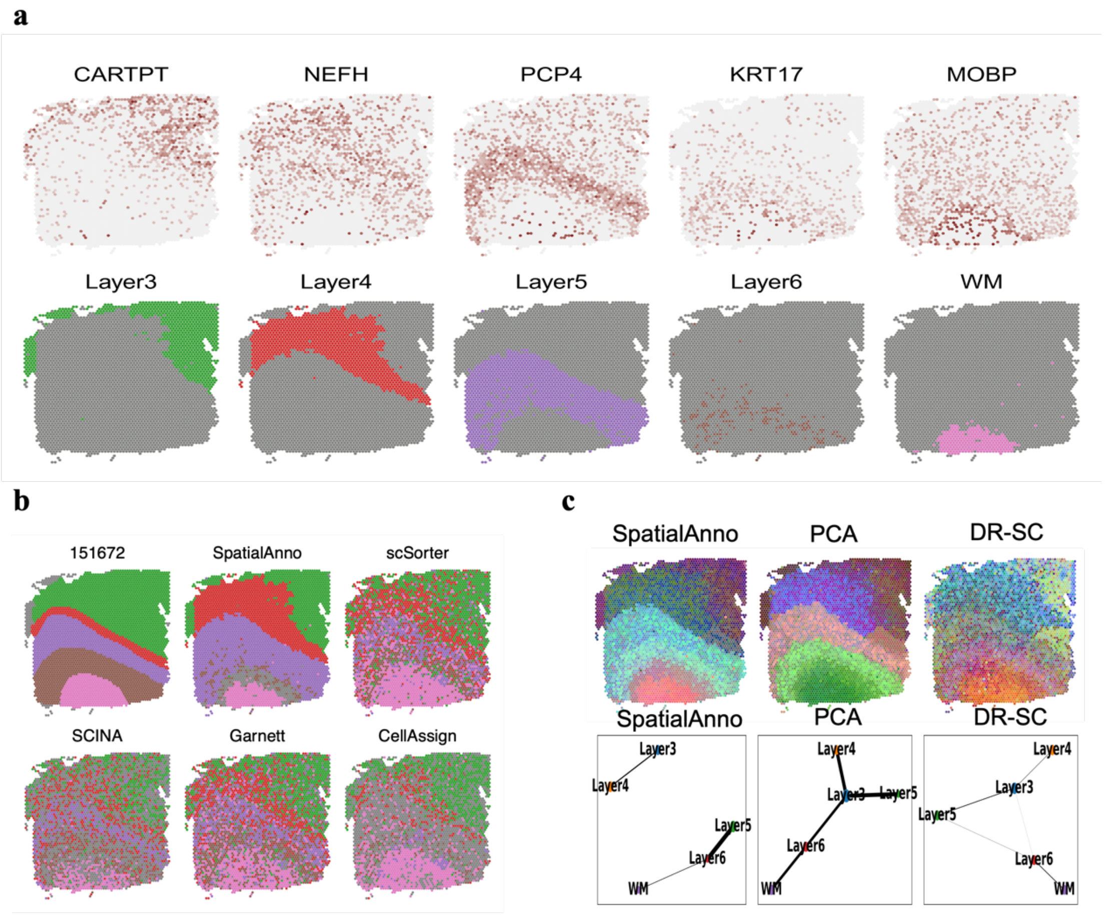

**Supplementary Figure 10. Spatial domain annotation of DLPFC section 151673**
**a** Spatial domain annotations of tissue section 151673 are shown for ground truth, SpatialAnno, scSorter, SCINA, Garnett, and CellAssign. **b** Top, annotation by SpatialAnno for each spot. Bottom, expression levels of corresponding layer-specific marker genes. **c** Top, RGB plots for low-dimensional embedding inferred by SpatialAnno, PCA, and DR-SC. As end-to-end annotation approaches, scSorter, SCINA, Garnett, and CellAssign cannot be utilized to extract low-dimensional embeddings. Bottom, PAGA graphs generated by SpatialAnno, PCA, and DR-SC embeddings.

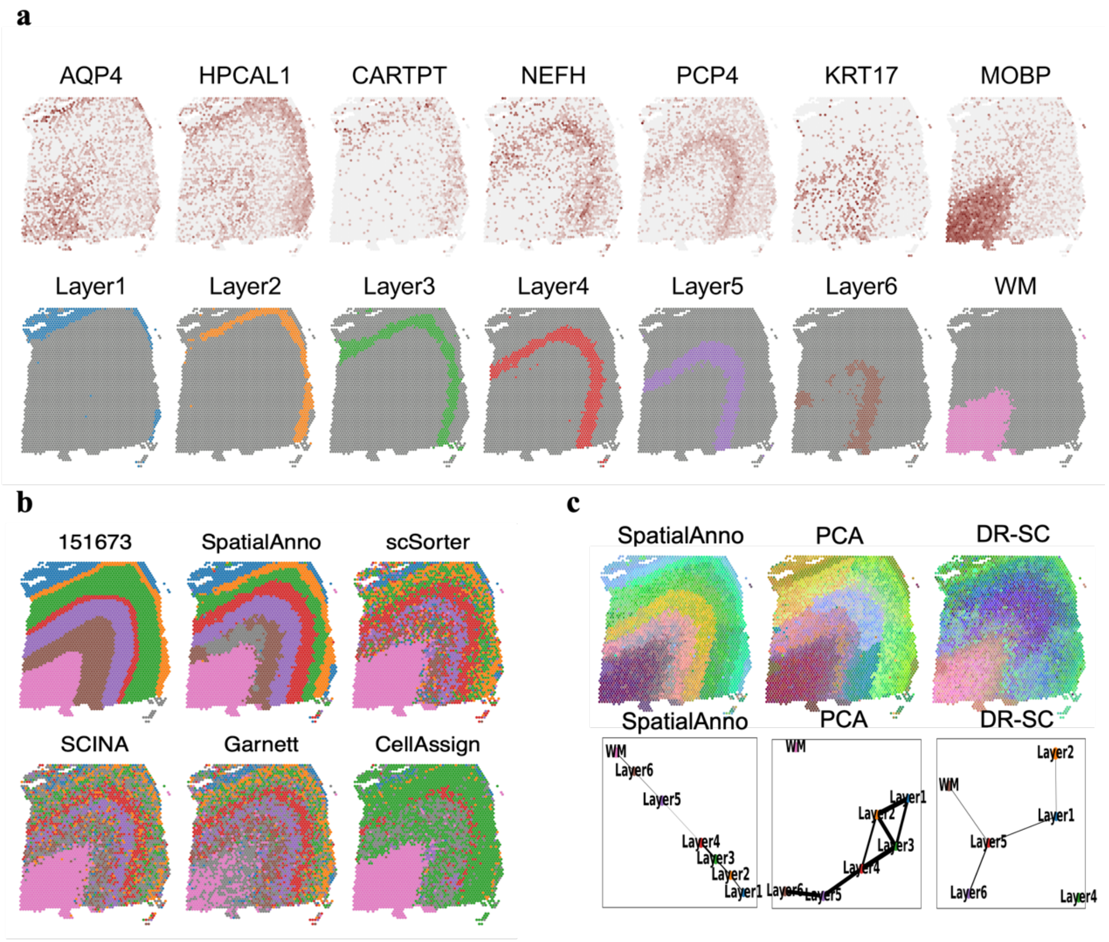

**Supplementary Figure 11. Spatial domain annotation of DLPFC section 151674**
**a** Spatial domain annotation of tissue section 151674 are shown for ground truth, SpatialAnno, scSorter, SCINA, Garnett, and CellAssign. **b** Top, annotation by SpatialAnno for each spot. Bottom, expression levels of corresponding layer-specific marker genes. **c** Top, RGB plots for low-dimensional embedding inferred by SpatialAnno, PCA, and DR-SC. As end-to-end annotation approaches, scSorter, SCINA, Garnett, and CellAssign cannot be utilized to extract low-dimensional embeddings. Bottom, PAGA graphs generated by SpatialAnno, PCA, and DR-SC embeddings.

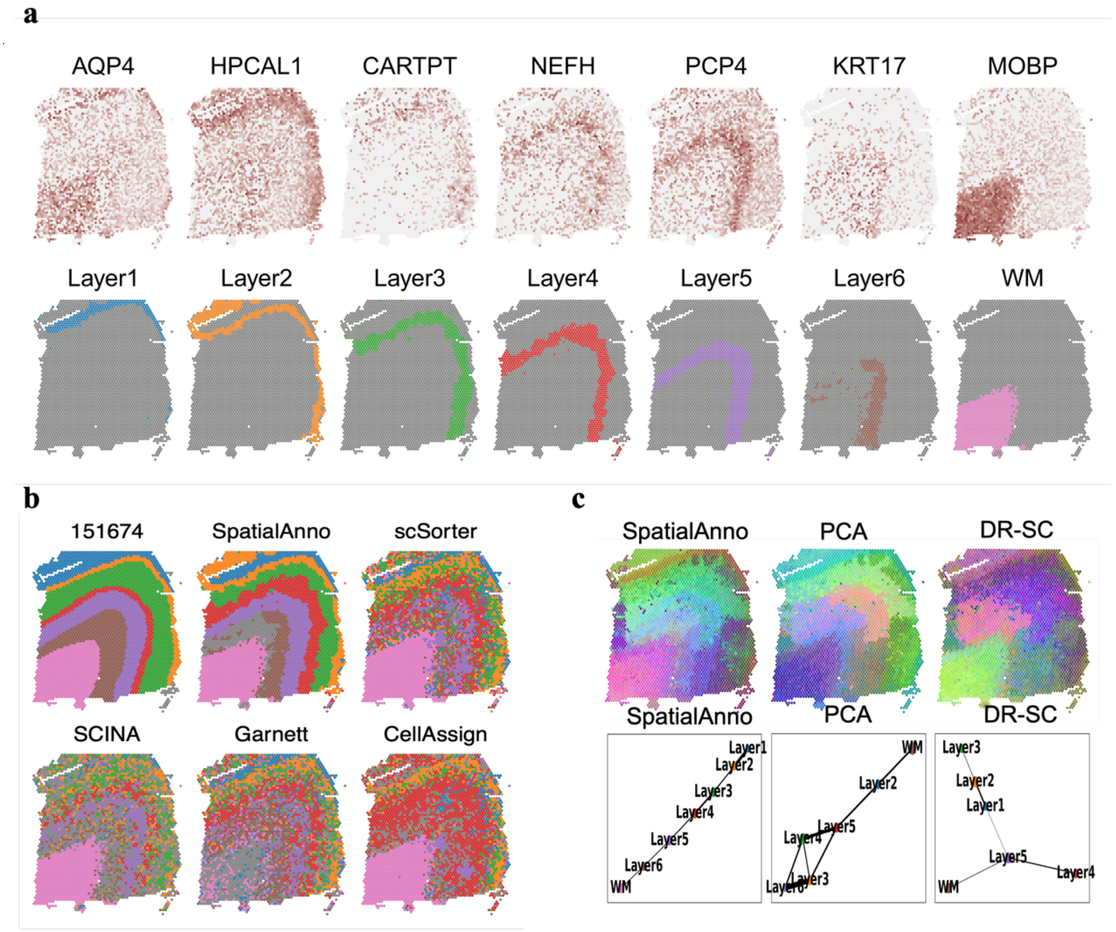

**Supplementary Figure 12. Spatial domain annotation of DLPFC section 151675**
**a** Spatial domain annotation of tissue section 151675 are shown for ground truth, SpatialAnno, scSorter, SCINA, Garnett, and CellAssign. **b** Top, annotation of SpatialAnno for each spot. Bottom, expression levels of corresponding layer-specific marker genes. **c** Top, RGB plots for low-dimensional embedding inferred by SpatialAnno, PCA, and DR-SC. As end-to-end annotation approaches, scSorter, SCINA, Garnett, and CellAssign cannot be utilized to extract low-dimensional embeddings. Bottom, PAGA graphs generated by SpatialAnno, PCA, and DR-SC embeddings.

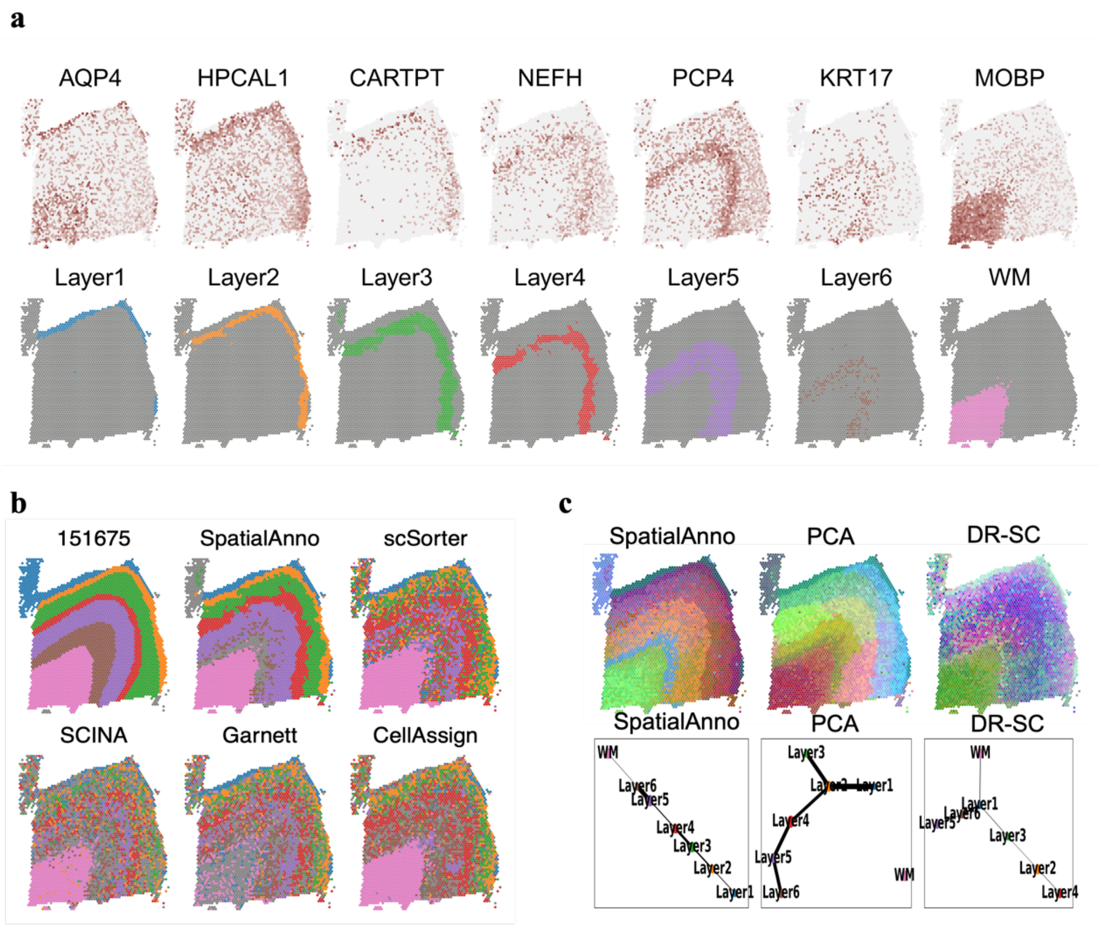

**Supplementary Figure 13. Spatial domain annotation of DLPFC Section 151676**
**a** Spatial domain annotation of tissue section 151676 are shown for ground truth, SpatialAnno, scSorter, SCINA, Garnett, and CellAssign. **b** Top, annotation by SpatialAnno for each spot. Bottom, expression levels of corresponding layer-specific marker genes. **c** Top, RGB plots for low-dimensional embedding inferred by SpatialAnno, PCA, and DR-SC. As end-to-end annotation approaches, scSorter, SCINA, Garnett, and CellAssign cannot be utilized to extract low-dimensional embeddings. Bottom, PAGA graphs generated by SpatialAnno, PCA, and DR-SC embeddings.

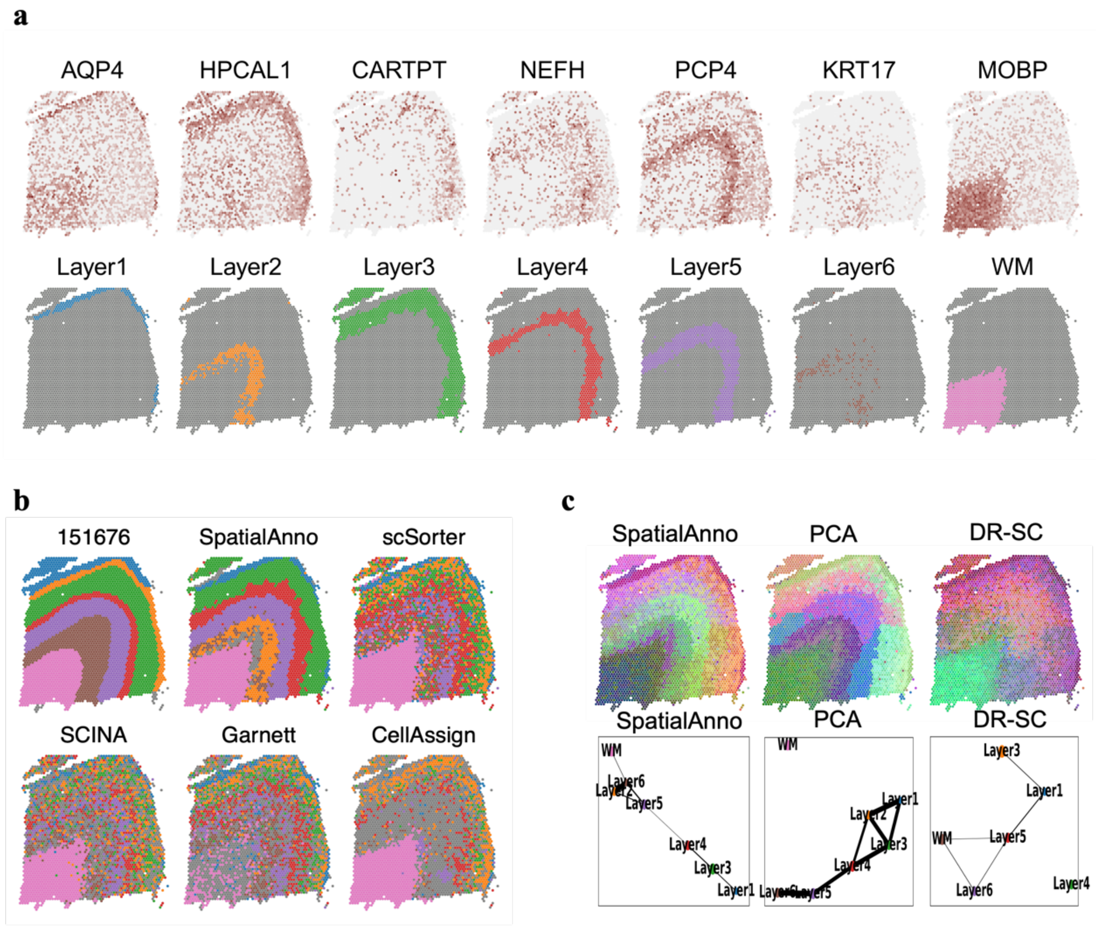

**Supplementary Figure 14. Additional analysis results for the DLPFC 10x Visium dataset**

**a** Boxplots for Kappa, mF1, and ACC show the accuracy of different methods using the top 5, 10, and 15 differentially expressed genes for domain annotation across 12 tissue sections. Two-sided Wilcoxon Rank Sum test was used to pair wisely test the difference between metrices of different methods, and the *p*-value is shown. **b** Boxplots of Kappa, mF1, and ACC show the accuracy of different methods when performing annotation with marker genes identified from section 151607 for samples with ID151669-151672 from Donor 2 that only contained five cortical layers. Correctly- (5) or over-specified (7) cell/domain types are provided in the marker gene list. **c** Clustering results measured by ARI (the higher the better) based on low-dimensional embeddings either from marker genes by PCA or non-marker genes by SpatialAnno, or a combination. Two-sided Wilcoxon Rank Sum test was used to pair wisely test the ARI difference, and the *p*-value is shown.

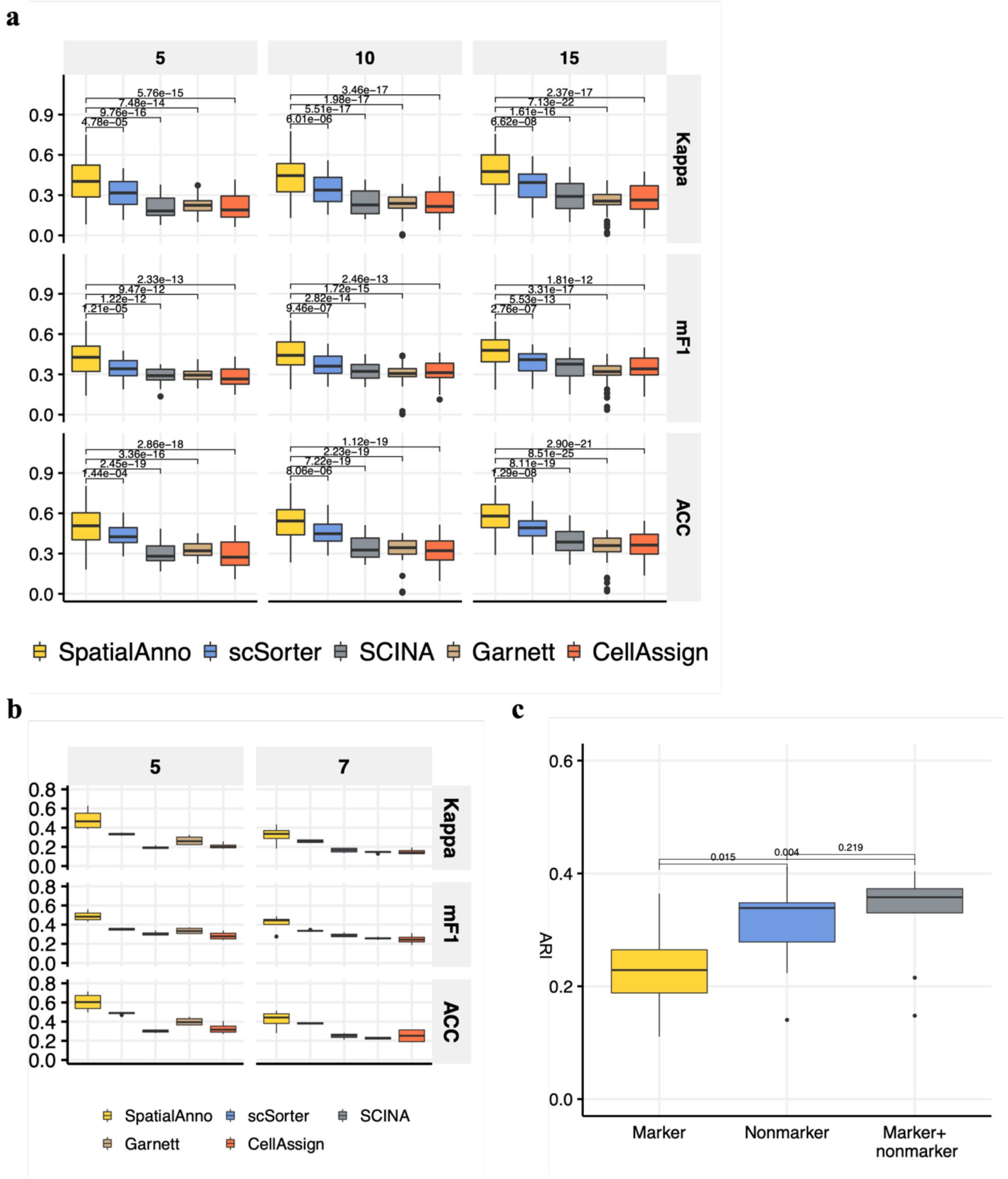

**Supplementary Figure 15. Spatial annotation of mouse olfactory bulb dataset**

**a** Bar plots of Kappa, mF1, and ACC showing the accuracy of the different methods when performing annotation with correctly- (5) or over-specified (7) cell types in the marker gene list. **b** Bar plots of Kappa and mF1 and ACC showing the accuracy of SpatialAnno, scSorter, and Garnett using 30, 300, and 3000 non-marker genes. **C** Bar plots of Kappa, mF1, and ACC showing the accuracy of different methods by manually combining EPL-IN and PGC. **d** Bar plots of Kappa, mF1, and ACC showing the accuracy of different methods by manually combing EPL-IN, M/TC, and PGC. **e** tSNE plot for SpatialAnno, where tSNE PCs were obtained based on the extracted 15-dimensional SpatialAnno embeddings.

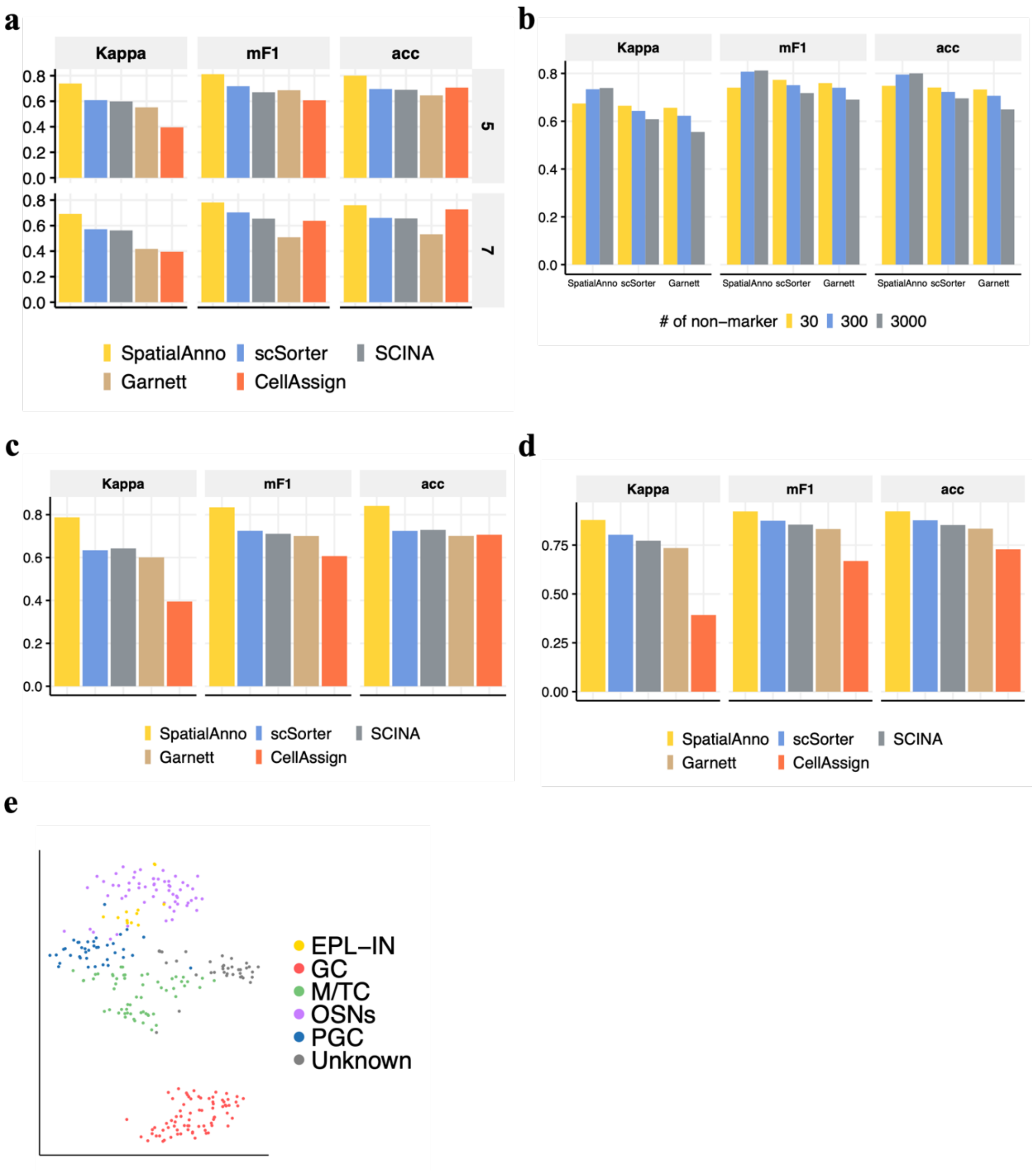

**Supplementary Figure 16. Spatial distribution of cell types in the mouse olfactory bulb dataset across 12 sections annotated via different methods**

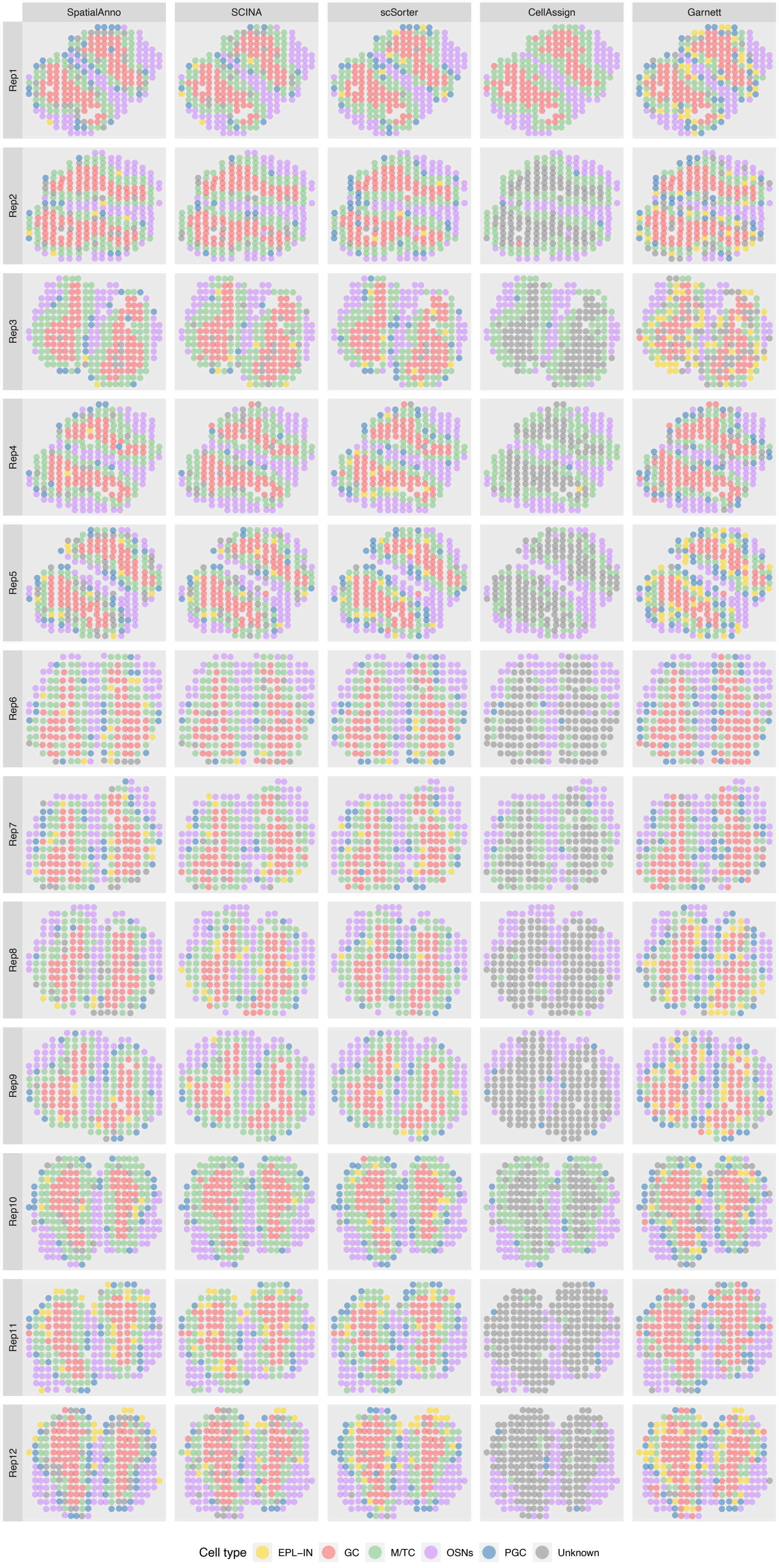

**Supplementary Figure 17. Spatial domain annotations in mouse olfactory bulb section 12 by SpatialAnno, scSorter, SCINA, Garnett, and CellAssign**

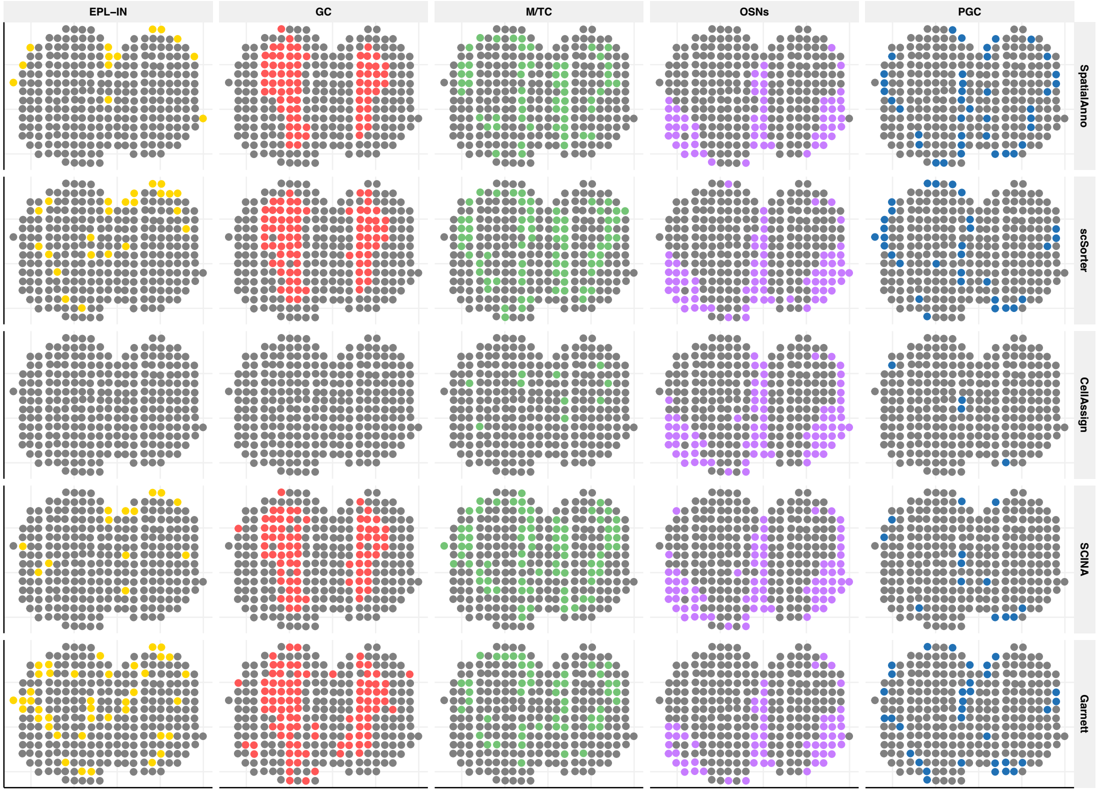

**Supplementary Figure 18. Spatial distribution of cell types in the mouse hippocampus Slide-seqV2 data annotated by different methods**

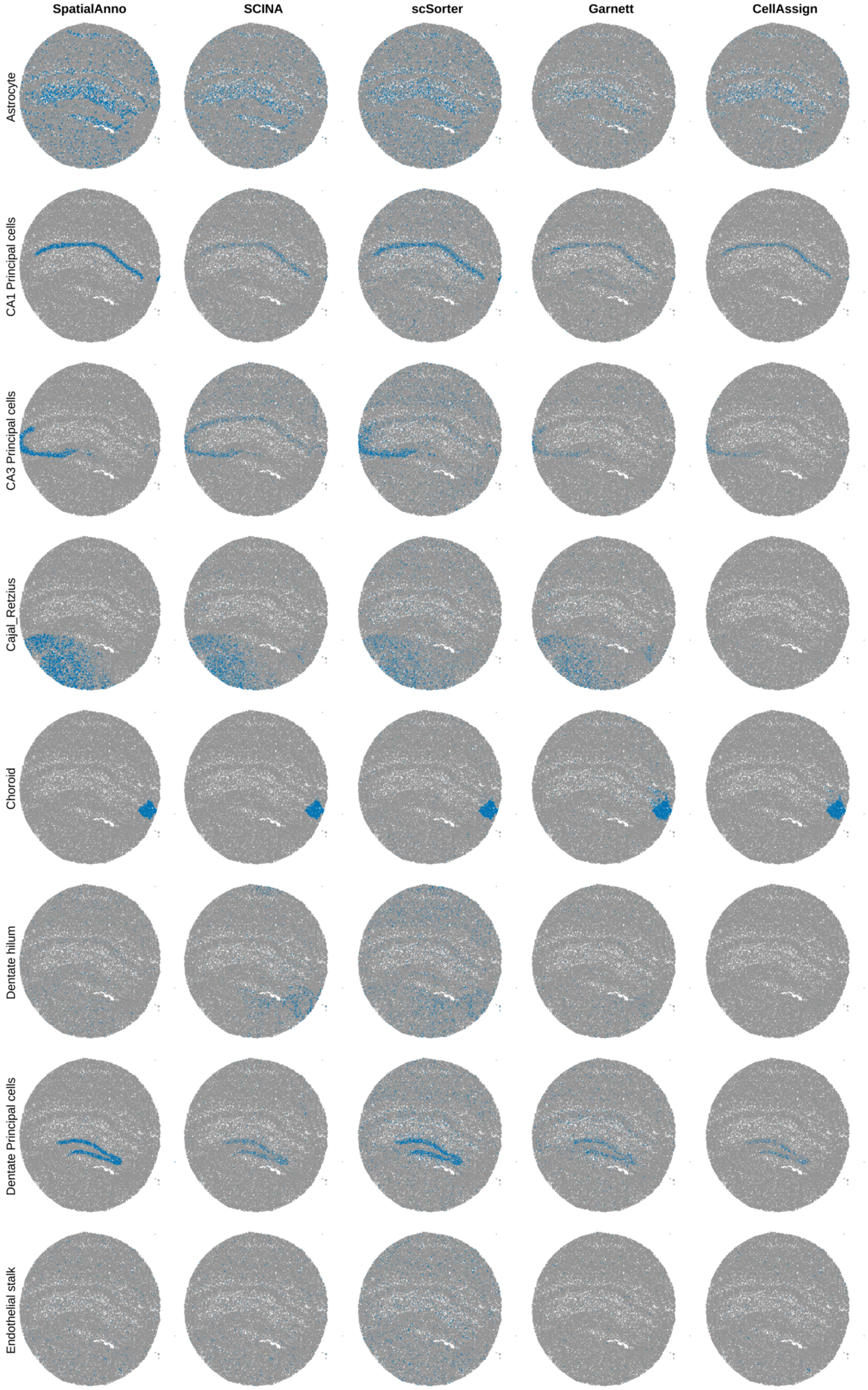

**Supplementary Figure 19. Spatial distribution of cell types in the mouse hippocampus Slide-seqV2 data annotated by different methods (continued)**

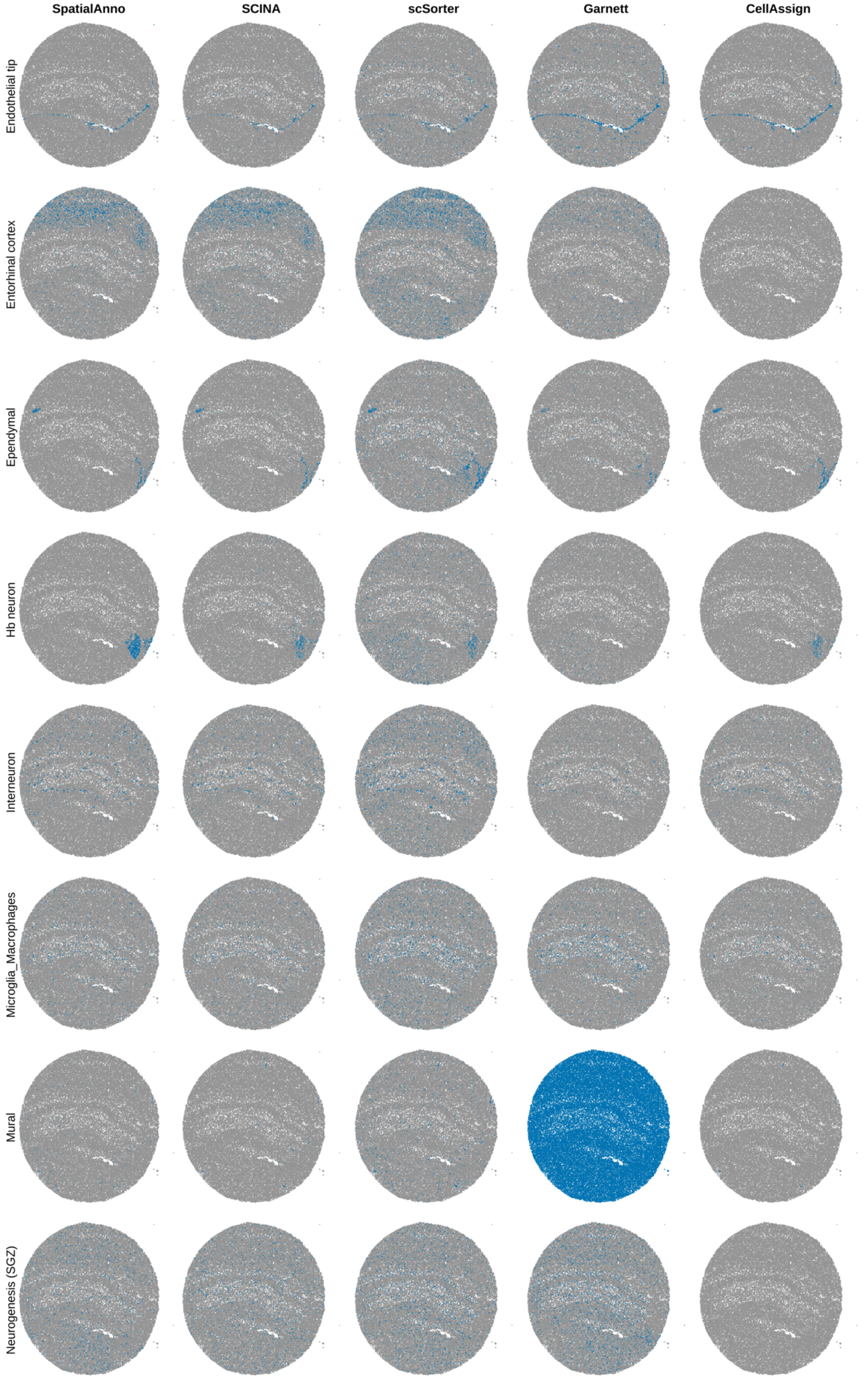

**Supplementary Figure 20. Spatial distribution of cell types in the mouse hippocampus Slide-seqV2 data annotated by different methods (continued)**

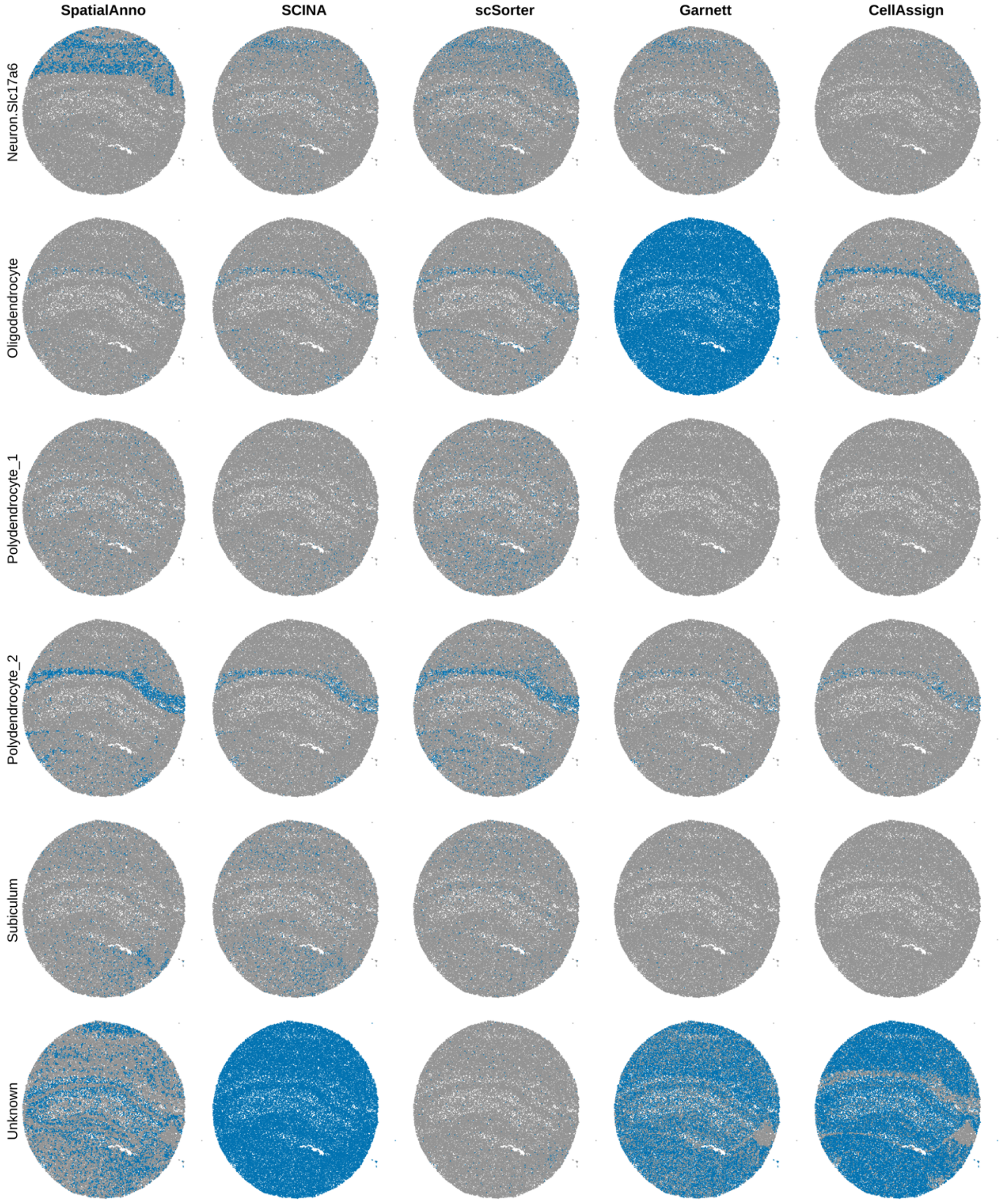

**Supplementary Figure 21. Visualizations and annotations of mouse hippocampus Slide-seqV2 and Slide-seq data**

**a** For SpatialAnno, PCA, and DR-SC, the inferred low-dimensional components of Slide-seqV2 data were summarized into three tSNE components and visualized with RGB plots. **b** Spatial annotation of Slide-seq data for SpatialAnno, scSorter, SCINA, Garnett, and CellAssign**.** **c** Results of Pearson's chi-squared test of correlation between the expression patterns of marker genes and the three hippocampal subfields identified by different methods in Slide-seq data. **d** For SpatialAnno, PCA, and DR-SC, the inferred low-dimensional components of Slide-seq data were summarized into three tSNE components and visualized with RGB plots.

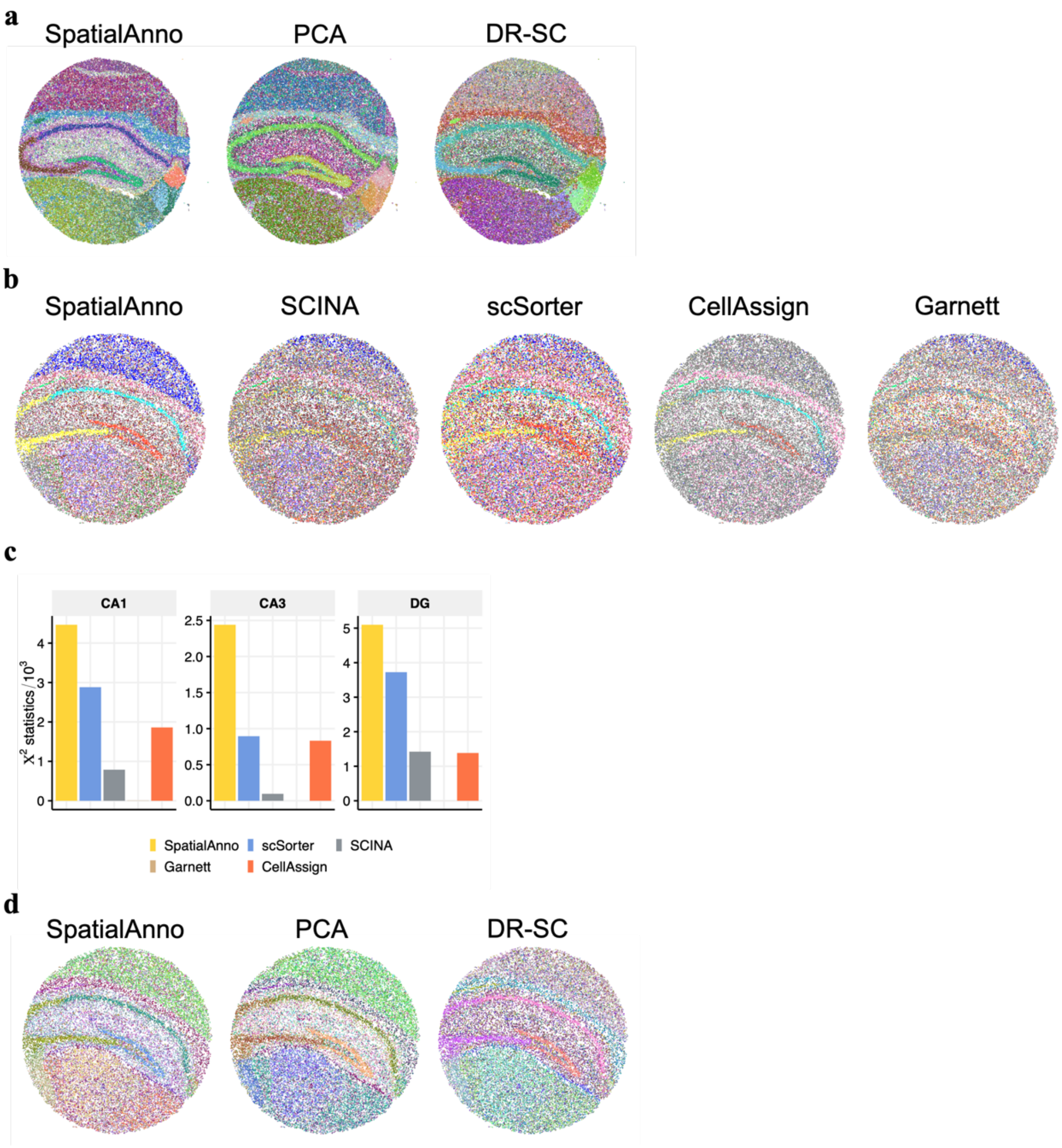

**Supplementary Figure 22. Spatial distribution of cell types in the mouse hippocampus Slide-seqV1 data annotated by different methods**

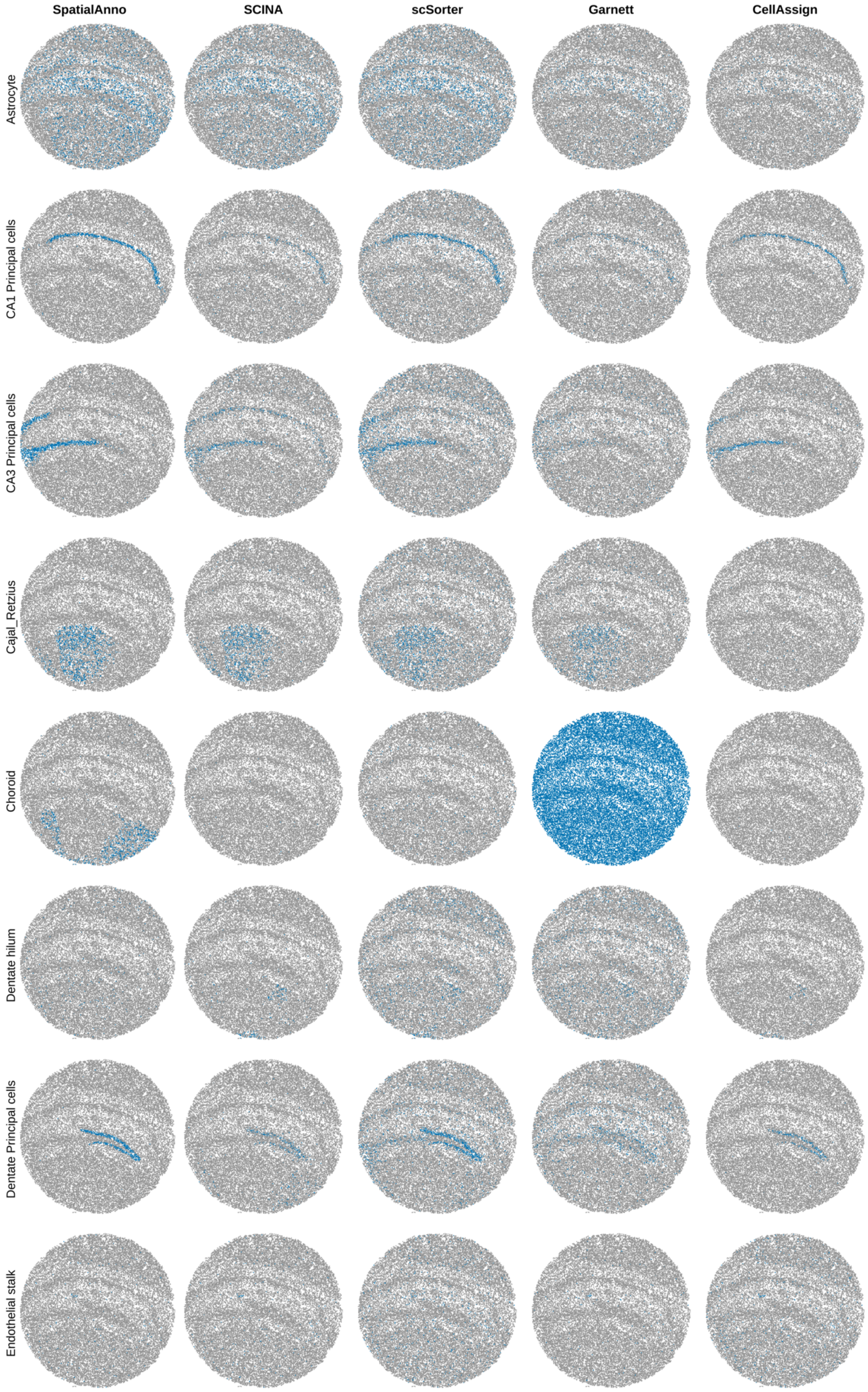

**Supplementary Figure 23. Spatial distribution of cell types in the mouse hippocampus Slide-seqV1 data annotated by different methods (continued)**

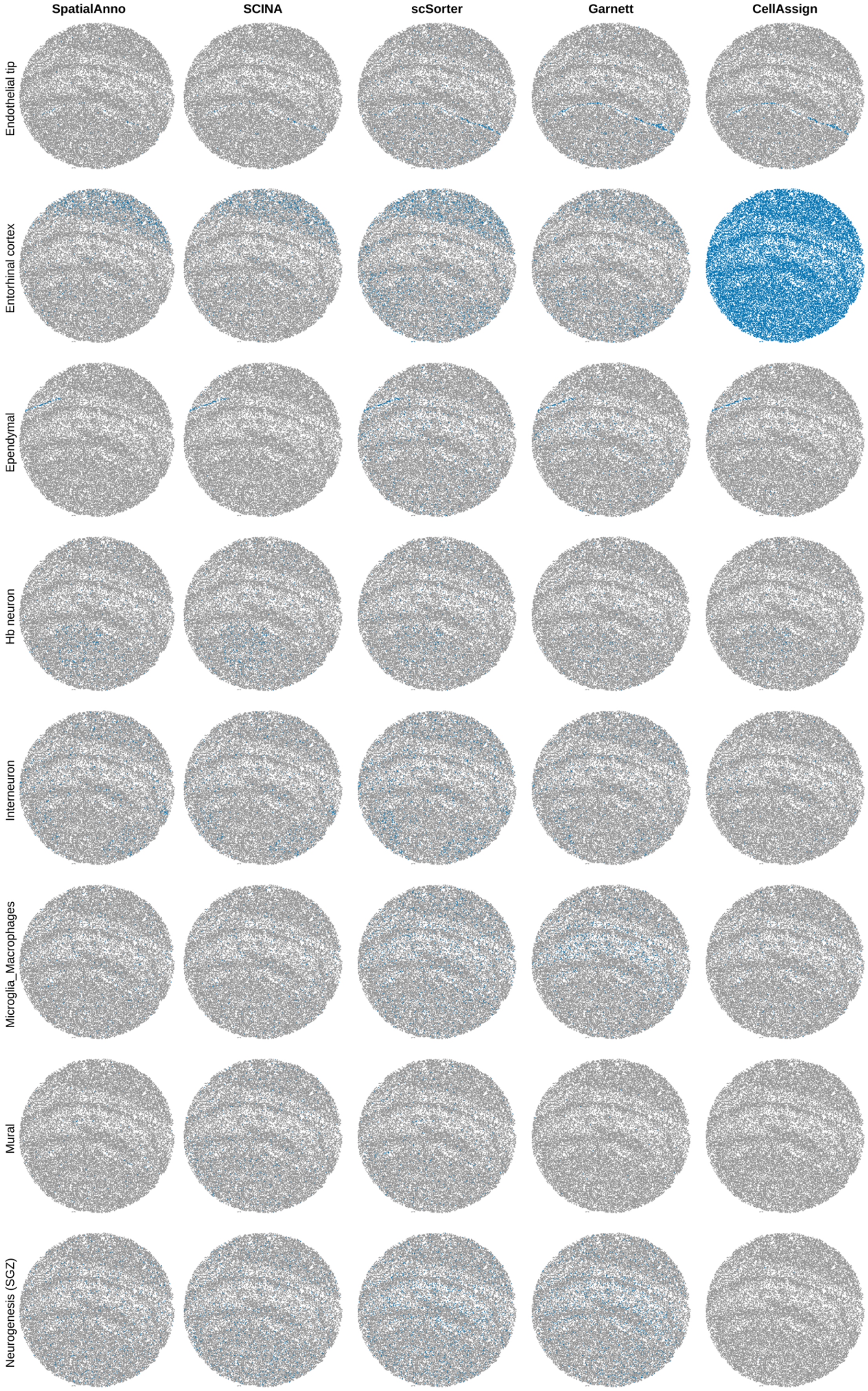

**Supplementary Figure 24. Spatial distribution of cell types in the mouse hippocampus Slide-seqV1 data annotated by different methods (continued)**

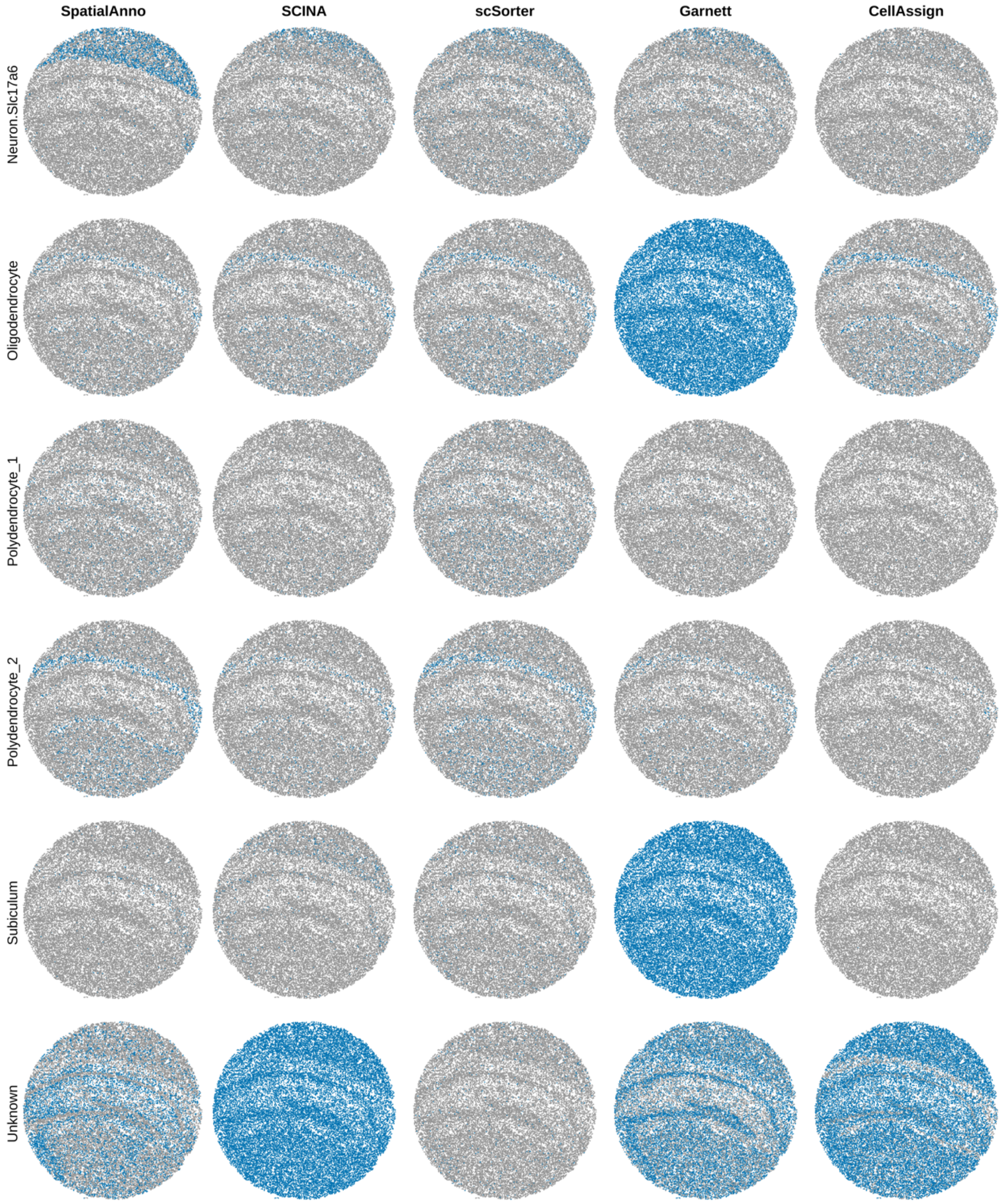

**Supplementary Figure 25. Annotation results for mouse embryo 2 and 3 from seqFISH**

**a** Bar plots of Kappa, mF1, and ACC showing the accuracy of different methods for cell type annotation of embryo 2. **b** Spatial annotations are shown for ground truth, SpatialAnno, scSorter, SCINA, Garnett, and CellAssign in embryo 2. **c** Bar plots of Kappa, mF1 and ACC showing the accuracy of different methods for cell type annotation of embryo 3. **d** Spatial annotations in embryo 3 are shown for ground truth, SpatialAnno, scSorter, SCINA, Garnett, and CellAssign.

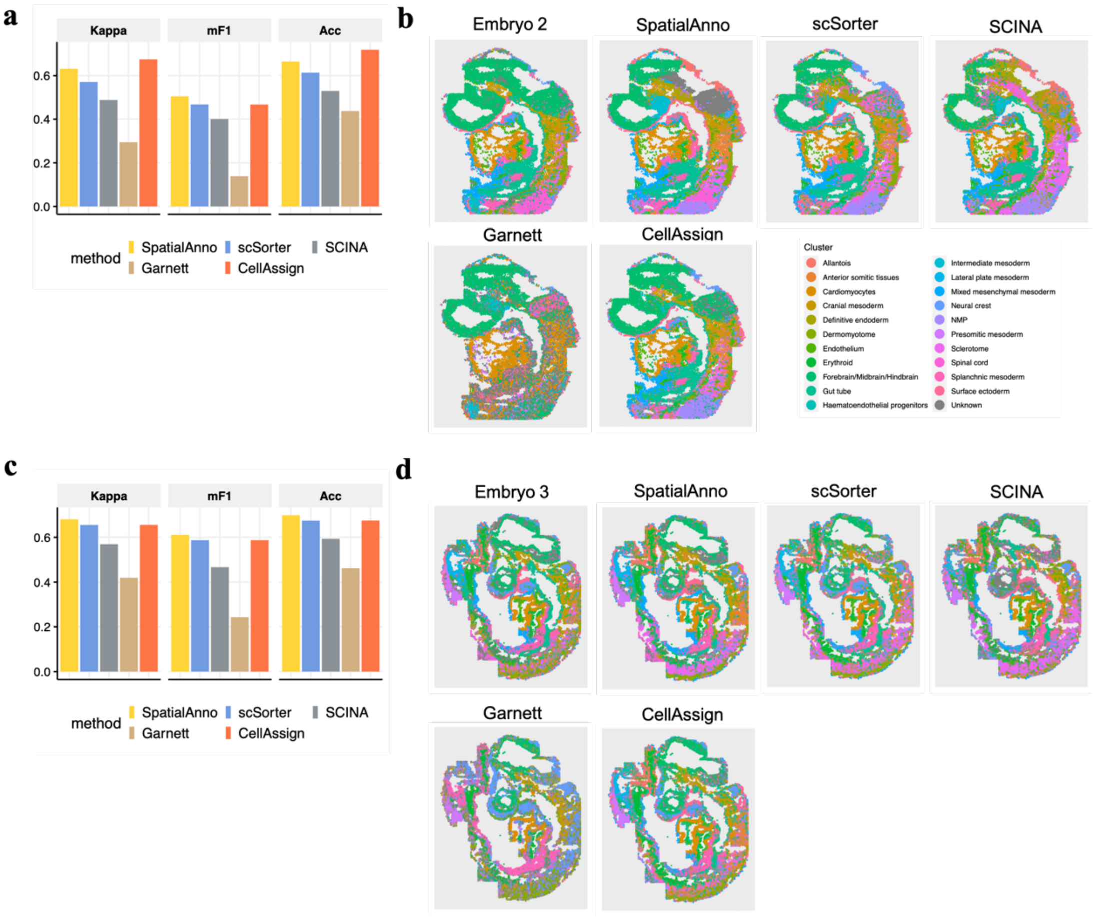

**Supplementary Figure 26. Trajectory inference for mouse embryo 1 data**

**a** Latent time trajectory generated by slingshot on low-dimensional embeddings of PCA. **b** Latent time trajectories generated by slingshot on low-dimensional DR-SC embeddings. **c** Heatmap of gene expression levels for the top 20 genes with significant expression changes with respect to the Slingshot pseudotime. Each column represents a spot that is mapped to this path and is ordered by its pseudotime value. Each row denotes the most significantly changed gene expression. **d** Boxplots of *Otx2* and *Sfrp1* expression counts in selected midbrain and hindbrain regions. **e** Spatial expression of *Otx2* and *Sfrp1* in brain region with corresponding virtual dissection (red line). For each location, only the gene with the higher expression value is plotted.

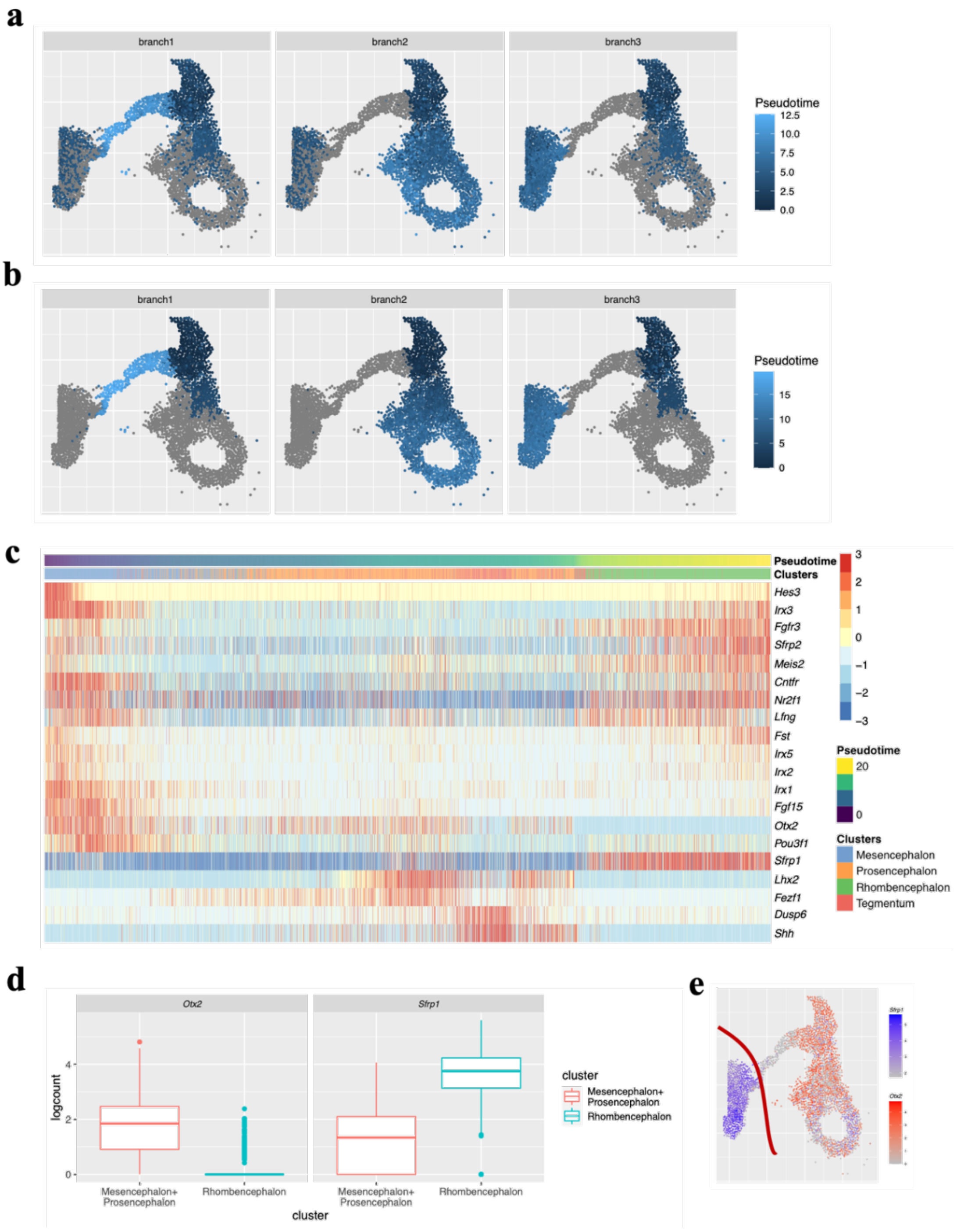

### Supplementary Tables

**Supplementary Table 1. List of 12 DLPFC sections**

| Section ID | Species | Protocol | Year | No. genes | No. spots |
|---|---|---|---|---|---|
| **151507** | Human | 10x Visium | 2021 | 33538 | 4226 |
| **151508** | Human | 10x Visium | 2021 | 33538 | 4384 |
| **151509** | Human | 10x Visium | 2021 | 33538 | 4789 |
| **151510** | Human | 10x Visium | 2021 | 33538 | 4634 |
| **151669** | Human | 10x Visium | 2021 | 33538 | 3661 |
| **151670** | Human | 10x Visium | 2021 | 33538 | 3498 |
| **151671** | Human | 10x Visium | 2021 | 33538 | 4110 |
| **151672** | Human | 10x Visium | 2021 | 33538 | 4015 |
| **151673** | Human | 10x Visium | 2021 | 33538 | 3639 |
| **151674** | Human | 10x Visium | 2021 | 33538 | 3673 |
| **151675** | Human | 10x Visium | 2021 | 33538 | 3592 |
| **151676** | Human | 10x Visium | 2021 | 33538 | 3460 |

#### Supplementary Table 2. List of 12 mouse olfactory bulb (MOB) sections

| Dataset | Species | Protocol | Year | No. genes | No. locations |
|---|---|---|---|---|---|
| **MOB Rep1** | Mouse | ST | 2016 | 16573 | 267 |
| **MOB Rep2** | Mouse | ST | 2016 | 15981 | 280 |
| **MOB Rep3** | Mouse | ST | 2016 | 16014 | 269 |
| **MOB Rep4** | Mouse | ST | 2016 | 15941 | 264 |
| **MOB Rep5** | Mouse | ST | 2016 | 15290 | 267 |
| **MOB Rep6** | Mouse | ST | 2016 | 16251 | 242 |
| **MOB Rep7** | Mouse | ST | 2016 | 16675 | 231 |
| **MOB Rep8** | Mouse | ST | 2016 | 15288 | 234 |
| **MOB Rep9** | Mouse | ST | 2016 | 15284 | 237 |
| **MOB Rep10** | Mouse | ST | 2016 | 16416 | 281 |
| **MOB Rep11** | Mouse | ST | 2016 | 16218 | 262 |
| **MOB Rep12** | Mouse | ST | 2016 | 16034 | 282 |

#### Supplementary Table 3. Features of two mouse hippocampus sections

| Section | Species | Protocol | Year | No. genes | No. locations |
|---|---|---|---|---|---|
| **1** | Mouse | Slide-seq | 2021 | 22457 | 34199 |
| **2** | Mouse | Slide-seqV2 | 2021 | 23264 | 53208 |

#### Supplementary Table 4. Features of three mouse embryo sections

| Section | Specie | Protocol | Year | No. genes | No. locations |
|---|---|---|---|---|---|
| **1** | Mouse | seqFISH | 2021 | 351 | 19451 |
| **2** | Mouse | seqFISH | 2021 | 351 | 14891 |
| **3** | Mouse | seqFISH | 2021 | 351 | 23194 |

**Supplementary Table 5. Computation time for four real data applications**

Computing time in seconds was recorded using a single thread on a 2.1 GHz Intel Xeon Gold 6230 CPU with 16 GB memory

| Method | DLPFC data (average time for 12 sections) | Mouse OB data (average time for 12 sections) | Slide-seq V2 data | seqFISH data (average time for 3 sections) |
|---|---|---|---|---|
| **SpatialAnno** | 137.5 | 2.3 | 5667.6 | 6272.8 |
| **scSorter** | 131.5 | 7.5 | 3262.7 | 1512.5 |
| **SCINA** | 3.8 | 0.1 | 44.9 | 23.3 |
| **Garnett** | 127.9 | 168.9 | 193.1 | 59.5 |
| **CellAssign** | 241.7 | 15.8 | 30608.6 | 16209.7 |

#### Supplementary Table 6. Marker genes used in DLPFC dataset

| Reference section | Type | Top 1-5 markers | Top 6-10 markers | Top 11-15 markers |
|---|---|---|---|---|
| 151507 | Layer 1 | *Gfap,Fabp4,Snorc,Aqp4,Mt1g* | *Fabp7,Saa1,Cxcl14,Vim,Sparc* | *Agt,S100b,Cst3,Gja1,Mlc1* |
| | Layer 2 | *Hpcal1,Enc1,Lamp5,Serpine2,Pcdh8* | *Hopx,Calb1,Atp2b1,Rasgrf2,Fkbp1a* | *Camk2n1,Sowaha,Hpca,Nnat,Itpka* |
| | Layer 3 | *Cartpt,Enc1,Tespa1,Hopx,Hapln4* | *Gria4,Calb1,Nrgn,Pcdh8,Cyp46a1* | *Cbln4,Hs6st1,Itpka,Dbndd1,Yjefn3* |
| | Layer 4 | *Nefh,Nefm,Nefl,Scn1b,Vamp1* | *Parm1,Pvalb,Map1b,Dclk1,Lgals1* | *Fabp3,Hbb,Slc30a3,Gpx3,Tuba1b* |
| | Layer 5 | *Pcp4,Camk2d,Ipcef1,Tmsb10,Hbb* | *Scgb2a2,Smyd2,Efhd2,Tubb2a,Syt1* | *Scgb1d2,Pfkp,Syn2,Diras2,Sncg* |
| | Layer 6 | *Krt17,Diras2,Efhd2,Nptx1,Tbr1* | *Scgb2a2,Scgb1d2,Tf,Scn3b,Mbp* | *Slc24a2,Syt1,Clstn2,Slc17a7,Slc35f1* |
| | WM | *Plp1,Mbp,Mobp,Cnp,Krt8* | *Tf,Mag,Cryab,S100a11,Krt18* | *Ppp1r14a,Septin4,Spp1,Bcas1,Cldnd1* |
| 151508 | Layer 1 | *Gfap,Fabp4,Snorc,Aqp4,Saa1* | *Fabp7,Vim,Agt,Cxcl14,Sparc* | *Mt1g,S100b,Mlc1,Mt1m,Cst3* |
| | Layer 2 | *Hpcal1,Lamp5,Enc1,Pcdh8,Cnr1* | *Serpine2,Hpca,Ncdn,Kcnip2,Gng2* | *Camk2n1,Itpka,Mfsd4a,Ppp3ca,Camkk1* |
| | Layer 3 | *Enc1,Hopx,Rasgrf2,Lmo4,Ca10* | *Cyp46a1,Pcdh8,Tespa1,Rgs4,Nrgn* | *Hpcal1,Ptk2b,Hapln4,Hs6st3,C11orf87* |
| | Layer 4 | *Nefm,Nefh,Hbb,Scrt1,Nefl* | *Vamp1,Scn1b,Pvalb,Dclk1,Gabra1* | *Lynx1,Map1b,Cplx1,Chga,Gars1* |
| | Layer 5 | *Pcp4,Ipcef1,Clstn2,Tmsb10,Tubb2a* | *Stmn2,Scgb2a2,Syt1,Slc24a2,Pfkp* | *Snca,Neurod6,Tuba1b,Efhd2,Diras2* |
| | Layer 6 | *Scgb2a2,Scgb1d2,Agr2,Iglc2,Krt8* | *S100a11,Mbp,Col1a2,Col1a1,Plp1* | *Mobp,Muc1,Diras2,Tf,Rps6* |
| | WM | *Mbp,Plp1,Mobp,Cnp,Tf* | *Bcas1,Cldnd1,Ppp1r14a,Cryab,Spp1* | *Mag,S100a11,Krt8,Paqr6,Lamp2* |
| 151509 | Layer 1 | *Gfap,Aqp4,Saa1,Snorc,Mt1g* | *Vim,Fabp7,Agt,Cxcl14,Fabp4* | *Mt1m,Sparc,Cst3,Mgp,S100b* |
| | Layer 2 | *Hpcal1,Enc1,Egr3,C1ql2,Nptx2* | *Pcdh8,Lamp5,Gng2,Ppp3ca,Cnr1* | *Psd3,Map2k1,Pcdh7,Ncdn,Serpine2* |
| | Layer 3 | *Nefm,Cartpt,Lmo4,Gap43,Rgs4* | *Enc1,Necab1,Ywhah,Nrgn,Nefl* | *Chn1,Btbd8,C11orf87,Rbfox1,Calb1* |
| | Layer 4 | *Nefh,Nefl,Nefm,Dclk1,Scn1b* | *Map1b,Tuba1b,Stmn2,Vamp1,Anxa6* | *Slc24a2,Chga,Tagln3,Spock1,Napb* |
| | Layer 5 | *Pcp4,Igkc,S100a11,Agr2,Krt18* | *Efhd2,Krt8,Scgb1d2,Slc24a2,Tmsb10* | *Iglc2,Diras2,Syt1,Col1a2,Scgb2a2* |
| | Layer 6 | *Col1a2,Krt8,S100a11,S100a10,Fn1* | *Col1a1,Col3a1,Muc1,Agr2,Krt18* | *Ccnd1,Krt19,Scgb1d2,Scgb2a2,Xbp1* |
| | WM | *Mbp,Plp1,Mobp,Tf,Cnp* | *Ppp1r14a,Cryab,Spp1,Cldnd1,Mag* | *Cldn11,Enpp2,Septin4,Rnase1,Mal* |
| 151510 | Layer 1 | *Gfap,Aqp4,Mt1g,Snorc,Sparc* | *Cxcl14,Vim,Agt,Fabp7,Saa1* | *Mgp,Malat1,Cst3,Mt2a,S100b* |
| | Layer 2 | *Hpcal1,Enc1,Pcdh8,Egr3,Lamp5* | *Serpine2,Pcdh7,Nptx2,Sst,Sowaha* | *Hopx,Ppp3ca,Ncdn,Baiap2,Gng2* |
| | Layer 3 | *Cartpt,Nefm,Gap43,Nefl,Enc1* | *Hapln4,Lmo4,Sncg,Ywhah,Ldb2* | *Atp1b1,Chn1,Necab1,Oxr1,Stmn2* |
| | Layer 4 | *Nefh,Nefl,Nefm,Dclk1,Tuba1b* | *Scn1b,Stmn2,Scn1a,Map1b,Ccni* | *Neurod6,Slc6a17,Vgf,Nrep,Ndrg4* |
| | Layer 5 | *Pcp4,Ipcef1,Slc24a2,Gabra5,Tmsb10* | *Tubb2a,Efhd2,Syt1,Clstn2,Camk2d* | *Pfkp,Diras2,Slc35f1,Kcnab1,Tuba1b* |
| | Layer 6 | *Krt8,Krt18,S100a10,Fn1,S100a11* | *Col1a2,Mbp,Col3a1,Krt19,Agr3* | *Col1a1,Plp1,Cldnd1,Slc24a2,Mobp* |
| | WM | *Mbp,Plp1,Mobp,Tf,Cnp* | *Mag,Ppp1r14a,Cldn11,Cryab,Cldnd1* | *Spp1,Bcas1,Enpp2,Rnase1,Mal* |
| 151673 | Layer 1 | *Myl9,Malat1,Mgp,Tagln,Vim* | *Mt1g,Hla-a,Krt19,Cox6c,Atp8* | *Slc1a2,Cst3,Clu,Gfap,Nd5* |
| | Layer 2 | *Hpcal1,Cxcl14,Lamp5,Camk2n1,Serpine2* | *Gnal,Calb2,Hopx,Itpka,Cnr1* | *Enc1,Rasgrf2,Gria2,Pcdh8,Rgs12* |
| | Layer 3 | *Cartpt,Hopx,Enc1,Calb1,Nefm* | *Vstm2a,Lratd1,Gpx3,Hapln4,Cux2* | *Sncg,Nefl,Pcdh8,C11orf87,Ywhah* |
| | Layer 4 | *Nefh,Nefm,Vamp1,Pvalb,Scn1b* | *Rorb,Nefl,Parm1,Ina,Vgf* | *Scn1a,Frmpd2,Dclk1,Gabrb2,Sncg* |
| | Layer 5 | *Pcp4,Tmsb10,Pcp4l1,Smyd2,Gabra5* | *Ipcef1,Syt1,Diras2,Hs3st2,Tubb2a* | *Camk2d,Fam3c,Efhd2,Snap25,Clstn2* |
| | Layer 6 | *Krt17,B3galt2,Scgb1d2,Diras2,Scgb2a2* | *Ifi27,Slc35f1,Tbr1,Krt19,Hla-b* | *Hs3st4,Scn3b,Mmd,Map2k1,Prkcb* |
| | WM | *Plp1,Mbp,Mobp,Cnp,Cryab* | *Tf,Mag,Ppp1r14a,Gfap,Cldn11* | *Ermn,Cldnd1,Spp1,Rnase1,Mog* |
| 151674 | Layer 1 | *Malat1,Myl9,Mgp,Reln,Acta2* | *Cxcl14,Tagln,Cox6c,C11orf96,Csta* | *Bambi,Sparc,Vim,Igfbp7,Krt19* |
| | Layer 2 | *Hpcal1,Serpine2,C1ql2,Cxcl14,Cnr1* | *Gnal,Sowaha,Enc1,Lamp5,Cux2* | *Camk2n1,Hopx,Sst,Calb2,Nptxr* |
| | Layer 3 | *Cartpt,Enc1,Saa1,Hopx,Calb1* | *Fabp4,Ca10,Cux2,Pcdh8,Adcyap1* | *Nefm,Nsg2,Vstm2a,Cbln4,C11orf87* |

| | | | | |
|---|---|---|---|---|
| | Layer 4 | *Nefh,Nefm,Vamp1,Pvalb,Nefl* | *Scn1b,Parm1,Rorb,Syt2,Gpx3* | *Scn1a,Nsg1,Cntnap2,Tpbg,Sncg* |
| | Layer 5 | *Pcp4,Hs3st2,Smyd2,Clstn2,Pcp4l1* | *Camk2d,Tubb2a,Ipcef1,Tmsb10,Nrep* | *Rorb,Nrn1,Vat1l,Fam3c,Syt1* |
| | Layer 6 | *Cpb1,Scgb1d2,Scgb2a2,Krt17,B3galt2* | *Hla-b,Diras2,Krt19,Ifi27,Cox6c* | *Isg15,Bambi,Slc35f1,Tbr1,Scgb2a1* |
| | WM | *Plp1,Mbp,Mobp,Tf,Cnp* | *Mag,Cryab,Gfap,Ppp1r14a,Cldn11* | *Spp1,Ermn,Cldnd1,Bcas1,Carns1* |
| **151675** | Layer 1 | *Azgp1,Myl9,Mgp,Tagln,Malat1* | *Vim,Cxcl14,Cst3,Slc1A2,Apoe* | *Clu,Camk2N1,Mtrnr2L8,Mt3,Nd14* |
| | Layer 2 | *Hpcal1,Cxcl14,Serpine2,Saa1,Fabp4* | *Hopx,Cnr1,Itpka,Camk2N1,Sst* | *Enc1,Lamp5,Linc00507,Rgs12,Cartpt* |
| | Layer 3 | *Cartpt,Hopx,Calb1,Fabp4,Enc1* | *Hs6St3,Saa1,Hapln4,Ca10,Pcdh8* | *Baiap3,Vstm2A,C11orf87,Nsg2,Tespa1* |
| | Layer 4 | *Nefh,Nefm,Pvalb,Rorb,Nefl* | *Scn1B,Parm1,Sncg,Gpx3,Saa1* | *Vamp1,Tpbg,Lgals1,Nsg1,Cabp1* |
| | Layer 5 | *Pcp4,Smyd2,Tmsb10,Pcp4L1,Tubb2A* | *Sncg,Syt1,Camk2D,Syn2,Clstn2* | *Cplx1,Snap25,Nrep,Etv1,Hs3St2* |
| | Layer 6 | *Scgb1D2,Scgb2A2,Krt17,Krt19,B3Galt2* | *Hla-B,Diras2,Tbr1,Scn3B,Slc35F1* | *Rap1Gap2,Map2K1,Hs3St2,Slc17A7,Ly6H* |
| | WM | *Plp1,Mbp,Mobp,Tf,Cnp* | *Cryab,Gfap,Mag,Ppp1R14A,Ermn* | *Cldn11,Cldnd1,Spp1,Carns1,Mog* |
| **151676** | Layer 1 | *Myl9,Tagln,Malat1,Cxcl14,Mt1G* | *Vim,Mgp,Fabp7,Sparc,Snorc* | *Aqp4,Atp8,Mt2A,Cst3,Mt1E* |
| | Layer 2 | *Hpcal1,Cxcl14,Cnr1,Serpine2,Lamp5* | *Hopx,Itpka,Enc1,Sez6L,Camk2N1* | *Linc00507,Tespa1,Mt1G,Necab2,Fkbp1A* |
| | Layer 3 | *Cartpt,Hopx,Enc1,Vstm2A,Calb1* | *Ca10,Cux2,Cbln4,Nefm,Saa1* | *Hapln4,Tespa1,Sncg,Linc01007,Adcyap1* |
| | Layer 4 | *Nefh,Nefm,Pvalb,Saa1,Nefl* | *Scn1B,Vamp1,Sncg,Rorb,Frmpd2* | *Gpx3,Dclk1,Slc30A3,Parm1,Fabp4* |
| | Layer 5 | *Pcp4,Tmsb10,Smyd2,Pcp4L1,Tubb2A* | *Hs3St2,Syn2,Snap25,Pfkp,Syt1* | *Diras2,Clstn2,Rorb,Ndrg4,Cdk14* |
| | Layer 6 | *Scgb1D2,Scgb2A2,Cpb1,Krt17,B3Galt2* | *Diras2,Tff3,Krt19,Slc35F1,Tbr1* | *Hla-B,Bambi,Mmd,Scn3B,Slc17A7* |
| | WM | *Plp1,Mbp,Mobp,Tf,Cnp* | *Gfap,Cryab,Cldnd1,Mag,Ermn* | *Cldn11,Spp1,Mog,Ppp1R14A,Carns1* |

**Supplementary Table 7. Marker genes used in mouse OB dataset**

| Cell type | Markers |
| --- | --- |
| **Granule cells (GC)** | *Gria2, Meis2, Prkca, Penk* |
| **Periglomerular cells (PGC)** | *Nppa, Nrsn1, Nxph1, Th* |
| **Mitral and tufted cell (M/TC)** | *Cdhr1, Slc17a7, Olfm1, Reln* |
| **Olfactory sensory neurons (OSNs)** | *Gng13, S100a5, Omp, Fam213b* |
| **External plexiform layer interneuron (EPL-IN)** | *Kit, Thy1, Dner, Spock2* |
| **Endothelial** | *Ly6c1, Slco1a4, Ly6a, Cldn5* |
| **Mural** | *Cald1, Slco1a4, Ly6c1,Igfbp7* |

**Supplementary Table 8. Marker genes used in hippocampus dataset**

| Cell type | Markers |
|---|---|
| **Entorhinal cortex** | *Mef2c, Nrgn, Vsnl1, Snap25, Meg3* |
| **Ependymal** | *Dbi, Ccdc153, Rarres2, Tmem212, Nnat* |
| **CA3 Principal cells** | *Chgb, Hs3st4, Nptxr, Cpne4, Neurod6* |
| **Dentate hilum** | *Calb2, Pde1a, Rab3c, Satb1, Ajap1* |
| **Subiculum** | *Nov, Dcn, Gap43, Pou3f1, Pde1a* |
| **Interneuron** | *Gad2, Gad1, Cnr1, Slc6a1, Nrxn3* |
| **CA1 Principal cells** | *Wfs1, Fibcd1, Atp2b1, Itpka, Ppp3ca* |
| **Oligodendrocyte** | *Plp1, Ptgds, Mbp, Mal, Mag* |
| **Astrocyte** | *Apoe, Cst3, Aldoc, Mt1, Clu* |
| **Endothelial stalk** | *Ly6c1, Bsg, Flt1, Itm2a, Ly6a* |
| **Endothelial tip** | *Ptgds, Apod, Igf2, Igfbp2, Col1a2* |
| **Polydendrocyte_2** | *Mbp, Sirt2, Plp1, Cnp, Mag* |
| **Polydendrocyte_1** | *Olig1, Cacng4, Ptprz1, Marcks, Lhfpl3* |
| **Mural** | *Acta2, Tpm1, Tpm2, Crip1, Tagln* |
| **Dentate Principal cells** | *C1ql2, Ppp3ca, Fam163b, Olfm1, Ncdn* |
| **Neurogenesis (SGZ)** | *Sox4, Tubb5, Sox11, Rps5, Rps9* |
| **Hb neurons** | *Rora, Slc17a7* |
| **Choroid** | *Ttr, Enpp2,1500015o10rik, Prlr, Igfbp2* |
| **Cajal_Retzius** | *Pcp4, Ndnf, Ramp1, Gap43, Reln* |
| **Neuron.Slc17a6** | *Camk2n1, Nrgn, Cck, Tshz2, Arpp21* |
| **Microglia_Macrophages** | *Hexb, Cst3, Ctss, Cx3cr1, C1qb* |

##### Supplementary Table 9. Marker genes used in embryo dataset

| Cell type | Markers |
|---|---|
| **Lateral plate mesoderm** | *Pitx1, Cdx4, Evx1, Cdx2, Hand2, Hoxa9, Hoxc6, Hoxc9* |
| **Erythroid** | *Slc4a1, Alas2, Klf1, Gata1, Epor, Acp5, Hemgn, Smim1* |
| **Allantois** | *Tbx4, Pitx1, Hand1, Wnt2, Col1a1, Hand2, Msx1, Tbx3* |
| **Gut tube** | *Cdh1, Cldn4, Foxa1, Shh, Cpn1, Krt18, Nepn, Clic6* |
| **Endothelium** | *Cdh5, Plvap, Eng, Cd34, Cldn5, Kdr, Sox18, Pecam1* |
| **Hematoendothelial progenitors** | *Cldn5, Sox18, Cdh5, Plvap, Cd34, Eng, Kdr, Sox7* |
| **Intermediate mesoderm** | *Lef1, Pitx1, Evx1, Tbx3, Tbx4, Bmp4, Cdx2, Cdx4* |
| **Mixed mesenchymal mesoderm** | *Hand1, Col1a1, Tmem108, Ahnak, Dlk1, Gata6, Postn, Smoc2* |
| **Spinal cord** | *Hoxb9, Hoxd4, Sox2, Hoxb8, Foxb1, Hoxc6, Hoxc8, Foxa2* |
| **Neural crest** | *Sox10, Tfap2b, Tfap2a, Nr2f1, Prrx1, Snai1, Msx1, Alx1* |
| **Splanchnic mesoderm** | *Foxf1, Osr1, Hoxb1, Isl1, Gata5, Tbx5, Kcng1, Gata4* |
| **Forebrain/Midbrain/Hindbrain** | *Sfrp1, Sox2, Otx2, Lfng, Nr2f1, Cdh2, Ptn, Cntfr* |
| **Cranial mesoderm** | *Tbx1, Col26a1, Foxc2, Marcks, Tmem119, Cxcl12, Col1a2, Fst* |
| **Surface ectoderm** | *Pdgfa, Tfap2a, Cdh1, Epcam, Cldn4, Itga3, Krt18, Gjb3* |
| **Definitive endoderm** | *Shh, Foxa1, Foxa2, Cdh2, T, Cldn4, Irx1, Irx3* |
| **NMP** | *Hoxb9, Hoxc8, Hoxd4, Sox2, Hoxb4, Hoxc6, Foxb1, Hoxb8* |
| **Anterior somitic tissues** | *Meox1, Foxc2, Col26a1, Cxcl12, Snai1, Marcks, Aldh1a2, Fst* |
| **Presomitic mesoderm** | *Meox1, Dll3, Foxc2, Dll1, Lef1, Cer1, Notch1, Mesp2* |
| **Dermomyotome** | *Meox1, Aldh1a2, Six1, Hoxb3, Col26a1, Foxc2, Hoxb4, Fst* |
| **Cardiomyocytes** | *Popdc2, Atp1b1, Tagln, Ttn, Smarcd3, Gata5, Tbx5, Hcn4* |
| **Sclerotome** | *Meox1, Aldh1a2, Foxc2, Col26a1, Cxcl12, Marcks, Pdgfra, Snai1* |
