## Supplementary Notes for "Probabilistic cell/domain-type assignment of spatial transcriptomics data with SpatialAnno"

#### 1 Details on evaluations in both simulations and real data sets

Cohen's kappa value (Kappa) is generally thought to be a more robust measure than classification accuracy (ACC), as it takes into account the possibility of the agreement occurring by chance. The definition of Kappa is

$$\text{Kappa} = \frac{\text{ACC} - p_e}{1 - p_e},$$

where $p_e$ is the hypothetical probability of chance agreement. For $K$ cell types, $N$ cells to annotate, and $n_k$ is the number of cells belonging to cell type $k$ in the ground truth:

$$p_e = \frac{1}{N^2} \sum a_k n_k,$$

where $a_k$ is the number of the $k$-th cell type assigned by some annotation method. A Kappa of 1 implies the annotation results are in complete agreement with the ground truth, whereas 0 implies that the annotation is no better than a random guess.

† The first two authors have contributed equally to this work.

mF1 is the average of F1 scores for different cell types. The cell-type-level F1 score considers each cell as an individual classification task with a true cell-type assignment. We calculated the F1 score as follows:

$$\text{F1 score} = 2 \times \frac{\text{precision} \times \text{recall}}{\text{precision} + \text{recall}},$$

where precision is the number of true positive samples divided by the number of all samples identified as this cell type, and recall is the number of true positive samples divided by the number of samples in this cell type.

### 2 Methodological details of SpatialAnno

The SpatialAnno model is defined by equations (1)-(3) in the main text, with the details described in the Methods section.

Statistical inference for SpatialAnno is done using a restricted expectation-maximization algorithm[4] with an iterative conditional mode[2]. The latent variables are $(\boldsymbol{y}, \boldsymbol{z})$, and the model parameters to be maximized are $\boldsymbol{m} = \{\boldsymbol{m}_k\}$, $V, L, \Lambda$, $\boldsymbol{\alpha} = \{\alpha_j\}$, $\boldsymbol{\beta} = \{\beta_{jk}\}$, $\boldsymbol{\sigma} = \{\sigma_j\}$, and $\xi$. To facilitate description, we denote $\boldsymbol{\theta}_1 = \{\boldsymbol{\alpha}, \boldsymbol{\beta}, \boldsymbol{\sigma}\}$ as the parameters in equation (1) from the main text and $\boldsymbol{\theta}_2 = \{\boldsymbol{m}, V, L, \Lambda\}$ as the parameters in equation (2) from the main text.

The complete data likelihood $p(\boldsymbol{x}, \boldsymbol{y}, \boldsymbol{z}) = p(\boldsymbol{x}, \boldsymbol{z}, \mid \boldsymbol{y}) p(\boldsymbol{y})$ is difficult to deal with because of the complicated form of $p(\boldsymbol{y})$. To overcome the bottleneck, one of the most used approaches is pseudo likelihood[1], which approximates the joint distribution of $\boldsymbol{y}$ as the product of the full-conditional distribution for each $y_i$. Therefore, the pseudo log-likelihood function of the complete-data is

$$\begin{aligned}
\log \tilde{p}(\boldsymbol{x}, \boldsymbol{y}, \boldsymbol{z}) &= \log p(\boldsymbol{x}, \boldsymbol{z} \mid \boldsymbol{y}) + \log \tilde{p}(\boldsymbol{y}) \\
&= \sum_i \log p(\boldsymbol{x}_{i1}, \boldsymbol{x}_{i2}, \boldsymbol{z}_i \mid y_i) + \sum_i \log p(y_i \mid y_{N_i}) \\
&= \sum_i \log p(\boldsymbol{x}_{i1} \mid y_i) + \sum_i \log p(\boldsymbol{x}_{i2}, \boldsymbol{z}_i \mid y_i) + \sum_i \log p(y_i \mid y_{N_i}) \\
&= \sum_i \log p(\boldsymbol{x}_{i1}, \boldsymbol{x}_{i2}, \boldsymbol{z}_i, y_i \mid \boldsymbol{y}_{N_i}).
\end{aligned} \tag{1}$$

Starting at some initial values for $\boldsymbol{y}$ and parameters $\boldsymbol{\theta} = \{\boldsymbol{\theta}_1, \boldsymbol{\theta}_2, \xi\}$, the algorithm iterates through an iterative conditional mode (ICM) step, expectation step (E-step), and maximization step (M-step) to maximize the pseudo log-likelihood. The initial values could be provided by a non-spatial annotation method. For transcriptome-wide SRT data (such as ST and Visium), we relied on SCINA[5] because of its computational efficiency. For SRT data with limited multiplexing capacity (such as from seqFISH), scSorter[3] can also be used. Alternative initialization can also be supplied by users. We describe the detailed optimization algorithm for each step in the following subsections.

#### 2.1 ICM-step: updating $\boldsymbol{y}$

Based on the Potts model, we know

$$\log p(y_i \mid \boldsymbol{y}_{N_i}) = -\log C_\xi(\boldsymbol{y}_{N_i}) - \xi \sum_{i' \in N_i} \left[1 - \mathrm{I}(y_i = y_{i'})\right],$$

where $C_\xi(\boldsymbol{y}_{N_i})$ is a normalization constant with respect to $\xi$ and $\boldsymbol{y}_{N_i}$.

Based on the properties of the normal distribution, we know $\boldsymbol{x}_{i2} \mid y_i = k$ is a normal, and its mean and variance can be derived from the law of total expectation:

$$\begin{aligned}
\mathrm{E}(\boldsymbol{x}_{i2} \mid y_i = k) &= \mathrm{E}\left[\mathrm{E}(\boldsymbol{x}_{i2} \mid y_i = k, \boldsymbol{z}_i) \mid y_i = k\right] \\
&= \mathrm{E}\left[\mathrm{E}(\boldsymbol{x}_{i2} \mid \boldsymbol{z}_i) \mid y_i = k\right] \\
&= \mathrm{E}(L\boldsymbol{z}_i \mid y_i = k) \\
&= L\boldsymbol{\mu}_k, \\
\mathrm{Var}(\boldsymbol{x}_{i2} \mid y_i = k) &= \mathrm{E}\left[\mathrm{Var}(\boldsymbol{x}_i \mid \boldsymbol{z}_i) \mid y_i = k\right] + \mathrm{Var}\left[\mathrm{E}(\boldsymbol{x}_i \mid \boldsymbol{z}_i) \mid y_i = k\right] \\
&= \mathrm{E}\left(\Lambda \mid y_i = k\right) + \mathrm{Var}\left(L\boldsymbol{z}_i \mid y_i = k\right) \\
&= \Lambda + LVL^\top.
\end{aligned} \tag{2}$$

We have

$$\log p(\boldsymbol{x}_{i2} \mid y_i = k) = -\frac{p}{2}\log 2\pi + \frac{1}{2}\log|S| - \frac{1}{2}(\boldsymbol{x}_{i2} - L\boldsymbol{\mu}_k)^\top S(\boldsymbol{x}_{i2} - L\boldsymbol{\mu}_k),$$

where $S = (\Lambda + LVL^\top)^{-1}$. To calculate $S$ efficiently, we can use the Woodbury formula:

$$S = \Lambda^{-1} - \Lambda^{-1}LML^\top\Lambda^{-1}.$$

where $M = (V^{-1} + L^\top\Lambda^{-1}L)^{-1}$.

In the ICM step, the estimate of $\boldsymbol{y}$ is obtained by maximizing its posterior with respect to $y_i$ coordinately:

$$p(\boldsymbol{y} \mid \boldsymbol{x}) = p(y_i, \boldsymbol{y}_{-i} \mid \boldsymbol{x}) = p(y_i \mid \boldsymbol{x}, \boldsymbol{y}_{-i})p(\boldsymbol{y}_{-i} \mid \boldsymbol{x}),$$

where $i = 1, \ldots, n$, until converge. As the second term in the last equation does not depend on $y_i$, and

$$p(y_i \mid \boldsymbol{x}, \boldsymbol{y}_{-i}) \propto p(\boldsymbol{x}_i \mid y_i)p(y_i \mid \boldsymbol{y}_{N_i} = \hat{\boldsymbol{y}}_{N_i}),$$

we have

$$\begin{aligned}
\hat{y}_i^u =& \arg\max_{y_i} \left[\log p(\boldsymbol{x}_i \mid y_i) + \log p(y_i \mid \boldsymbol{y}_{N_i} = \hat{\boldsymbol{y}}_{N_i})\right] \\
=& \arg\max_{y_i} \left[\log p(\boldsymbol{x}_{1i} \mid y_i) + \log p(\boldsymbol{x}_{2i} \mid y_i) + \log p(y_i \mid \boldsymbol{y}_{N_i} = \hat{\boldsymbol{y}}_{N_i})\right] \\
=& \arg\min_{k} \left\{ \sum_{j=1}^{m} \left[\frac{1}{2}\log 2\pi + \frac{1}{2}\log\sigma_j^2 + \frac{1}{2\sigma_j^2}(x_{ij} - \alpha_j - \rho_{jk}\beta_{jk})^2\right] \right. \\
& + \frac{p}{2}\log 2\pi - \frac{1}{2}\log|S| + \frac{1}{2}(\boldsymbol{x}_{i2} - L\boldsymbol{\mu}_k)^\top S(\boldsymbol{x}_{i2} - L\boldsymbol{\mu}_k) \\
& \left. + \log C_\xi(\hat{\boldsymbol{y}}_{N_i}) + \xi \sum_{i' \in N_i} \left[1 - \mathrm{I}(\hat{y}_{i'} = k)\right] \right\}.
\end{aligned} \tag{3}$$

#### 2.2 E-step: updating responsibility

Now, we define the responsibility that component $k$ takes for explaining the observation $\boldsymbol{x}_i$ as

$$\begin{aligned}\gamma_{ik} =& p(y_i = k \mid \boldsymbol{x}_i, \boldsymbol{y}_{N_i} = \hat{\boldsymbol{y}}_{N_i}) \\ =& \frac{p(\boldsymbol{x}_i \mid y_i = k)p(y_i = k \mid \boldsymbol{y}_{N_i} = \hat{\boldsymbol{y}}_{N_i})}{\sum_{k'} p(\boldsymbol{x}_i \mid y_i = k')p(y_i = k') \mid \boldsymbol{y}_{N_i} = \hat{\boldsymbol{y}}_{N_i})} \\ =& \frac{p(\boldsymbol{x}_{i1} \mid y_i = k)p(\boldsymbol{x}_{i2} \mid y_i = k)p(y_i = k \mid \boldsymbol{y}_{N_i} = \hat{\boldsymbol{y}}_{N_i})}{\sum_{k'} p(\boldsymbol{x}_{i1} \mid y_i = k')p(\boldsymbol{x}_{i2} \mid y_i = k')p(y_i = k') \mid \boldsymbol{y}_{N_i} = \hat{\boldsymbol{y}}_{N_i})}.\end{aligned} \tag{4}$$

It is easy to show that the optimal posterior distribution of $(y_i = k, \boldsymbol{z}_i)$ is

$$\begin{aligned}p(y_i = k, \boldsymbol{z}_i \mid \boldsymbol{x}_i, \boldsymbol{y}_{N_i} = \hat{\boldsymbol{y}}_{N_i}) &= p(y_i = k \mid \boldsymbol{x}_i, \boldsymbol{y}_{N_i} = \hat{\boldsymbol{y}}_{N_i})p(\boldsymbol{z}_i \mid \boldsymbol{x}_i, y_i = k) \\ &= \gamma_{ik}\mathcal{N}(\boldsymbol{z}_i \mid \boldsymbol{w}_{ik}, M),\end{aligned} \tag{5}$$

where

$$\begin{aligned}\boldsymbol{w}_{ik} &= M\left[V^{-1}\boldsymbol{m}_k + L^\top \Lambda^{-1}\boldsymbol{x}_{i2}\right], \\ M &= \left[V^{-1} + L^\top \Lambda^{-1} L\right]^{-1}.\end{aligned} \tag{6}$$

Note that the conditional expectation of $\boldsymbol{z}_i$ given $(\boldsymbol{x}_i, \boldsymbol{y}_{N_i} = \hat{\boldsymbol{y}}_{N_i})$ provides a low-dimensional embedding for spot $i$:

$$\mathrm{E}(\boldsymbol{z}_i \mid \boldsymbol{x}_i, \boldsymbol{y}_{N_i} = \hat{\boldsymbol{y}}_{N_i}) = \sum_k \gamma_{ik}\boldsymbol{w}_{ik}.$$

Obviously, the embedding is label-relevant due to $\gamma_{ik}$.

#### 2.3 Restricted M-step

Taking expectation of (1) w.r.t. the posterior distribution of $(y_i, \boldsymbol{z}_i)$ in (5), the $Q$ function is

$$\begin{aligned}Q(\boldsymbol{\theta}) &= \sum_i \mathrm{E}_{q_i} \log\left[p(\boldsymbol{x}_i, \boldsymbol{z}_i, y_i \mid \boldsymbol{y}_{N_i} = \hat{\boldsymbol{y}}_{N_i})\right] \\ =& \sum_i\sum_k \int \log\left[p(\boldsymbol{x}_i, \boldsymbol{z}_i, y_i = k \mid \boldsymbol{y}_{N_i} = \hat{\boldsymbol{y}}_{N_i})\right] p(y_i = k, \boldsymbol{z}_i \mid \boldsymbol{x}_i, \boldsymbol{y}_{N_i} = \hat{\boldsymbol{y}}_{N_i}) d\boldsymbol{z}_i \\ =& \sum_i\sum_k \int \gamma_{ik}\left[\log p(\boldsymbol{x}_i, \boldsymbol{z}_i \mid y_i = k) + \log p(y_i = k \mid \boldsymbol{y}_{N_i} = \hat{\boldsymbol{y}}_{N_i})\right]\mathcal{N}(\boldsymbol{w}_{ik}, M) d\boldsymbol{z}_i \\ =& \sum_i\sum_k \int \gamma_{ik}\left[\log p(\boldsymbol{x}_i \mid \boldsymbol{z}_i) + \log p(\boldsymbol{z}_i \mid y_i = k) + \log p(y_i = k \mid \boldsymbol{y}_{N_i} = \hat{\boldsymbol{y}}_{N_i})\right]\mathcal{N}(\boldsymbol{w}_{ik}, M) d\boldsymbol{z}_i \\ =& \sum_i\sum_{j=1}^m\sum_k \gamma_{ik}\left[-\frac{1}{2}\log 2\pi - \frac{1}{2}\log\sigma_j^2 - \frac{1}{2\sigma_j^2}(x_{ij} - \alpha_j - \rho_{jk}\beta_{jk})^2\right] \\ &+ \sum_i\sum_k \gamma_{ik}\left[-\frac{p}{2}\log 2\pi - \frac{1}{2}\log|\Lambda| - \frac{1}{2}(\boldsymbol{x}_{2i} - L\boldsymbol{w}_{ik})^\top\Lambda^{-1}(\boldsymbol{x}_{2i} - L\boldsymbol{w}_{ik}) - \frac{1}{2}\mathrm{Tr}(\Lambda^{-1}LML^\top)\right] \\ &+ \sum_i\sum_k \gamma_{ik}\left[-\frac{q}{2}\log 2\pi - \frac{1}{2}\log|V| - \frac{1}{2}(\boldsymbol{w}_{ik} - \boldsymbol{m}_k)^\top V^{-1}(\boldsymbol{w}_{ik} - \boldsymbol{m}_k) - \frac{1}{2}\mathrm{Tr}(V^{-1}M)\right] \\ &- \sum_i \log C_i(\xi, \hat{\boldsymbol{y}}_{N_i}) - \xi\sum_i\sum_k\gamma_{ik}\sum_{i'\in N_i}\left[1 - \mathrm{I}(y_{i'} = k)\right],\end{aligned} \tag{7}$$

##### 2.3.1 Updating $\boldsymbol{\theta}_1$

The parameter $\beta_{jk} \geq 0$ corresponds to the average fold-change in the expression of gene $j$ overexpressed in type $k$. In real data applications, where well-understood marker genes exist, a *priori* information about their expression levels in corresponding cell types may be available. To incorporate such information, other lower boundaries can be imposed.

As there is a linear inequality constraining $\beta_{jk} \geq 0$, we performed constrained optimization via the following Lagarangian

$$Q(\boldsymbol{\theta}) + \sum_j \sum_k \eta_{jk}\rho_{jk}\beta_{jk}.$$

The Karush-Kuhn-Tucker (KKT) conditions were

$$\begin{aligned}
\rho_{jk}\beta_{jk} &\geq 0, \\
\eta_{jk} &\geq 0, \\
\eta_{jk}\rho_{jk}\beta_{jk} &= 0, \\
\frac{\partial Q}{\partial \beta_{jk}} &= \sum_i \frac{\gamma_{ik}}{\sigma_j^2}(x_{ij} - \alpha_j - \rho_{jk}\beta_{jk})\rho_{jk} + \eta_{jk}\rho_{jk} = 0 \text{ for } \rho_{jk} = 1, \\
\frac{\partial Q}{\partial \alpha_j} &= \sum_i \sum_k \frac{\gamma_{ik}}{\sigma_j^2}(x_{ij} - \alpha_j - \rho_{jk}\beta_{jk}) = 0, \\
\frac{\partial Q}{\partial \sigma_j^2} &= \sum_i \sum_k \gamma_{ik}\left[-\frac{1}{2\sigma_j^2} + \frac{1}{2\sigma_j^4}(x_{ij} - \alpha_j - \rho_{jk}\beta_{jk})^2\right].
\end{aligned} \tag{8}$$

Examining the KKT conditions will show the final solution to be

$$\begin{aligned}
\alpha_j =& \frac{1}{n}\sum_i \sum_k \gamma_{ik}(x_{ij} - \rho_{jk}\beta_{jk}), \\
\beta_{jk} =& \max\left\{\frac{\sum_i \gamma_{ik}(x_{ij} - \alpha_j)}{\sum_i \gamma_{ik}}, 0\right\} \text{ for } \rho_{jk} = 1, \\
\sigma_j^2 =& \frac{1}{n}\sum_i \sum_k \gamma_{ik}(x_{ij} - \alpha_j - \rho_{jk}\beta_{jk})^2.
\end{aligned} \tag{9}$$

##### 2.3.2 Updating $\boldsymbol{\theta}_2$

By setting the first partial derivative of $Q(\theta)$ w.r.t $\boldsymbol{\theta}_2$ to zero, we obtained the updates for all parameters as follows:

$$\begin{aligned}
\boldsymbol{m}_k =& \frac{\sum_i \gamma_{ik}\boldsymbol{w}_{ik}}{\sum_i \gamma_{ik}}, \\
V =& \frac{\sum_i \sum_k \gamma_{ik}(\boldsymbol{w}_{ik} - \boldsymbol{m}_k)(\boldsymbol{w}_{ik} - \boldsymbol{m}_k)^\top}{\sum_i \sum_k \gamma_{ik}} + M, \\
L =& \left[\sum_i \sum_k \gamma_{ik}\boldsymbol{x}_i\boldsymbol{w}_{ik}^\top\right]\left[\sum_i \sum_k \gamma_{ik}\boldsymbol{w}_{ik}\boldsymbol{w}_{ik}^\top + nM\right]^{-1}, \\
\lambda_j =& \frac{1}{n}\sum_i \sum_k \gamma_{ik}(\boldsymbol{x}_{ij} - L_j^\top\boldsymbol{w}_{ik})^2 + L_j^\top M L_j.
\end{aligned} \tag{10}$$

##### 2.3.3 Updating $\xi$

Taking the first derivative of $Q(\boldsymbol{\theta})$ w.r.t as the parameter $\xi$ in the Potts model is difficult because of the normalization constant $C(\xi)$, which requires evaluating the probability mass function of the Potts model over all possible configurations of $\boldsymbol{y}$, and it is thus known to be NP hard. We optimized it numerically via a grid search strategy:

$$\xi = \arg \max_{l \in 1,\ldots,R} Q(\boldsymbol{\theta}_1, \boldsymbol{\theta}_2, \xi_l),$$

where the sequence $\xi_1, \ldots, \xi_R$ is a vector of evenly spaced points in the interval $[0, \xi_{\max}]$. We set the upper bound $\xi_{\max}$ in the uniform distribution to be a large number (set to be 2.5 here) representing the other extreme cases in which spatial location information is highly informative and in which the resulting spatial domain boundaries are extremely smooth.
